# A Characterization of Axolotl Digit Regeneration: Conserved Mechanisms, Divergent Patterning, and a Critical Role for Hedgehog Signaling

**DOI:** 10.1101/2025.06.03.657663

**Authors:** Jackson R. Griffiths, Melissa Miller, Timothy J. Duerr, Ashlin E. Owen, James R. Monaghan

## Abstract

Axolotl digits offer an experimentally versatile model for studying complex tissue regeneration. Here, we provide a comprehensive morphological and molecular characterization of digit regeneration, revealing both conserved features and notable divergences from classical limb regeneration. Digit blastemas progress through similar morphological stages, are nerve-dependent, contain key regenerative cell populations, and express many canonical morphogens and mitogens. However, they exhibit minimal expression of the A-P patterning genes *Shh, Fgf8,* and *Grem1*; suggesting distal outgrowth and patterning occur independently of these signals. Joint regenerative fidelity varies significantly across digits and cannot be explained by differences in nerve supply, cell proliferation, or differential expression of any patterning genes assessed in this study. Furthermore, functional experiments reveal Hedgehog signaling is essential for interphalangeal joint regeneration, but activation alone is insufficient to improve fidelity in less robust digits. This system combines experimental accessibility with intrinsic variation in regenerative outcomes, making it an ideal platform to identify critical determinants of successful tissue regeneration and refine models of appendage patterning.

## Introduction

Urodele amphibians (salamanders and newts) can regenerate complex, multi-tissue appendages throughout their lives. Perhaps the most widely studied example is that of limb regeneration. Upon amputation anywhere along the proximo-distal axis of the limb, salamanders fully and robustly reproduce all missing structures. A critical characteristic of salamander tissues is their ability to redeploy gene expression profiles and signaling circuits in response to injury that were first responsible for the initial development of the limb. This recapitulation of development is thought to be triggered, in part, due to the interaction between axially disparate cell types. The limb blastema, the regenerating structure which forms in response to injury, is largely comprised of connective tissue cells which by the earliest stages of blastema formation exhibit expression profiles similar to those found in the developing limb bud(Gerber et al., 2018; Lin et al., 2021; Muneoka et al., 1986). These dedifferentiated connective tissue cells are thought to retain critical positional “memory” with regards to their origin across the 3 major axes: the anteroposterior, dorsoventral, and proximodistal (Duerr et al., 2024; Nacu et al., 2016; Oliveira et al., 2022; Yamamoto et al., 2022). This positional memory is maintained in the spatial organization of connective tissue cells within the blastema and coincides with the induction of patterning genes, leading to morphogenesis of the regenerating tissue. This is well established with regards to the anteroposterior (A-P) axis, wherein posterior fibroblasts express *Sonic hedgehog (Shh)* while distal anterodorsal fibroblasts express *Fibroblast growth factor 8 (Fgf8*)(Lovely et al., 2022; Nacu et al., 2016). Much like in development, the spatially restricted and juxtaposed expression of these factors leads to a positive feedback loop which directs and maintains the distal outgrowth of limb tissue(Laufer et al., 1994; Tanaka, 2016).

While this phenomenon has been demonstrated in blastemas formed from amputations within the stylopod and zeugopod segments, it is unknown whether the same established models apply to regenerates formed within the phalangeal elements. Digit regeneration was first described by Bonnet in 1777, and since then endeavors to characterize regeneration from these distal-most elements in urodeles have been sporadic (Koussoulakos & Kiortsis, 1989; Smith, 1978). More recently, regenerating digits have exhibited great promise as a model owing to their practical advantages for *in vivo* imaging; as their small size and optical transparency make them ideal for live imaging of transgenic reporter lines(Currie et al., 2016a; Riquelme-Guzmán et al., 2022). These advantages prompt the need for a better understanding of what characteristics and molecular mechanisms are conserved between regenerates from these sites and those of more proximal origin. Importantly, the phenomena of digit regeneration prompts a reevaluation of current models used to explain the essential requirements to mount complex tissue regeneration in the limb. Each digit occupies a distinct and segmented position along the anteroposterior axis, with its number and morphology believed to emerge from developmental signals that specify their anterior-posterior identity. This raises the question of what positional identities are held by digit cells and whether the juxtaposition of disparate axial cell types is required for their regeneration. Moreover, if these interactions and their resulting signaling loops are not reinitiated, then by what mechanisms is distal outgrowth and tissue repatterning achieved?

In this study we first sought to generally characterize digit regeneration using histological and transgenic tools. We found that digit blastemas progress through morphological stages which closely resemble those of more proximal blastemas and similarly contain blastema-associated cell populations and signaling domains. We also confirmed that, as in more proximal blastemas, cell cycle progression in digit blastemas is dependent on an intact nerve supply; and surgical disruption of this nerve supply leads to cell cycle arrest in the G1 phase. However, in contrast to their more proximal counterparts, digit blastemas fail to upregulate the key positionally encoded distal outgrowth genes *Shh, Fgf8,* and *Grem1*. When amputation was performed to remove the distal-most synovial joint and phalanx, we found significant inter-digital differences in the frequency with which digits regenerated the lost joint and proper number of skeletal elements. Treatment with the hedgehog antagonist cyclopamine resulted in a complete loss of joint regenerative fidelity across all digits, while treatment with the hedgehog agonist SAG failed to improve the frequency of joint regeneration in less robust digits. Overall, our analysis reveals similarities and notable differences between digit blastemas and proximal limb blastemas; and our hedgehog perturbation studies suggest a necessary but insufficient role for hedgehog signaling in interphalangeal joint regeneration.

## Results

### Amputated digits undergo similar morphological changes as amputated limbs, and experience nerve dependence similar to limb blastemas

Axolotl forelimbs contain 4 digits termed DI through DIV in anterior to posterior order. Digits DI, DII, and DIV are composed of a proximal and distal phalange, while DIII contains 3 phalanges: proximal, intermediate, and distal. To assess the gross morphological features of digit regeneration, we performed hematoxylin, eosin, and alcian blue staining across multiple regenerative timepoints. Regenerating tissues were obtained via amputations through the two terminal phalanges of digits I–IV resulting in eight distinct amputation sites (**Fig.1A**); with uninjured samples serving as baseline reference (**Fig.1B**). Across all digits and amputation planes, wound closure was achieved by 24 hours in most cases(**Fig.1C, Fig.S1**). Extensive histolysis and skeletal resorption was observed in the underlying stump tissue and remaining phalanx cartilage between 5-10 days post amputation (dpa) as evidenced by a decrease in eosin and alcian staining(**Fig.1D-E, Fig.S1**). This was followed by the accumulation of cells beneath a thickened apical epithelial cap, which begins around 10 dpa and is clearly evident by 20 dpa(**Fig.1E-F, Fig.S1**). By 30 dpa, digit blastemas exhibited distal outgrowth of the mesenchymal tissue and the emergence of stacked chondrocytes distal to the amputated skeleton (**Fig.1G, Fig.S1**).

**Figure 1:**
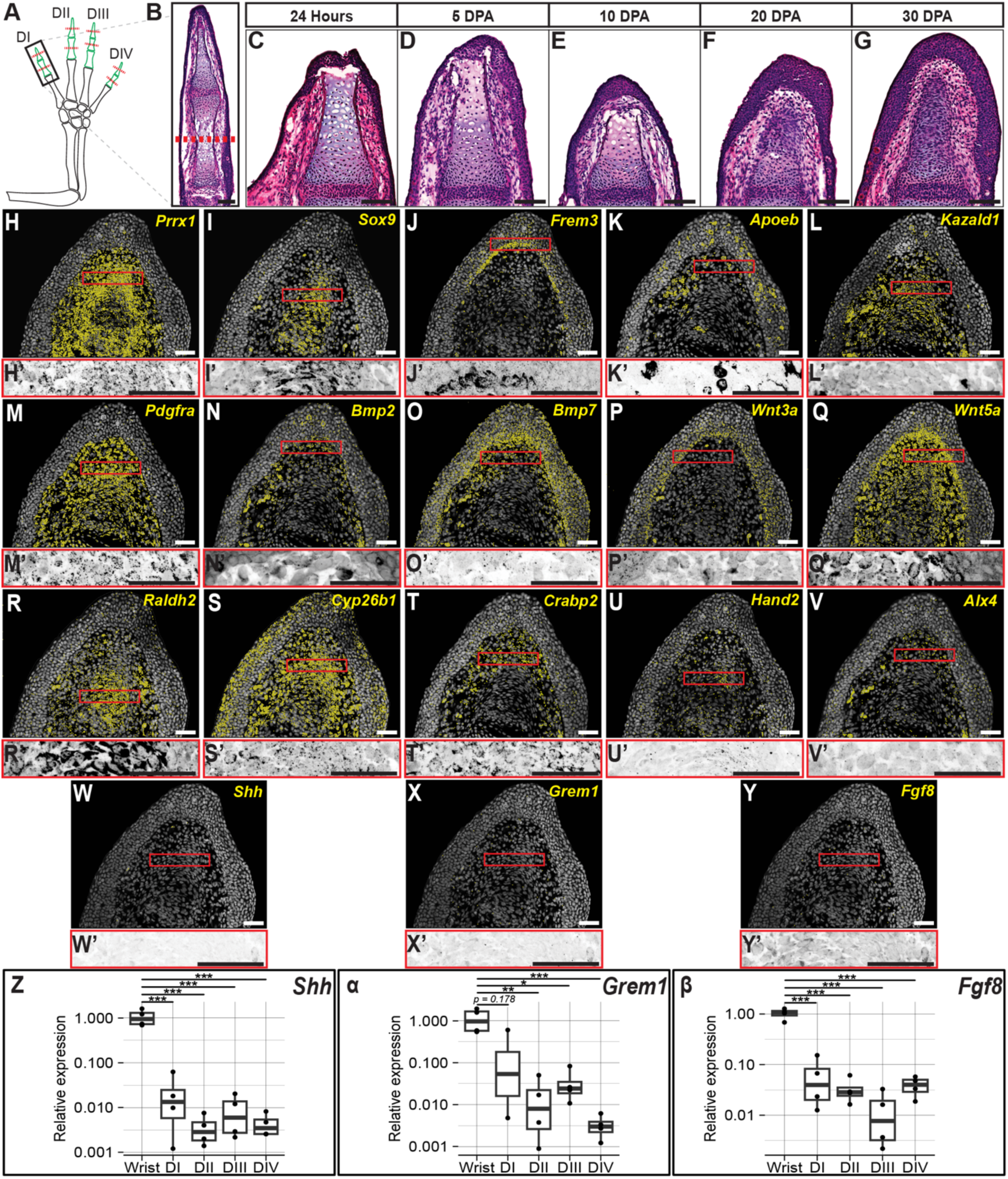
Digit blastema development and marker gene expression. (A) Schematic of amputation planes used to generate histological samples. Phalangeal elements are shown in green, amputation planes are denoted by red dashed lines, and the black box outlines the digit used for representative images in B-Y. (B) Example of an uninjured DI stained with hematoxylin, eosin and alcian (n = 3). Red dashed line denotes amputation plane used to generate tissues in C-Y. (C) DI blastema at 24 hours post amputation exhibiting wound closure (n = 3/3). (D-G) DI blastema at 5, 10, 20, 30 dpa (n = 4, 4, 6, 4). (H-Y) Multiplexed HCR FISH for blastema marker genes and signaling molecules in a 20 dpa DI blastema (n = 3-4 blastemas per gene). FISH signal is displayed as yellow pseudo-dots identified from raw images. Red boxes denote inset location. (H’-Y’) Insets from H-Y showing raw HCR FISH signal. All scale bars = 100 um. (Z-β) qRT-PCR for *Shh*, *Grem1*, and *Fgf8* in 20 dpa wrist and digit blastemas. Four Wrist blastemas and eight digit blastemas were used per replicate with 2-4 replicates per group. Y-axis shows relative gene expression (fold change) plotted on a log_10_ scale. Samples lacking detectable target expression were assigned a Cq value of 40, consistent with the assay’s cycle threshold. Statistical testing was performed via one-way ANOVAs with Tukey-Kramer post hoc tests. All p-values represented by asterisks (* = p < 0.05; ** = p < 0.001; *** = p < 0.001).

To assess whether digit blastemas are composed of cell populations and signaling domains known to play critical roles in regeneration of more proximal limb regions, multiplexed HCR fish was performed for selected targets in tissue sections of 20 and 30 dpa digit blastemas generated from amputations through the proximal (DI, DII, DIV) or intermediate (DIII) phalanges. The *Paired related homeobox 1* (*Prrx1*) transcription factor is expressed by connective tissue fibroblasts during limb development and by dedifferentiated fibroblasts during limb regeneration(Gerber et al., 2018); and we observe *Prrx1* expression throughout the digit blastema mesenchyme by 20dpa (**Fig.1H-H’, Fig.S2**). *Sox9* expressing chondrogenic progenitor cells are also present in the digit blastema by 20dpa (**Fig.1I-I’, Fig.S2**). These cells are positioned adjacent to the existing skeleton in a stacked arrangement and appear to be elongated along the anterior-posterior axis, indicating chondrocyte condensation. A specialized wound epithelium (WE) which directly interfaces with underlying mesenchymal cells is an essential structure within the regenerating limb blastema, and its removal or replacement with full thickness skin prohibits regeneration(Goss, 1956; Mescher, 1976; Thornton, 1957; Tsai et al., 2020). Previous work has shown that *Frem2* serves as a marker of basal epithelial cells of the WE(Leigh et al., 2018), and strong expression of this transcript can be seen in the basal layer of epithelial cells distal to the level of digit amputation (**Fig.1J-J’, Fig.S2**). Extensive myeloid cell recruitment is characteristic of blastema development in more proximal limb injuries and is a requisite for successful regeneration(Godwin et al., 2013). To this effect, we detect *Apolipoprotein eb* (*Apoeb*) expressing myeloid-derived cells throughout the blastema mesenchyme and epithelium in 20 and 30dpa digit blastemas (**Fig.1K-K’, Fig.S2, Fig.S3**). *Kazald1*, a gene specifically enriched in limb blastema cells during regeneration(Bryant et al., 2017) exhibits expression throughout mesenchymal cells of the digit blastema (**Fig.1L-L’, Fig.S2, Fig.S3**). Cell migration is required for blastema formation, and members of the platelet-derived growth factor family have been shown to be essential for promoting connective tissue cell migration during limb and digit regeneration(Currie et al., 2016b). To this effect we see expression of *Platelet-derived growth factor receptor alpha* (*Pdgfra*) throughout the blastema mesenchyme of all digits at 20 dpa (**Fig.1M-M’, Fig.S2, Fig.S3**). Bone morphogenic proteins regulate several processes throughout regenerative stages of proximal blastemas including cell proliferation, skeletal condensation, and apoptosis(Guimond et al., 2010; Sader & Roy, 2022; Vincent et al., 2020). We observe expression of both *Bmp2* and *Bmp7* in 20 and 30dpa digits within the blastema mesenchyme, with *Bmp7* expression also detected in the epithelium (**Fig.1N-N’, Fig.1O-O’, Fig.S2, Fig.S3)**. Wnt ligands of both the canonical/b-catenin and noncanonical Wnt pathways have been shown to play important roles in the initiation and maintenance of limb bud development and regeneration. As in proximal blastemas, we see a similar localization of the canonical Wnt ligand *Wnt3a* in the basal and intermediate epithelium(**Fig.1P-P’, Fig.S2**) and the noncanonical Wnt ligand *Wnt5a* in the distal mesenchyme and basal epithelium (**Fig.1Q-Q’, Fig.S2**)(Glotzer et al., 2022; Lovely et al., 2022). The small molecule retinoic acid acts as a morphogen specifying proximal-distal identity during limb development and regeneration. Recent work has shown that a P-D gradient of retinoic acid signaling is achieved during regeneration through increased degradation of retinoic acid by CYP26B1 in progressively more distal blastema tissue(Duerr et al., 2024). HCR-FISH for genes involved in retinoic acid synthesis (*Raldh2*), transport (*Crabp2*), and breakdown (*Cyp26b1*) reveal that these components are all expressed in the regenerating digit (**Fig.1R-R’, Fig1S-S’, Fig.1T-T’, Fig.S2**). Notably, strong expression of *Cyp26b1* is observed throughout the digit blastema mesenchyme, suggesting active retinoic acid breakdown. To assess A-P identity within the digit blastema, we examined expression of the transcription factors *Hand2* and *Alx4*, which are classically restricted to posterior and anterior mesenchyme, respectively. HCR-FISH reveals that both genes are expressed across all digits but without detectable spatial restriction (**Fig.1U-U’, Fig.1V-V’, Fig.S2**). This uniform expression pattern suggests that the positional asymmetry typically required for A-P patterning may not be well established in the regenerating digit. Building on this observation, the signaling loop established between *Shh* expressing posterior cells, *Fgf8* expressing distal antero-dorsal cells, and *Grem1* expressing cells nested between these domains is critical for distal outgrowth and A-P patterning in both limb development and regeneration(Han et al., 2001; Nacu et al., 2016; Tickle & Towers, 2017; Torok et al., 1999). Notably, expression of these genes was not detected across any digits (DI-DIV) at 20 or 30 dpa (**Fig.1W-Y, Fig.1W’-Y’, Fig.S2, Fig.S3**).

To complement our *in situ* hybridization findings and provide a quantitative comparison of gene expression we performed qRT-PCR across 20dpa digit and wrist blastemas. Consistent with the lack of detectable *in situ* expression, *Shh* and *Fgf8* transcript abundances were markedly reduced across all digits, with fold changes often exceeding a 100-fold decrease relative to wrist blastemas (**Fig.1Z-β**). While *Grem1* expression was also significantly lower in digits DII–DIV compared to wrist, no significant difference was observed in DI (p = 0.178), suggesting partial or variable downregulation in regenerates of this digit. To assess broader signaling dynamics, qPCR was also performed for genes associated with limb patterning, morphogen signaling, and positional identity (**Fig.S4**). No significant differences in expression were detected across digit and wrist blastemas for *Alx1*, *Alx4*, *Bmp2*, *Bmp7*, *Cyp26b1*, or *Ptch1*(**Fig.S4A-E, K**); indicating that these genes are maintained at relatively consistent levels across these amputation sites. Notably, expression of *Pdgfra* remained consistent across digit and wrist tissue, suggesting digit blastemas are composed of a comparable population of migratory blastema cells and that reduced morphogen expression is not simply a reflection of reduced blastema cell composition (**Fig.S4J**). The hedgehog signaling effectors *Gli1*, *Gli2*, and *Gli3* as well as the posterior identity gene *Hand2* all displayed significantly decreased expression in at least one digit compared to wrist (**Fig.S4F-I**). All four genes were significantly less expressed in digits DII and DIV, while *Hand2* and *Gli2* also exhibited decreased expression in DIII. The reduction of these signals in the posterior digits is interesting given the role of Gli transcription factors in responding to posterior Shh and the role of Hand2 in posterior positional memory (Otsuki et al., 2025). Lastly, among all the patterning genes tested, *Gli1* was the only gene assayed which exhibited any significant differential expression between digits.

Overall, these histological and transcriptomic data suggest a similar progression of wound healing and blastema maturation as seen after amputation through more proximal limb regions. However, the absence or reduced expression of A-P patterning genes, particularly *Shh, Fgf8,* and *Grem1,* would suggest that digit blastemas do not contain cells with the full range of A-P positional identities or alternatively do not upregulate these signals in response to injury as observed in more proximal regenerates.

The limb blastema is a densely innervated structure and denervation of the limb’s peripheral nerves via transection at the brachial plexus results in a halt to blastema cell proliferation; as evidenced by a decrease in regenerate size(Singer & Craven, 1948), DNA replication(Duerr et al., 2020; Geraudie & Singer, 1978; Maden, 1978), and mitotic events(Singer, 1952). Previous work by our group led to the generation of a FUCCI (fluorescent ubiquitination-based cell-cycle indicator) axolotl line which provides a live readout of cell cycle state(Duerr et al., 2022) (**Fig.2A**). Using this line, Duerr et al. determined a potential cell-cycle arrest of proximal limb blastema cells in the G1-phase following denervation. To assess whether digit blastemas exhibit a similar response to denervation, bilateral amputations were performed through the mid-diaphysis of the intermediate phalange of DIII in FUCCI axolotls. At 6 dpa right limbs were denervated (n=4 blastemas) while left limbs received a sham surgery (n=4 blastemas). We employed 2-photon microscopy to longitudinally image FUCCI digit blastema volumes immediately preceding denervation (6dpa) followed by reimaging every three days: 9, 12, 15, 18, 21dpa (limbs were re-denervated at 10 days post denervation to circumvent tissue reinnervation) (**Fig.2B**). As epithelial cells in all samples were largely in G1 phase, these cell layers were excluded from the population statistics, similar to previous literature (Geraudie & Singer, 1978). Both innervated and denervated digit blastemas underwent significant histolysis between 6-15dpa (**Fig.2C**). For this reason, the amputation plane could not serve as a reliable marker; therefore, the proximal joint space was used as a reference, and population statistics were conducted on all mesenchymal cells distal to this point. As mentioned, both innervated and denervated blastemas experienced a decrease in total cell number from 6-15dpa (**Fig.2D**); however, while deneverated blastemas continued to decline linearly with time, innervated blastemas experienced a rebound in cell number beginning around 18dpa. In part, the continuous decline in cell number in denervated blastemas could be explained by the significant decrease in the proportion of cycling cells observed by 3 days post denervation and continuing until the 21dpa time point (**Fig.2E**). In comparison, the proportion of cycling cells did not change significantly across time in innervated blastemas (**Fig.2E**). Thus, following the termination of histolysis, innervated blastemas exhibit a rebound in total cell number, while denervated blastemas undergo continuous degeneration (**Fig.2D**).

**Figure 2:**
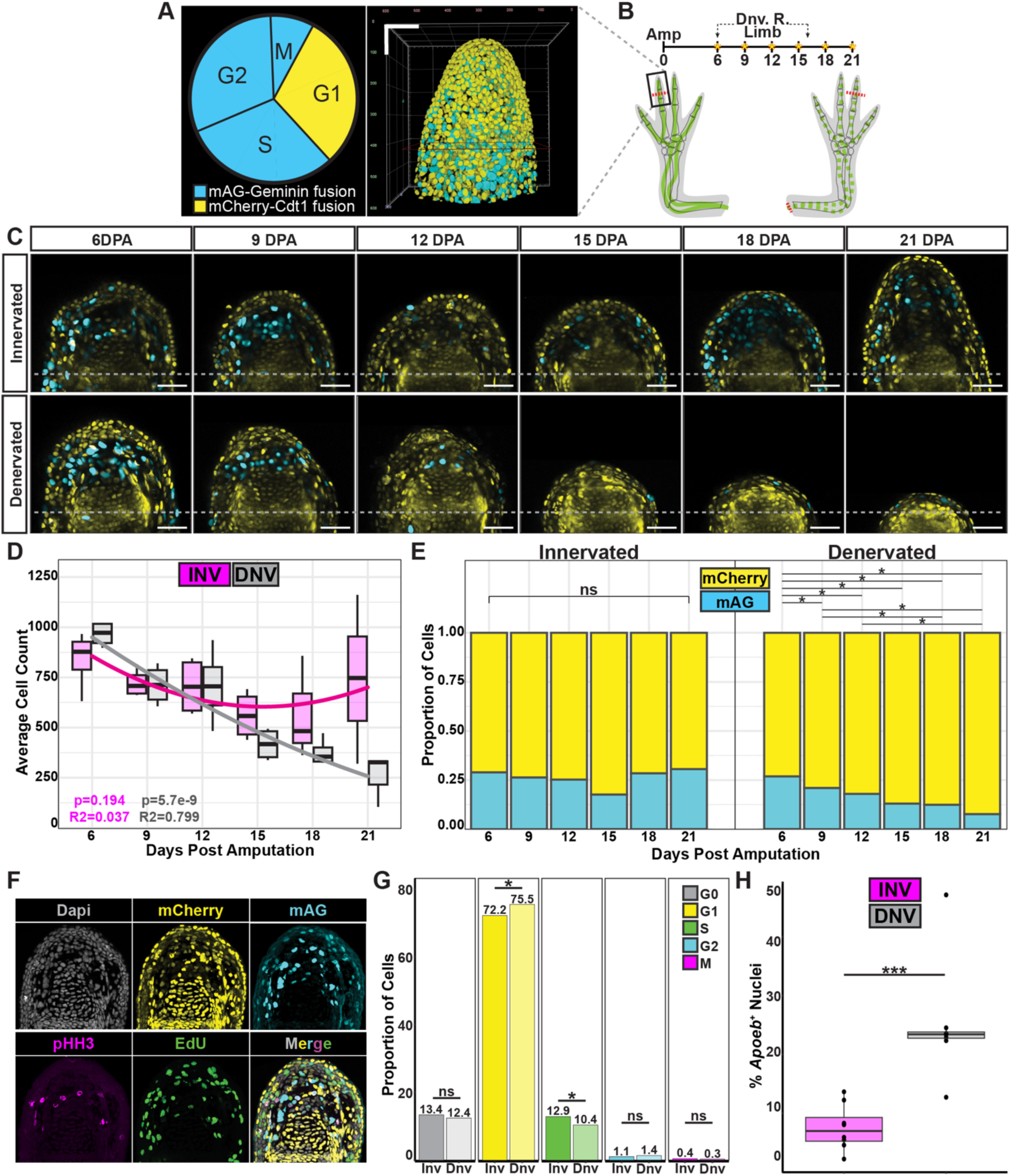
Digit blastema proliferation is nerve-dependent and denervation leads to G1 cell-cycle arrest. (A) Cell cycle schematic showing expression of the FUCCI construct accompanied by a volumetric image of a FUCCI digit blastema. (B) Experimental timeline used to generate *in vivo* time course. Red dashed lines denote amputation planes and nerve transections, yellow asterisks represent image acquisitions, and dashed black arrows represent denervation timepoints. Solid or dashed green lines represent intact or transected nerve, respectively. (C) Representative 2D image planes from an innervated and denervated FUCCI digit blastema across time. Gray dashed line denotes proximal joint used for reference. (D) Average total cell number of innervated and denervated digit blastemas as a function of time (n = 3-4 blastemas per group). Polynomial lines of regression are shown while R^2^ and p-values from linear regression analysis are displayed. (E) Proportion of mAG^+^ and mCherry^+^ cells across time in denervated and innervated digit blastemas (n = 3-4 blastemas per group). Kruskall-Wallis ranked sum analysis was used within groups, followed by one-sided pairwise Wilcoxon tests with FDR correction. (F) Representative images of fluorescent staining patterns in cell cycle pinpoint data. (G) Proportion of cells in each cell cycle phase in innervated and denervated digit blastemas (n = 8 blastemas per group). Chi-squared analyses were used to identify differences in proportion of cells in each phase between groups. Reveals an increase in G1 cells and a decrease in S-phase cells in denervated blastemas. (H) Percentage of *Apoeb*^+^ nuclei in innervated and denervated digit blastemas (n = 8 blastemas per group), showing a significant increase in myeloid derived cells after denervation. Statistical analysis was performed via Wilcoxon ranked sum test with FDR correction. All p-values represented by asterisks (* = p < 0.05; *** p < 0.001). All scale bars = 100 um.

To more accurately resolve the impact of denervation on cell cycle progression, siblings of the animals used for the *in vivo* time course underwent identical procedures, followed by pulse-chase injection of ethynyldeoxyuridine (EdU) for 4 hours at 15 dpa at the mid-blastema stage (n= 8 blastemas per group). Tissue sections were immunohistochemically labelled for phospho-histone H3 (pHH3), imaged, photobleached, click chemistry stained for EdU, and then reimaged (**Fig.2F**). Comparing the proportion of cells in each phase of the cell cycle revealed that denervation leads to a significant decrease in the proportion of cells in S-phase as well as an increase in the proportion of G1 cells (**Fig.2G**), agreeing with previous literature (Duerr et al., 2022; Maden, 1978). These data could suggest that denervation may hinder G1 to S-phase transition, leading to a halt in G1. Moreover, the longitudinal data reveals a continuous decline in cell number following denervation, but the mechanism by which cells are removed after cell cycle arrest are unclear. To that effect, HCR-FISH for the myeloid lineage marker *Apoeb* reveals a significant increase in the number of myeloid derived cells within denervated blastemas (**Fig.2H**), which could suggest heightened activity of immune cells.

### Digit regenerative fidelity varies across the antero-posterior axis

In a pilot cohort of digits amputated through the proximal (DI, DII, and DIV) or intermediate phalanges (DIII) we noted a frequent failure to regenerate the missing joint and distal phalanx. To assess the regenerative fidelity of digit blastemas at these amputation planes animals were amputated bilaterally through the wrist (left limbs) or digits (right limbs) and were allowed to regenerate for three months before alcian blue/alizarin red skeletal staining. Left wrists were saved at the time of amputation and served as uninjured controls. 100 percent of uninjured control digits exhibited the appropriate number of phalanges and joints (**Fig.3A,B**). After wrist amputation, appropriate phalanx and joint numbers were restored with high fidelity and no significant differences were observed between digits (DI: 23/25; DII: 25/25; DIII: 25/25; DIV: 22/25) (**Fig.3A,B**). In contrast, amputation through the digits yielded striking variability in fidelity. DII reproduced the missing joint and phalanx in 100 percent of cases (25/25); while the remaining digits proved to be less robust with DI, DIII, and DIV regenerating the appropriate structures in 56 (14/25), 8 (2/25), and 24 (6/25) percent of cases, respectively (**Fig.3A,B**).

**Figure 3:**
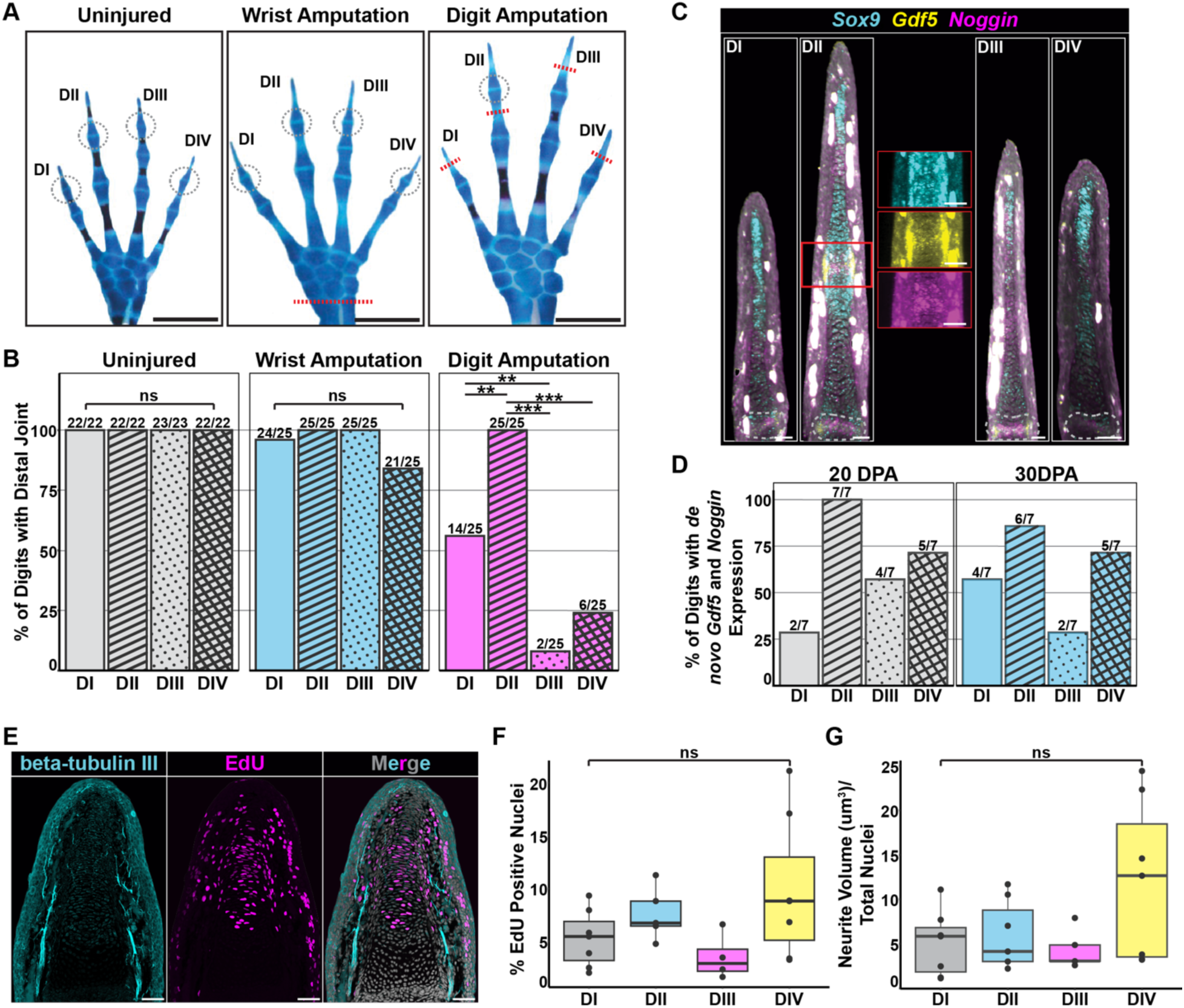
The frequency of joint regeneration and expression of joint morphogenesis genes varies inter-digitally, but there are no differences in proliferation or innervation. (A) Brightfield images of uninjured, wrist amputated, and digit amputated limbs 3 months post amputation. Note the lack of distal joint reproduction after digit amputation of DI, DIII, and DIV. Grey circles outline the distal joint and red dashed lines denote amputation planes. Scale bars = 10 mm. (B) Percent of digits with the distal most joint in uninjured, wrist amputated, and digit amputated limbs (n = 22-23, 25, and 25 respectively). Statistical analysis was performed using chi-squared tests and Holm-Bonferroni corrections (** = p < 0.01; *** p < 0.001). (C) Whole-Mount HCR FISH for *Sox9*, *Gdf5*, and *Noggin* in 20 dpa digit blastemas. Red box denotes inset region and dashed gray outlines indicate joints proximal to the amputation plane. Scale bars = 100 um. (D) Percent of digits with *de novo* expression of *Gdf5* and *Noggin* distal to the amputation plane (n = 7 per digit per time point). (E) Representative images of Beta-3-tubulin staining (combined *Mmu*.*Btub:* memGFP and Immunofluorescence) and EdU click-chemistry stain in a 21 dpa DII blastema. Scale bars = 100 um. (F) Percent of EdU^+^ nuclei in 21 dpa DI, DII, DIII, and DIV blastemas (n = 7, 5, 4, 7). Statistical analysis performed via the Kruskal-Wallis ranked sum test with post-hoc Wilcoxon ranked sum tests using FDR corrections. (G) Total neurite volume um^3^/ total nuclei in 21 dpa DI, DII, DIII, and DIV blastemas (n = 7, 7, 5, 7). Statistical analysis performed as in (F).

To identify whether this failure to restore the distal joint correlated with a failure to redeploy joint morphogenesis genes, we collected digit blastemas at 20 and 30 dpa and performed whole-mount HCR-FISH for *Sox9*, *Gdf5*, and *Noggin*(**Fig.3C**). A precise balance between the expression of *Gdf5* (BMP family member) and *Noggin* (BMP antagonist) is required for appropriate chondrocyte differentiation, cavitation, and interzone formation in synovial joints. When assaying digits for the expression of these factors distal to the amputation plane, a similar trend was seen as in the gross morphological analysis. More specifically, DII appeared to express these genes in a localized region distal to the amputation plane in nearly 100 percent of cases (20dpa: 7/7; 30dpa: 6/7), presumably at the site of de novo joint formation(**Fig.3D**). In contrast, digits DI, DIII, and DIV exhibited this characteristic expression at varying frequencies (DI 20dpa: 2/7; DI 30dpa: 4/7; DIII 20dpa: 4/7; DIII 30dpa: 2/7; DIV 20dpa: 5/7; DIV 30dpa: 5/7) (**Fig.3D**). Together, these data would suggest that digits vary in their ability to appropriately upregulate these joint morphogenesis genes in response to injury.

We next sought to determine whether this variability in the frequency of joint regeneration might coincide with interdigital differences in proliferation or innervation volume. To address this, we performed amputations through the proximal (DI, DII, and DIV) or intermediate phalanges (DIII) of *Beta-3-tubulin* reporter axolotls (Mmu.*Btub:* memGFP); followed by a four-hour pulse of EdU before tissue collection at 21 dpa (**Fig.3E**). Nuclei and neurites were segmented in 3D to quantify total neurite volume (normalized to total cell number) and the percent of EdU positive nuclei. Notably, no significant differences were found between any digits with regards to innervation volume or the percent of proliferating cells (**Fig.3F,G**), suggesting these factors are unlikely to contribute to the observed differences in joint regenerative fidelity.

### Hedgehog signaling is necessary but not sufficient for interphalangeal joint regeneration

The hedgehog family members *Indian hedgehog* (*Ihh*) and *Sonic hedgehog* (*Shh*) both play essential roles in limb morphogenesis. During axolotl limb development and regeneration pharmacological inhibition of hedgehog signaling with cyclopamine treatment leads to a dose-dependent reduction in A-P pattern, resulting in digit loss in a posterior to anterior progression (Purushothaman et al., 2022; Roy & Gardiner, 2002). Similarly, *Shh*^-/-^ mice develop limbs which lack proper A-P pattern in the zeugopod or autopod regions, forming single skeletal elements of anterior-only character(Chiang et al., 2001). In contrast, *Ihh*^-/-^ mice develop limbs with appropriate A-P pattern but fail to form interphalangeal joints, instead forming uninterrupted cartilaginous elements for each digit(Koyama et al., 2007). Notably, joint interzones do form between carpal and zeugopodial skeletal elements, suggesting a direct role of *Ihh* signaling in interphalangeal joint development. Pharmacological activation of hedgehog signaling has been shown to be permissive for limb morphogenesis in injuries lacking the required juxtaposition of axially disparate cell types, as treatment with the smoothened agonist SAG is sufficient to drive full accessory limb development in accessory limb blastemas lacking posterior cells(Nacu et al., 2016). Together these and other data suggest a role for *Shh* in A-P pattern specification, cell proliferation, and survival (Zhu et al., 2022); while *Ihh* regulates chondrocyte differentiation and proliferation as well as interphalangeal joint formation(Koyama et al., 2007; St-Jacques et al., 1999).

We found little to no transcripts of *Shh* could be detected in digit blastemas of DI-DIV at 20 or 30 dpa**(Fig.1O-Q’, Fig.S2, Fig.S3)**. In contrast, strong *Ihh* expression can be seen at both time points concentrated within the *Sox9*-expressing region of chondrogenic progenitors (**Fig.4A-B’, Fig.S2, Fig.S3**). The hedgehog receptor *Patched-1* (*Ptch1*) is expressed at these time points and exhibits localized expression within the regenerating skeleton (**Fig.4C-C’, Fig.S2, Fig.S3**). In contrast the GPCR-like transmembrane hedgehog transducer *Smoothened* (*Smo*) and the hedgehog effector *Gli3* are expressed throughout the blastema mesenchyme (**Fig.4D-E’, Fig.S2, Fig.S3**).

**Figure 4:**
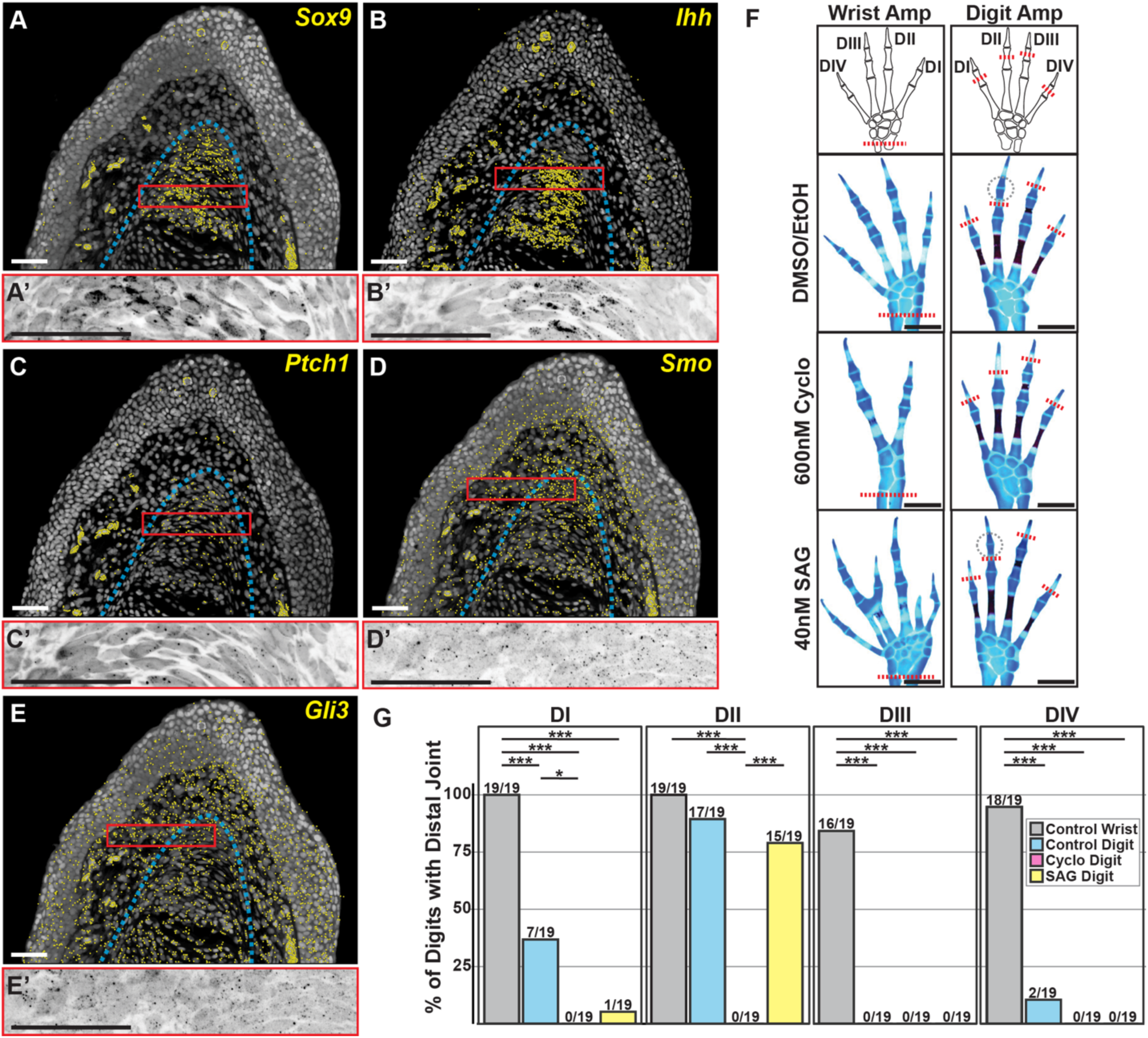
Hedgehog signaling is necessary for interphalangeal joint regeneration but activation of hedgehog signaling does not improve joint regeneration frequency. (A-E) Multiplexed HCR FISH for *Sox9* and Hedgehog signaling genes in a 20 dpa DI blastema (n = 2-3 blastemas per digit). FISH signal is displayed as yellow pseudo-dots identified from raw images. Dashed blue line denotes skeletal boundary and red box denotes inset location of A’-E’. Scale bars = 100 um. (A’-E’) Insets from A-E showing raw HCR FISH signal. Scale bars = 100 um. (F) Brightfield images of wrist or digit amputated limbs 3 months after amputation. Animals were treated for the first 6 weeks of regeneration with: control DMSO/EtOH; 600nM cyclopamine, or 40nM SAG. Red dashed lines denote amputation planes and gray dashed circles outline the distal joint. (G) Percent of digits which regenerated the distal most joint after control treated wrist amputation, control treated digit amputation, cyclopamine treated digit amputation, and SAG treated digit amputation (n= 19 animals per group). Statistical analysis was performed across groups using chi-squared tests followed by pairwise z-tests of proportions with Holm-Bonferroni corrections (* = p < 0.05; *** p < 0.001).

We sought to test whether hedgehog signaling is essential for joint regeneration after digit amputation and whether supplemental hedgehog activation could promote joint formation; either by providing necessary positionally disparate signaling in digits lacking the adequate discontinuities or by directly inducing interphalangeal joint morphogenesis. Animals were amputated bilaterally through the wrist (left limbs) or digits (right limbs) and were randomly separated into 3 treatment groups (40nm SAG, 600nm cyclopamine, control DMSO/EtOH). Animals were treated for 6 weeks, and then allowed to regenerate for 3 months. Left wrist amputations were used to confirm the effectiveness of drug treatments. To that effect, 100 percent of cyclopamine treated wrist amputations exhibited fewer carpals and only 2 digits compared to normal limb morphology, confirming a reduction in A-P patterning (**Fig.4F**). In contrast, SAG treated wrist amputations exhibited an expansion of A-P patterning with a variety of abnormalities. 100 percent of SAG treated wrist amputations exhibited carpal additions/fusions, 52.6 percent exhibited at least 1 additional digit (10/19), and 26.3 percent exhibited the addition of 2 digits (5/19) (**Fig.4F**). Strikingly, 0 percent of cyclopamine treated digit amputations across all four digits regenerated the amputated joint and appropriate number of skeletal elements (0/19 for all digits DI-DIV)(**Fig.4F,G**). In contrast, SAG treatment of regenerating digits had more nuanced effects; with the joint and distal phalanx returning in 5.3 percent of DI amputations (1/19), 78.9 percent of DII amputations (15/19), 0 percent of DIII amputations (0/19), and 0 percent of DIV amputations (0/19) (**Fig.4F,G**). Overall, these data suggest that hedgehog signaling is essential for the regeneration of interphalangeal joints following digit amputation, but activation of hedgehog signaling alone is not sufficient to improve regenerative fidelity in less robust digits (DI, DIII, DIV).

## Discussion

Despite early descriptions of digit regeneration dating back to the origins of research into regenerative phenomena (Bonnet, 1777), our understanding of the mechanisms which govern this process remain inadequate. Here we report that digit blastemas exhibit strong parallels to proximally derived limb blastemas: progressing through comparable morphological stages; containing similar cell populations; expressing many of the same mitogenic and morphogenic signals; and depending upon adequate innervation to sustain proliferation. These parallels reinforce the digit as a relevant and tractable model for investigating limb regeneration. Beyond this, digits offer several practical advantages compared to more proximal regenerates. As we and others have demonstrated, their small size and relative optical transparency make them highly compatible with *in vivo* imaging(Currie et al., 2016b; Riquelme-Guzmán et al., 2022). Experimentally, the presence of multiple digits per limb facilitates increased replicate numbers, and the lack of musculature and motor innervation offers a system with reduced tissue complexity. Finally, the variability in regenerative fidelity between digits, which ranges from near 100% to near 0%, offers a unique opportunity to interrogate what elements contribute to regenerative failure and success in analogous tissues derived from the same genetic background.

While digit blastemas share many features with proximally derived regenerative tissues, they also display striking differences. Most notably, an absence or significant reduction of the positionally encoded A-P patterning genes *Shh, Grem1, and Fgf8*. The failure to activate this morphogen loop may reflect the limited A-P field encompassed by individual digits, in contrast to the broader positional diversity present in more proximal limb regions. These findings suggest that the juxtaposition of axially opposed cells may not be required to initiate a regenerative program in the digit. Moreover, they indicate that the distal outgrowth of digit tissue can proceed independently of this canonical signaling loop, which is otherwise considered essential in established models of complex limb regeneration. Interestingly, while digits appear to lack signals associated with patterning the A-P axis, they do express signals involved in proximodistal patterning and distal outgrowth. Our *HCR* and qPCR data demonstrate the presence of retinoic acid signaling regulators. In particular, strong *Cyp26b1* expression throughout the digit blastema mesenchyme suggests ongoing retinoic acid degradation and distal identity specification (Duerr et al., 2024). We also observe the expression of canonical and noncanonical Wnt ligands which are essential for proximodistal axis establishment and distal outgrowth. The presence of Wnt and RA pathway components in the absence of A-P signals could support a reduced model for tissue morphogenesis in which distal outgrowth and tissue repatterning is largely driven by P-D regulators, with limited contribution by A-P programs.

The observed variability in the fidelity of distal joint and phalanx regeneration across digits is particularly interesting. While our results confirm that digit blastema cell cycling is nerve-dependent, we detected no significant differences in nerve volume or the proportion of proliferating cells among digits. This would suggest that incomplete regeneration in less robust digits cannot be explained by either a failure to reach the nerve supply permissive for complete regeneration or by a deficiency in the quantity of progenitor cells. It’s possible that differential expression of tissue patterning genes across digits could contribute to the variation in regenerative fidelity. However, our analysis did not reveal any detectable differences in the expression of target genes between robust (DII) and less robust (DI, DIII, DIV) digit blastemas. *Gli1* was the only gene that exhibited differential expression between digits, with higher expression observed in DI compared to DII and DIV. However, given that DI and DIV both exhibit lower regenerative fidelity compared to DII, it is unlikely that *Gli1* alone accounts for the observed variation in outcomes. It is also plausible that variation in the extent of cellular dedifferentiation contributes to interdigital differences in regenerative success. The presence of blastema-associated cell populations, morphogens, and mitogens within the regenerating digits is indicative of a regenerative program being initiated, but the degree to which cells in different digits dedifferentiate could contribute to the frequency of proper morphogenesis. Future studies utilizing unbiased, sequencing-based approaches will be critical to identifying interdigital differences in gene expression and assessing the degree to which cells return to a more limb bud-like state.

While inhibition of hedgehog signaling led to complete ablation of joint regenerative fidelity, suggesting an essential role for hedgehog signaling in interphalangeal joint formation, supplemental activation of hedgehog signaling alone was not sufficient to improve the frequency of joint regeneration in less robust digits. This could suggest that other genes involved in joint morphogenesis are absent in these blastemas and their expression is independent of hedgehog signaling. To this point, whole-mount HCR FISH revealed a frequent absence of localized expression of *Gdf5* or *Noggin* distal to the amputation in digits DI, DIII, and DIV. Interestingly, *Ihh*^-/-^ mice develop digits as single uninterrupted skeletal elements, but *Gdf5* expression is still present at high density around the sites where presumptive joints should develop(Koyama et al., 2007). Thus, hedgehog signals and other joint morphogenesis genes may be independently regulated, and rescue of joint regeneration in deficient digits requires activation of multiple signaling pathways. Additionally, a lack of appropriate BMP inhibition may contribute to the failure to reliably reproduce the missing joint. Our data shows strong expression of *Bmp2* and *Bmp7* but reduced expression of the BMP inhibitor *Grem1* and a lack of joint-localized expression of the BMP antagonist *Noggin*. This lack of BMP signal attenuation could lead to premature condensation of chondrocytes or excessive apoptosis, as was described when BMP overexpression was driven in regenerating proximal limbs(Guimond et al., 2010). Alternatively, previous mathematical modeling work (Márquez-Flórez et al., 2018) proposed that joint patterning in digits can arise from a Turing-type reaction-diffusion system involving BMP and WNT interactions. It is possible that in some digits, the regenerative environment fails to meet the necessary conditions to initiate or sustain this self-organizing pattern, leading to an absence of joint formation despite intact upstream signaling pathways.

## Materials and Methods

### Animal care and procedures

d/d axolotl salamanders (*Ambystoma mexicanum)* were bred at Northeastern University or obtained from the Ambystoma Genetic Stock Center (AGSC). FUCCI transgenic axolotls were bred at Northeastern University and beta-3-tubulin reporter animals (Mmu.*Btub:*memGFP) were obtained from the University of Kentucky (Khattak et al., 2013). Surgical procedures, live imaging, and tissue collections were performed under anesthesia using 0.01% benzocaine in axolotl housing water(Yandulskaya & Monaghan, 2023). All procedures were approved by the Northeastern University Institutional Animal Care and Use Committee. Amputations were performed either through the proximal carpals or through the mid-diaphysis of phalangeal elements.

### Sample preparation for histological staining

Immediately following collection, samples were fixed in 4% paraformaldehyde overnight at 4° C. The next morning samples were washed in 1X PBS three times for 5 mins. Samples were cryoprotected in 30% sucrose until equilibrated (approx. 3 hours at RT). Samples were then mounted in OCT and frozen at -80° C. Tissues were then sectioned at 20uM thickness and baked for 15 minutes at 65° C to adhere sections to the slide. Slides were then washed in 1X PBS for 5 mins to remove excess OCT. From here slides were then used for: hematoxylin, eosin, and alcian staining; HCR FISH; click-chemistry; and IHC staining

### Hemotoxylin, eosin, and alcian blue staining

Uninjured and regenerating digit tissues were collected from d/d axolotls with an average length of 10.75cm. Regenerating tissues were collected at 24hr, 5dpa, 10dpa, 20dpa, and 30dpa. Blastemas were generated from amputations through the distal phalanges; or the proximal (DI, DII, DIV) and intermediate phalanges (DIII). Following slide preparation, tissue sections were post-fixed for an additional 15 mins in 4% PFA at RT to ensure tissue adherence to the slide. Slides were then washed with 1X PBS 3 times for 5 mins. Staining was performed as previously published(Duerr et al., 2024), Briefly, slides were submerged in alcian blue solution (0.0001% w/v alcian blue in 60% ethanol and 40% acetic acid) for 10 mins at RT. Slides were then dehydrated in 100% ethanol for 1 min and then airdried. Slides were then submerged in hematoxylin solution for 7 minutes at RT, followed by repeated submersion in fresh tap water 5, 15, and 15 times. Bluing reagent was then applied for 2 mins followed by submersion in fresh tap water 5 times. Eosin solution was then applied for 3 minutes followed by 10 submersions in clean tap water. Slides were then allowed to airdry and mounted using Permount media and glass coverslips. Slides were imaged with a 20X objective using a Zeiss Axio Scan.Z7 microscope at the Harvard Center for Biological Imaging (HCBI).

### Multiplexed HCR FISH

Slides containing consecutive sections of the 20 and 30dpa proximal (DI, DII, DIV) and intermediate phalange (DIII) tissues described above were used for multiround HCR FISH. Version 3 HCR FISH was performed as previously published (Lovely et al., 2023) with minor alterations. Slide preparation was performed as described above followed by an additional 10 min postfixation of tissue sections in 4% paraformaldehyde at RT. Slides were then washed in 1X PBS 3 times for 5 minutes before being moved to 70% ethanol at 4° C overnight, or until use. Slides were then washed in 1X PBS 3 times for 5 mins before beginning the V3 HCR FISH protocol. After each round of staining, probes were wiped by submerging slides in 80% formamide solution (diluted in RNAse free water) four times for 30 minutes each at 37° C. Slides were then washed in 5X SSCT buffer twice for 15 minutes at RT followed by a 5 min wash is 1X PBS. At this point the next round of HCR FISH could be performed beginning with pre-hybridization. Oligonucleotide probe sequences were designed using the [ProbeGenerator](http://ec2-44-211-232-78.compute-1.amazonaws.com/) web application and a full list of sequences is provided in **Table S1**.

Slides were imaged on a Zeiss LSM 880 confocal microscope using Airyscan Fast acquisition. Images were acquired as tiles taken with a 20X objective with 6-11 optical planes across each 20uM section. After acquisition, tiled images were processed using automatic 3D Airyscan Processing; tile stitching; and maximum intensity projection along the z-axis. FISH dot identification and visualization was performed using the FIJI RS-FISH plugin as described previously (Duerr et al., 2024). Background (non-tissue containing region) was subtracted using automatic background removal in Adobe Photoshop.

### qRT-PCR

Amputations were performed bilaterally through the wrist or digits of 9.5cm d/d axolotls. Tissues were allowed to regenerate for 20 days before fresh freezing in liquid nitrogen. 2-4 biological replicates were used for each amputation site and each replicate consisted of 8 or 4 blastemas for digits and wrist, respectively. RNA was isolated from whole tissue, and qRT-PCR was performed via standard procedures. A total of 1 ng of cDNA was used per reaction and all reactions were run in duplicate. Fold changes in mRNA expression were calculated using the comparative Cq (2^−ΔΔCq) method(Livak & Schmittgen, 2001), with *Ef1α* serving as the endogenous control. Statistical testing was performed on ΔCT values using one-way ANOVAs followed by Tukey-Kramer post hoc tests for individual comparisons. Samples lacking detectable expression were assigned a Cq value of 40, consistent with the assay’s cycle threshold, to represent non-expression in quantitative comparisons. A full list of primer sequences is provided in **Table S2**.

### Two-Photon longitudinal imaging

Bilateral amputations were performed through the intermediate phalange of DIII in 4cm FUCCI axolotls. Denervations were performed in right limbs at 6 dpa by making a surgical window above the brachial plexus and severing the three peripheral nerve bundles. Sham denervations were performed in left limbs by making similar windows but leaving the nerve bundles intact. Redenervations and shams were performed at 16 dpa via identical procedures but targeting the nerve bundles proximal to the initial transection. For image acquisition, animals were anaesthetized in 0.01% benzocaine and positioned in a petri dish on their backs over a liquid bed of 1.5% low melt agarose. Before agarose solidification, the digit to be imaged was positioned such that it was flat against the petri dish and extended maximally away from the trunk of the animal. The animal was then submerged in 0.01% benzocaine for the duration of image acquisition. After each image acquisition the animal was removed from the agarose bed and returned to standard housing conditions until the next timepoint.

Two-photon imaging was performed using a Zeiss LSM 880 NLO upright microscope. Images were taken with a water submersible 20X objective. 89-123 optical planes were imaged in each acquisition with a z-resolution of 2.72 um, equating to a depth of imaging between 242.08-334.56 microns.

### 3D segmentation and quantification

3D volumes of image stacks generated from regenerating FUCCI tissues were rendered using Zeiss Arivis Vision4D software. Images were denoised using a discrete gaussian filter and 3D segmentation of cells in each channel was performed using the “cyto2” Cellpose2 neural network model. Mesenchymal tissue distal to the interzone of the proximal joint was then manually segmented throughout each image volume and the resulting ROI was used to compartmentalize data to obtain mesenchyme specific population statistics. The Kruskall-Wallis non-parametric ranked sum test was used to determine whether there were statistically significant differences in the proportion of cycling cells across timepoints in the denervated or innervated group. Post-Hoc one-sided pairwise Wilcoxon ranked sum tests with FDR p-value corrections were then used to determine which timepoints had a significant decrease in the proportion of cycling cells compared to previous timepoints. Linear regression analysis was used to evaluate the impact of time on total cell number in innervated and denervated samples.

### Cell cycle stage analysis

Siblings of the FUCCI animals used in the longitudinal study underwent identical procedures and at 15 dpa received a 4-hour pulse of EdU before tissue collection. EdU was administered via intraperitoneal injection at a dosage of 8 ug/gram of animal.

Samples were processed as described above and resulting slides were post-fixed for 10 mins in 4% PFA at RT. Slides were then washed in 1X PBS 3 times for 5 mins. Sections were permeabilized by applying 0.2% Triton X-100 (diluted in 1X PBS) for 6 mins at RT. Slides were then washed in 1X PBS 3 times for 5 mins. Blocking buffer (15 ul goat serum in 1 mL of 1X PBS) was then applied for 1 hour at RT. Slides were then incubated with rabbit anti-pHH3 antibodies (Cell Signaling, 9701S) diluted 1:100 in blocking buffer for 3 days at 4° C. Slides were washed in 1X PBS 3 times for 5 mins. Alexa-fluor 647 conjugated goat anti-rabbit antibodies (Invitrogen, A21244) were then diluted 1:200 in 1X PBS and applied to slides for 2 hours at RT. Slides were then washed in 1X PBS for 5mins; incubated with DAPI (2.86 uM) for 5 mins; and washed in 1X PBS 3 times for 5mins. SlowFade gold antifade mounting media was then applied to sections and coverslips were applied. After the first round of imaging sections were photobleached by setting the 488, 594, and 647 lasers to 100% laser power and continuously exposing the tissue sections. Images of a subset of tissue sections were acquired after photobleaching to confirm the loss of fluorescent signal. After photobleaching coverslips were floated off tissue sections with RODI water. Slides were then click chemistry stained with Alexa-fluor 647 conjugated azide as previously described (Duerr et al., 2020).

Imaging and photobleaching were performed with a Zeiss LSM 800 confocal microscope using a 20X objective. Images from the consecutive rounds of imaging were then merged in Arivis Vision4D software using manual landmark registration. Nuclei were segmented by first denoising the 405 channels with a discrete gaussian filter, followed by segmentation using the Cellpose2 neural network model. The mean fluorescent intensity for each channel was then calculated for each nucleus. Mean intensity thresholds were then used to classify each nucleus as positive or negative for each fluorescent protein or stain (488: mAG; 594: mCherry; 647 round 1: pHH3; 647 round 2: EdU). Each nucleus was then assigned a cell cycle stage depending on its combination of fluorescent signatures (a full list of observed fluorescent combinations and corresponding cell states is provided in **Table S3**). Chi-squared analyses were used to compare the proportion of cells in each cell phase between innervated and denervated blastemas.

### Apoeb^+^ cell quantification

Slides containing consecutive sections of the innervated and denervated tissues described in the cell cycle stage analysis were used for this analysis (n= 8 blastemas per group). HCR-FISH for *Apoeb* was performed as described above. Imaging was performed on a Zeiss LSM 880 confocal microscope using Airyscan Fast acquisition. Images were acquired as tiles taken with a 20X objective with 6 optical planes across each 20uM section. Images were processed using automatic 3D Airyscan Processing and maximum intensity projection along the z-axis. Nuclei were segmented in AriVis vision 4D software by first denoising with a discrete gaussian filter, followed by segmentation using the Cellpose2 neural network model. *Apoeb*^+^ nuclei were then identified using mean fluorescent intensity thresholding. Statistical tests were performed via the Wilcoxon ranked sum test with FDR corrections

### Whole-Mount alcian blue and alizarin red skeletal staining

At the time of collection samples were fixed overnight in 4% paraformaldehyde at 4°C. Samples were then washed in 1X PBS 3 times for 5 minutes. Skin and soft tissues were then removed using forceps and surgical scissors. Skeletal staining was performed as previously published(Riquelme-Guzmán & Sandoval-Guzmán, 2023) with minor alterations. Samples were dehydrated in a series of 10 minute washes at 25%, 50%, 75%, and 100% ethanol concentrations. Samples were then moved to alcian blue solution (0.0001% w/v alcian blue in 60% ethanol and 40% acetic acid) and placed on a rocker at room temperature overnight. The next day, samples were rehydrated through a series of 10 minute washes at 75%, 50%, 25% and 0% ethanol concentrations. Samples were then incubated in trypsin solution (1% trypsin in 30% Borax) for 30 minutes on a rocker at RT, followed by a wash in 1% KOH for 30 minutes. Samples were then moved to alizarin red solution (0.0001% alizarin red in 1% KOH) and rocked overnight at RT. The next day, samples were washed twice in 1% KOH followed incubation on a rocker in tissue clearing solution (1%KOH and 20% glycerol) overnight at RT. Samples were then dehydrated in a series of 10 minute washes at 25%, 50%, 75%, and 100% ethanol concentrations. Samples were then moved through a series of 30 minute glycerol washes at 30%, 60%, and 100% glycerol concentrations (diluted in ethanol). Tissues were then stored in 100% glycerol until used for imaging.

### Wrist vs digit amputation fidelity assessment

d/d axolotls with an average length of 9.1cm were amputated through the proximal carpals of their left limbs and the proximal (DI, DII, and DIV) or intermediate phalanges (DIII) of their right limbs. Tissues from wrist amputations were saved to serve as uninjured controls. Limbs were collected 3 months post amputation and alcian blue/alizarin red staining was carried out as described above. Brightfield images of each limb were then taken on a Leica M165 FC dissection microscope. Each digit was then assessed for the presence of the distal most joint and assigned a 1 or 0 value according to whether the joint was reproduced or absent, respectively. Chi-squared tests were then calculated across samples within each group (uninjured, wrist, or digit amputation) to determine if any significant differences were observed. Pairwise z-tests of proportions with Holm-Bonferroni corrections were then conducted to identify significant differences between individual digit pairs. Adobe Photoshop automatic background removal was applied to representative images to aide in visualization.

### Whole-Mount HCR FISH for Sox9, Gdf5, and Noggin

d/d axolotls with an average length of 4.1cm received bilateral amputations through the proximal (DI, DII, and DIV) or intermediate (DIII) phalanges. Digits were allowed to regenerate for 20 or 30 days before collection and fixation overnight in 4% paraformaldehyde at 4°C. Whole-mount version 3 HCR FISH was then performed as previously published (Lovely et al., 2023). After HCR FISH staining, samples were mounted in 1.5% low melt agarose and equilibrated in EasyIndex refractive index matching solution (LifeCanvas Technologies, RI: 1.46) overnight at 4°C. Samples were imaged with a 20X objective in EasyIndex on a Zeiss Lightsheet Z1 microscope. Each digit was then assessed for the expression of *Gdf5* and *Noggin* distal to the amputation plane and assigned a 1 or 0 value according to whether expression was present or absent, respectively. Significance testing was then performed as described in the previous section. Representative images were generated from 20 dpa samples that were resliced and maximum intensity projected to capture a 55.92 um thick region along the longitudinal axis of the regenerating digits. Adobe Photoshop automatic background removal was applied to representative images to aide in visualization.

### Interdigital neurite volume and EdU quantification

*Beta-3-tubulin* reporter animals (Mmu.*Btub:*memGFP) with an average length of 6.4cm received bilateral amputations to all digits through the proximal (DI, DII, and DIV) or intermediate phalanges (DIII). At 21 dpa animals received a 4-hour EdU pulse via intraperitoneal injection, followed by tissue collection. Samples were processed and sectioned as described above. To strengthen the labelling of nervous tissue, sections were immunohistochemically stained for B3TUB protein using the protocol outlined above and the following primary and secondary antibodies: rabbit anti-B3TUB (Invitrogen, PA5-85639, 1:100) and Alexa-fluor 488 conjugated goat anti-rabbit antibodies (Life Technologies, A11034, 1:200). During the secondary antibody incubation an ATTO-488 conjugated alpaca anti-GFP antibody (Proteintech, gba488) was used to additionally increase the signal. Slides were then washed in 1X PBS three times for 5 minutes and click chemistry stained as described above.

Images were taken on a Zeiss LSM 800 confocal microscope with a 20X objective. 7 optical planes were imaged across each 20 um section. Tiled images were stitched in ZEN Blue software with automatic settings. Image stacks were then rendered in Arivis Vision 4D software and cropped and rotated to isolate only the tissue distal to the interzone of the proximal joint. Cell nuclei were segmented and mean intensity of the EdU channel was calculated for each nucleus as described above. Neurites were segmented using the Arivis Threshold Based Reconstructor and resulting traces were converted to segments to calculate total neurite volume. For comparisons neurite volumes were normalized to total nuclei number. Statistical analyses were performed with the Kruskal-Wallis test by ranks and pairwise comparisons were made using the Wilcoxon ranked sum test with FDR corrections.

### Pharmacological perturbation of hedgehog signaling

d/d axolotls with an average length of 5.8 cm were amputated through the proximal carpals of their left limbs and the proximal (DI, DII, and DIV) or intermediate phalanges (DIII) of their right limbs. Immediately following amputation, animals were moved to housing water dosed with either 40 nm SAG, 600nM cyclopamine, or dimethyl sulfoxide (DMSO)/ethanol. Note: cyclopamine is dissolved in ethanol while SAG is dissolved in DMSO, thus all 3 types of housing water were made such that they contained equivalent amounts of ethanol and DMSO. Animals were moved to freshly treated housing water every other day for 6 weeks, before being returned to standard housing conditions. After 3 months of regeneration, tissues were collected and processed for alcian blue/ alizarin red staining as described above. Quantification of the frequency of joint regeneration and statistical measurements were carried out as described above.

## Acknowledgements

The authors thank Guoxin Rong and the Institute for Chemical Imaging of Living Systems (RRID:SCR_022681) at Northeastern University for consultation and instrument support. We also appreciate Alex Lovely and the Harvard Center for Biological Imaging (RRID:SCR_018673) for infrastructure and support. The Ambystoma Genetic Stock Center at the University of Kentucky, supported by the NIH (P40-OD019794), is recognized for providing animals for this study.

