## Supplementary Material for "A Characterization of Axolotl Digit Regeneration: Conserved Mechanisms, Divergent Patterning, and a Critical Role for Hedgehog Signaling"

### Figure S1. Histological time course of digit blastemas

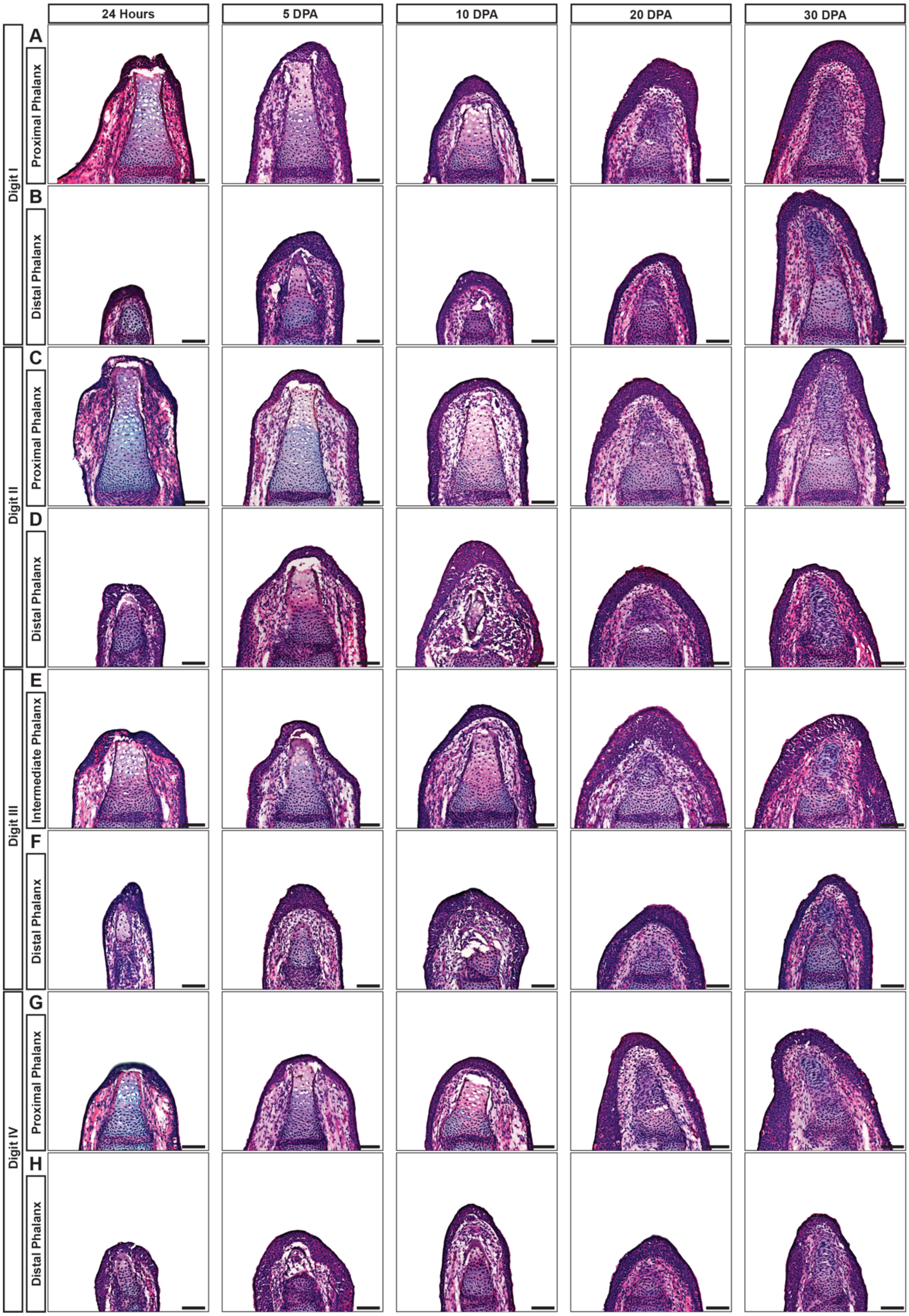

**Figure S1: Histological time course of digit blastemas.** (A) DI proximal phalange blastema at 1, 5, 10, 20, and 30 dpa (n = 3, 4, 4, 6, 4). Wound closure observed by 24 hours in 3/3 cases. (B) DI distal phalange blastema at 1, 5, 10, 20, and 30 dpa (n = 3, 3, 4, 3, 4). Wound closure observed by 24 hours in 3/3 case. (C) DII proximal phalange blastema at 1, 5, 10, 20, and 30 dpa (n = 4, 3, 4, 6, 4). Wound closure achieved by 24 hours in 2/4 cases. (D) DII distal phalange blastema at 1, 5, 10, 20, and 30 dpa (n = 3, 5, 3, 3, 3). Wound closure achieved by 24 hours in 3/3 cases. (E) DIII proximal phalange blastema at 1, 5, 10, 20, and 30 dpa (n = 4, 4, 4, 5, 4). Wound closure achieved by 24 hours in 3/4 cases. (F) DIII distal phalange blastema at 1, 5, 10, 20, and 30 dpa (n = 3, 3, 3, 4, 4). Wound closure achieved by 24 hours in 3/3 cases. (G) DIV proximal phalange blastema at 1, 5, 10, 20, and 30 dpa (n = 4, 4, 4, 4, 3). Wound closure achieved by 24 hours in 4/4 cases (H) DIV distal phalange blastema at 1, 5, 10, 20, and 30 dpa (n = 2, 3, 3, 3, 4). Wound closure achieved by 24 hours in 2/2 cases.

**Figure S2. HCR FISH for blastema marker genes in 20 dpa digit blastemas**

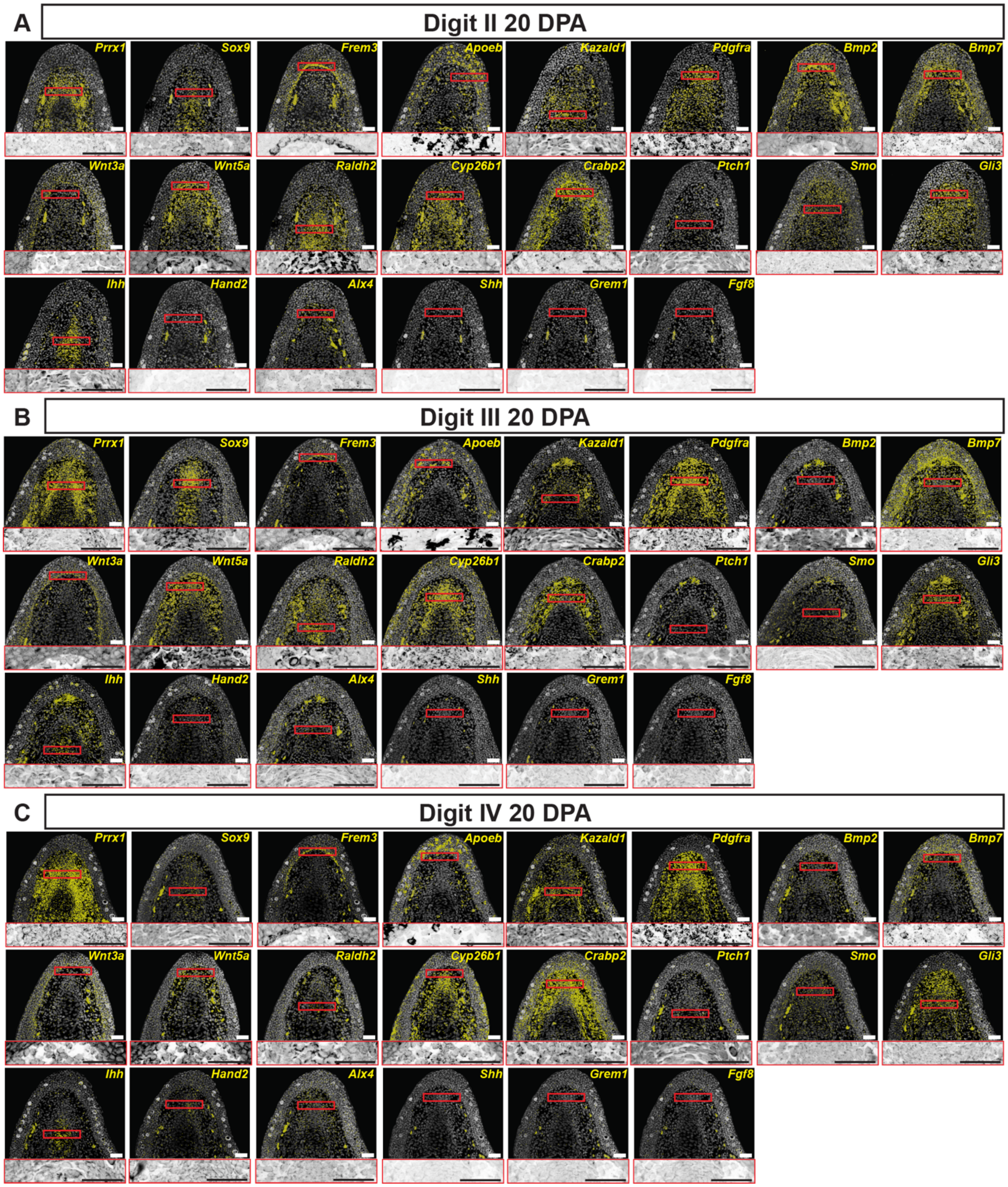


**Figure S2: HCR FISH for blastema marker genes in 20 dpa digit blastemas.** (A) Multiplexed HCR FISH in a DII blastema (n = 2-4 blastemas per gene). (B) Multiplexed

HCR FISH in a DIII blastema (n = 3-4 blastemas per gene). (C) Multiplexed HCR FISH in a DIV blastema (n = 2-5 blastemas per gene). FISH signal is displayed as yellow pseudo-dots identified from raw images. Red boxes denote inset locations. Insets displayed as raw HCR FISH signal. All scale bars = 100 um.

**Figure S3. HCR FISH for blastema marker genes in 30 dpa digit blastemas**

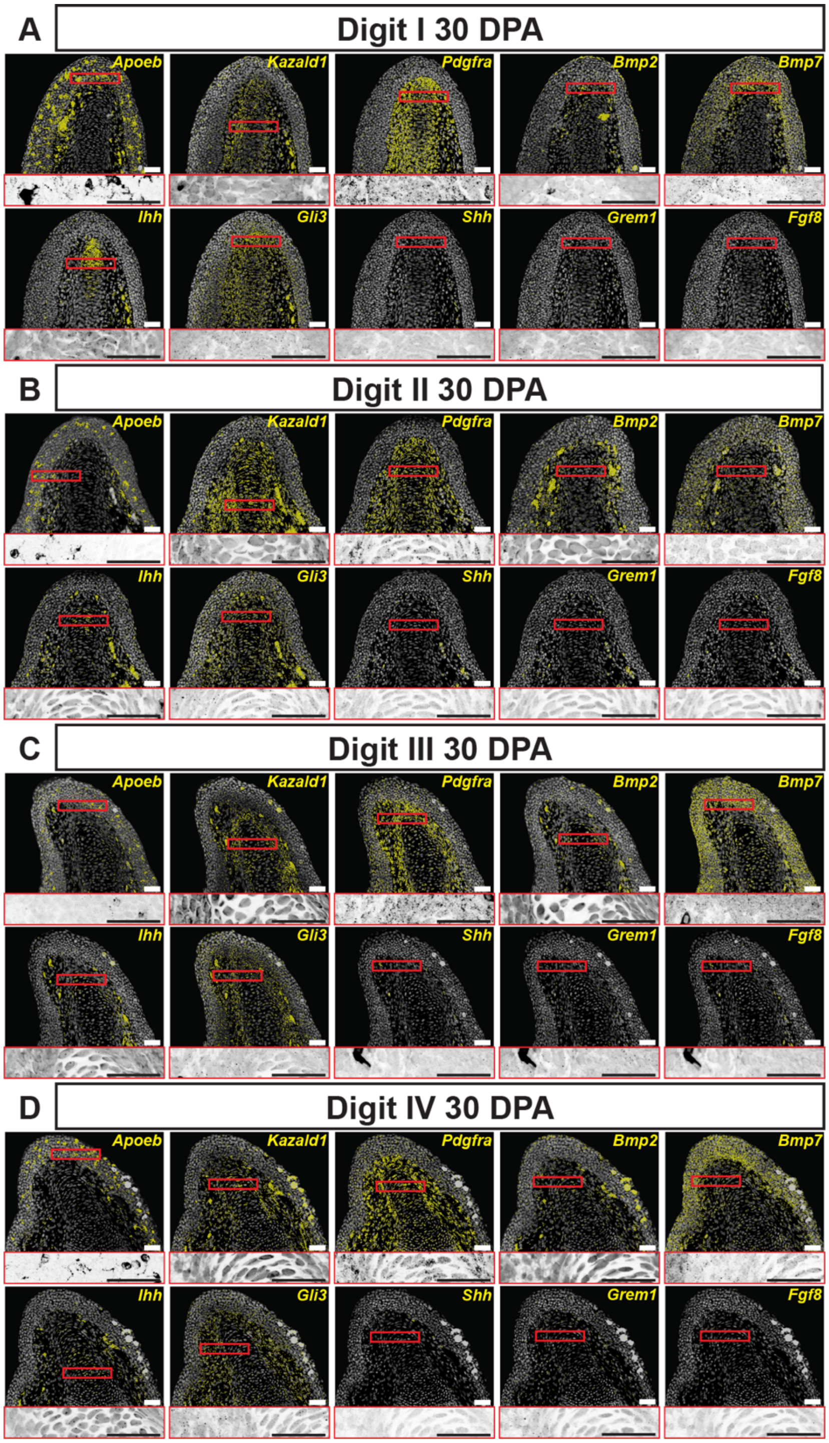


**Figure S3: HCR FISH for blastema marker genes in 30 dpa digit blastemas.** (A) Multiplexed HCR FISH in a DI blastema (n = 3 blastemas per gene). (B) Multiplexed HCR FISH in a DII blastema (n = 2-3 blastemas per gene). (C) Multiplexed HCR FISH in a DIII blastema (n = 3 blastemas per gene). (D) Multiplexed HCR FISH in a DIV blastema (n = 3-4 blastemas per gene). FISH signal is displayed as yellow pseudo-dots identified from raw images. Red boxes denote inset locations. Insets displayed as raw HCR FISH signal. All scale bars = 100 um.

**Figure S4. qRT-PCR of blastema-associated genes across wrist and digit tissues**

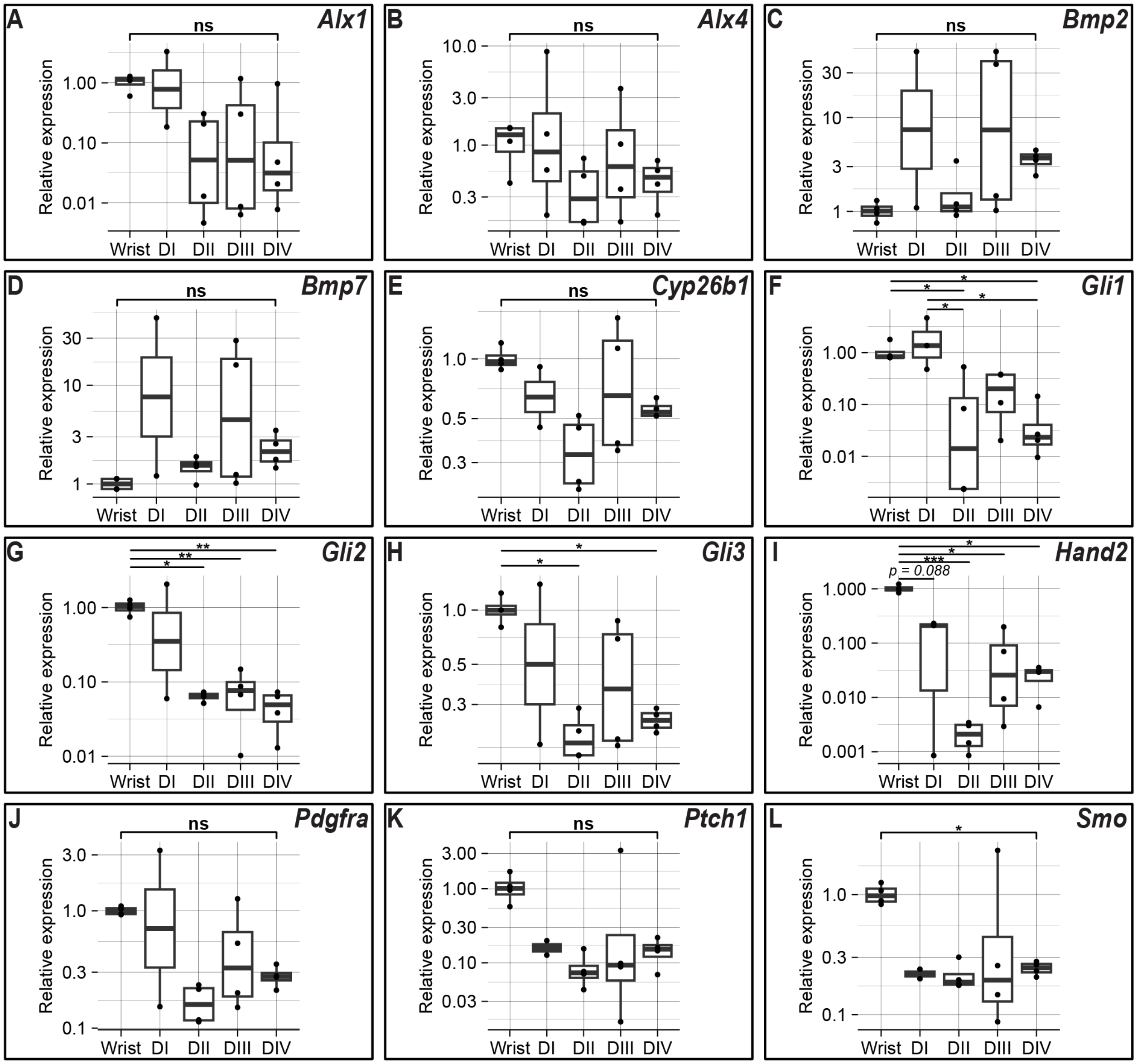


**Figure S4. qRT-PCR of blastema-associated genes across wrist and digit tissues.** (A-L) qRT-PCR for target genes in 20 dpa wrist and digit regenerates. 2-4 biological replicates were used for each amputation group and each replicate consisted of 8 or 4

blastemas for digits and wrist, respectively. Two technical replicates were run for every biological replicate. Fold changes in mRNA expression were calculated using the comparative Cq ($2^{-\Delta\Delta Cq}$) method(Livak & Schmittgen, 2001), with *Ef1α* serving as the endogenous control. Y-axis shows relative gene expression (fold change) plotted on a $\log_{10}$ scale. Statistical testing was performed on ΔCT values using one-way ANOVAs followed by Tukey-Kramer post hoc tests for individual comparisons. All p-values represented by asterisks (* = $p < 0.05$; ** = $p < 0.001$; *** = $p < 0.001$). Note the only gene which shows significantly different expression between digits is *Gli1*.

**Table S1:**

| **Wnt3a** | |
|---|---|
| Wnt3a_B1 | gAggAgggCAgCAAACggAAAGCCCATCTCAGAAGTAGCACAAAA |
| Wnt3a_B1 | CACAAAGAAACAGGGTACATCCAAATAgAAgAgTCTTCCTTTACg |
| Wnt3a_B1 | gAggAgggCAgCAAACggAAGGTAACTTGACAGCACTTGCGCCAG |
| Wnt3a_B1 | GACCAATGGCTAATGACCACCAAATTAgAAgAgTCTTCCTTTACg |
| Wnt3a_B1 | gAggAgggCAgCAAACggAAGCTGTGTGCCCAGGGAAGAGTACTG |
| Wnt3a_B1 | GCCCCGGGATGCTGCCGCACAGGATTAgAAgAgTCTTCCTTTACg |
| Wnt3a_B1 | gAggAgggCAgCAAACggAAGACAGAAGCGCAGCTGCTTGGGCAC |
| Wnt3a_B1 | CACTGGGCATGATCTCCACATAGTTTAgAAgAgTCTTCCTTTACg |
| Wnt3a_B1 | gAggAgggCAgCAAACggAAGAATGCCGATCTTCACGCCTTCCGC |
| Wnt3a_B1 | GGCCCCGGAACTGGTGTTGACACTCTAgAAgAgTCTTCCTTTACg |
| Wnt3a_B1 | gAggAgggCAgCAAACggAACGTTGACTGTCGTGCAGTTCCACCT |
| Wnt3a_B1 | ACACTGGCCCAAAGATAGCCAGGTTTAgAAgAgTCTTCCTTTACg |
| Wnt3a_B1 | gAggAgggCAgCAAACggAAAGGCGGACTCTCTCGTGGCTTTATC |
| Wnt3a_B1 | CTCCAGCAGAGGCGATTGCGTGTACTAgAAgAgTCTTCCTTTACg |
| Wnt3a_B1 | gAggAgggCAgCAAACggAACACAGGAGCGGGTTACTGCAAATGC |
| Wnt3a_B1 | AGCCACATATTGTGGCAGATCCTTCTAgAAgAgTCTTCCTTTACg |
| Wnt3a_B1 | gAggAgggCAgCAAACggAACCCCATTTCCAGCCTTCGCCAGGGG |
| Wnt3a_B1 | CCAAACTCCACATCTTCGCTGCAGCTAgAAgAgTCTTCCTTTACg |
| Wnt3a_B1 | gAggAgggCAgCAAACggAATCGGCAAACTCTCGCGAGACCATGC |
| Wnt3a_B1 | CGAGCATCTGGCCTGTTCTCTCTGGTAgAAgAgTCTTCCTTTACg |
| Wnt3a_B1 | gAggAgggCAgCAAACggAATCATTGTTGTGCCGGTTCATGGCAG |
| Wnt3a_B1 | TGATCAATGATCGACGTTCGTCCAGTAgAAgAgTCTTCCTTTACg |
| Wnt3a_B1 | gAggAgggCAgCAAACggAACCATGACATTTGCATTTCAGGTGCA |
| Wnt3a_B1 | GTCTTCACCTCACAGCTTCCGGATATAgAAgAgTCTTCCTTTACg |
| Wnt3a_B1 | gAggAgggCAgCAAACggAACTGAAGTCTGGCTGGGACCACCAGC |
| Wnt3a_B1 | TTGTCCTTTAGATAGTCGCCAATGATAgAAgAgTCTTCCTTTACg |

| | |
|---|---|
| Wnt3a_B1 | gAggAgggCAgCAAACggAAACGACCATTTCAGAGGCGCTGTCGT |
| Wnt3a_B1 | CAGCCACGGGATTCACGATGCTTCTTAgAAgAgTCTTCCTTTACg |
| Wnt3a_B1 | gAggAgggCAgCAAACggAAGTGTATTTGGGACGTAGGGTCTCAA |
| Wnt3a_B1 | TCTCTCTCAGTTGGTGGCTTGAAGATAgAAgAgTCTTCCTTTACg |
| Wnt3a_B1 | gAggAgggCAgCAAACggAATTTGGCGAGCTCTCGTAGTAAACCA |
| Wnt3a_B1 | CCTGTTTCAGGGTTTGGCTCGCAAATAgAAgAgTCTTCCTTTACg |
| Wnt3a_B1 | gAggAgggCAgCAAACggAACAAATCCTGTCACGGGTGCCAAAGG |
| Wnt3a_B1 | CCATCAATGCCGTGAGAGGTCACATTAgAAgAgTCTTCCTTTACg |
| Wnt3a_B1 | gAggAgggCAgCAAACggAACCACGACCACAGCACAGGAGATCAC |
| Wnt3a_B1 | TTCCTTTTCTCAGTCCGTGTGTTGTTAgAAgAgTCTTCCTTTACg |
| Wnt3a_B1 | gAggAgggCAgCAAACggAACCAGTGGAAGATGCAGTGACATTTT |
| Wnt3a_B1 | GCATTCTTGACAGCTCACGTAGCAATAgAAgAgTCTTCCTTTACg |
| Wnt3a_B1 | gAggAgggCAgCAAACggAAAAGGCTATCATTGACTTCCTCAAAA |
| Wnt3a_B1 | CTTTGTCAGGAAAGTTTTCAGCAAATAgAAgAgTCTTCCTTTACg |
| Wnt3a_B1 | gAggAgggCAgCAAACggAACAACCACAGTGGAAACTGAAGACAT |
| Wnt3a_B1 | ACAGTCATGATCTTAAGAGTTCTGCTAgAAgAgTCTTCCTTTACg |
| Wnt3a_B1 | gAggAgggCAgCAAACggAACGGTTCTGTCTTGGTGCTCGATCAT |
| Wnt3a_B1 | TCCTGGGAGGCTTCAGGATGAAGATTAgAAgAgTCTTCCTTTACg |
| Wnt3a_B1 | gAggAgggCAgCAAACggAAATCCTTTCTCTAGTGATGATAGATA |
| Wnt3a_B1 | ACATGTTGCATGTTGTCCTAAGAGCTAgAAgAgTCTTCCTTTACg |
| Wnt3a_B1 | gAggAgggCAgCAAACggAAGGGATTTTAAAATCTACACTTAAAA |
| Wnt3a_B1 | GTTTAAATGTGCTTTCAAAATGCCATAgAAgAgTCTTCCTTTACg |
| **Wnt5a** | |
| Wnt5a_B2 | CCTCgTAAATCCTCATCAAAATTGGCAAAGGAGGCTGGCGTAGGC |
| Wnt5a_B2 | TGCTCAGTATCCGAGCACGAGCGAGAAATCATCCAgTAAACCgCC |
| Wnt5a_B2 | CCTCgTAAATCCTCATCAAAGTTCATGGCCAGGGACCACCAGGAG |
| Wnt5a_B2 | GATGTATGCCTCCGGGATTTGAACAAAATCATCCAgTAAACCgCC |
| Wnt5a_B2 | CCTCgTAAATCCTCATCAAACAGCTTCTTCTGCCCAGGGGACAGC |
| Wnt5a_B2 | AGGCATGTGATCCTGGTACAGCTGGAAATCATCCAgTAAACCgCC |
| Wnt5a_B2 | CCTCgTAAATCCTCATCAAAACCGGTCTTTGCCCCTTCTCCAATA |
| Wnt5a_B2 | CCGGAACTGGTACTGGCACTCTTTAAAATCATCCAgTAAACCgCC |
| Wnt5a_B2 | CCTCgTAAATCCTCATCAAAGTCCACCGTGCTGCAGTTCCACCTC |
| Wnt5a_B2 | CATAACTCGGCCAAACACAGAGGCGAAATCATCCAgTAAACCgCC |
| Wnt5a_B2 | CCTCgTAAATCCTCATCAAATATTCCAGATTGTCCCCGCAGCCCC |
| Wnt5a_B2 | ACGAACTCTTTGGCAAAGCGGTAGCAAATCATCCAgTAAACCgCC |
| Wnt5a_B2 | CCTCgTAAATCCTCATCAAATGTATTTTCTCTCTCTCTCGGGCAT |
| Wnt5a_B2 | CTGGAGCTTTCGTAGGACCCCTTGGAAATCATCCAgTAAACCgCC |
| Wnt5a_B2 | CCTCgTAAATCCTCATCAAATCGTTGTTGTGTATGTTCATTAATG |
| Wnt5a_B2 | AGATTATACACTGTCCTTCTGCCAGAAATCATCCAgTAAACCgCC |

| | |
|---|---|
| Wnt5a_B2 | CCTCgTAAATCCTCATCAAAGAAACTCCGTGGCATTTGCAGGCGG |
| Wnt5a_B2 | CAGCAGGTCTTCAGGCTGCAGGATCAAATCATCCAgTAAACCgCC |
| Wnt5a_B2 | CCTCgTAAATCCTCATCAAAACCTTGCGGAAGTCAGCCAACTGCA |
| Wnt5a_B2 | TCATACTTCTCCTTCAGGAAGTCCCAAATCATCCAgTAAACCgCC |
| Wnt5a_B2 | CCTCgTAAATCCTCATCAAAGCGTTCAGCCTCATAGAGGCAGCGC |
| Wnt5a_B2 | CTGTTCACCTGCACCAGCTTACCCCAAATCATCCAgTAAACCgCC |
| Wnt5a_B2 | CCTCgTAAATCCTCATCAAAATCGTTCGTGGTGGCGGGTTGAAG |
| Wnt5a_B2 | GTCTGGGCTCGTGTCCACATACACCAAATCATCCAgTAAACCgCC |
| Wnt5a_B2 | CCTCgTAAATCCTCATCAAAGTTGCAGAGTCGGCCCTGAGTTCCC |
| Wnt5a_B2 | GCAGCCATCCATGCCCTCAGATGTCAAATCATCCAgTAAACCgCC |
| Wnt5a_B2 | CCTCgTAAATCCTCATCAAATGGTCCACGATCTCAGTGCACTTTT |
| Wnt5a_B2 | GGGCCGGTGCACTACTTGCACACAAAAATCATCCAgTAAACCgCC |
| Wnt5a_B2 | CCTCgTAAATCCTCATCAAAAGTGGATAAACATTGTCTCTGAGCG |
| Wnt5a_B2 | GCCACACTGCCTGAGCTCCATAAATAAATCATCCAgTAAACCgCC |
| Wnt5a_B2 | CCTCgTAAATCCTCATCAAACGAAGAACAAGTTCCAGTCACTTTT |
| Wnt5a_B2 | TCTGCCGCTGAGCAGTCCTCGGATAAAATCATCCAgTAAACCgCC |
| Wnt5a_B2 | CCTCgTAAATCCTCATCAAATAGTTTGGGTCAGATTCAACGCCCA |
| Wnt5a_B2 | GTCCATAACGTCTTTGTTGTAAATAAAATCATCCAgTAAACCgCC |
| Wnt5a_B2 | CCTCgTAAATCCTCATCAAATAGTTACTTCCCCTTACTCTGCTGT |
| Wnt5a_B2 | CCGTGACCACCCCATGTACTTAGTTAAATCATCCAgTAAACCgCC |
| Wnt5a_B2 | CCTCgTAAATCCTCATCAAACGCCCTGTGCTAGGCAACTCTGTGC |
| Wnt5a_B2 | ACACTGTTAAGCGGAAAGTGCACACAAATCATCCAgTAAACCgCC |
| Wnt5a_B2 | CCTCgTAAATCCTCATCAAACAGAGGAGGCCTATAGGCCTGGATG |
| Wnt5a_B2 | TATTAAAGCAGTAAAATGTGTGTTAAAATCATCCAgTAAACCgCC |
| Wnt5a_B2 | CCTCgTAAATCCTCATCAAATTACACAACTGACCCAATGCATGAA |
| Wnt5a_B2 | GACATCAGTGCACACACACAGTGTCAAATCATCCAgTAAACCgCC |
| Wnt5a_B2 | CCTCgTAAATCCTCATCAAACATTGCTATCATCAAATTGCAATAC |
| Wnt5a_B2 | TCTCACATCCACAAAACAAATACTTAAATCATCCAgTAAACCgCC |
| Wnt5a_B2 | CCTCgTAAATCCTCATCAAATTGGTAAACGCTTGTGTGAACTTTT |
| Wnt5a_B2 | TCAATCTATTAGAGCCCAGGGATGTAAATCATCCAgTAAACCgCC |
| Wnt5a_B2 | CCTCgTAAATCCTCATCAAATAAAGGTCAATACACAGAGTTTGCT |
| Wnt5a_B2 | GAGGGGTCAATTAAACCCCACTAGTAAATCATCCAgTAAACCgCC |
| **Grem1** | |
| Grem1_B2 | CCTCgTAAATCCTCATCAAACAGCTCCACGCGCGGAGAGATGGGG |
| Grem1_B2 | AGCAGAGCAGAGCGCGGGGCGGCTGAAATCATCCAgTAAACCgCC |
| Grem1_B2 | CCTCgTAAATCCTCATCAAAGGCGCTGCAGATGCGCGGGCTCAGT |
| Grem1_B2 | AGCCTAGGCGTGGGCACTCGCGGGGAAATCATCCAgTAAACCgCC |
| Grem1_B2 | CCTCgTAAATCCTCATCAAATGTCAGGCGAAAGGCGGGCTCGTCC |
| Grem1_B2 | GATTCTCTGCTTCTGGGCGCTATGTAAATCATCCAgTAAACCgCC |

| | |
|---|---|
| Grem1_B2 | CCTCgTAAATCCTCATCAAAGTCCCGCGGCTCCCGGGGCCAATC |
| Grem1_B2 | TAGCTCCCCGTGCTCAGTATTCTGGAAATCATCCAgTAAACCgCC |
| Grem1_B2 | CCTCgTAAATCCTCATCAAATCCTGTTGAGTGGGTGGGTGGAACA |
| Grem1_B2 | CGCCAACTGCGTAGACCAATCGAAAAAATCATCCAgTAAACCgCC |
| Grem1_B2 | CCTCgTAAATCCTCATCAAAGCATCAGCCCGTAGAGCAGGAGAAG |
| Grem1_B2 | TTTTCTTTTCTCCCTCTGCCACAGGAAATCATCCAgTAAACCgCC |
| Grem1_B2 | CCTCgTAAATCCTCATCAAAGTGGGATGGCTCCCTGAGACCCTCG |
| Grem1_B2 | AGTCATTGGGCTGTGCCTTGTCGGGAAATCATCCAgTAAACCgCC |
| Grem1_B2 | CCTCgTAAATCCTCATCAAACATGGCTGTGCCTTTTCCTCGCCCC |
| Grem1_B2 | GCTGGACTCCAGCACCTCCTCGGCAAAATCATCCAgTAAACCgCC |
| Grem1_B2 | CCTCgTAAATCCTCATCAAAGCGCTCAGTGACGTGAAGGGCTTCC |
| Grem1_B2 | CTTACACCAGTCCCGCTTAAGGTACAAATCATCCAgTAAACCgCC |
| Grem1_B2 | CCTCgTAAATCCTCATCAAAGTGGATGGTCTGCTTCAGCGGTTGG |
| Grem1_B2 | GATGGTGCGGCTGTTGCAGCCTTCCAAATCATCCAgTAAACCgCC |
| Grem1_B2 | CCTCgTAAATCCTCATCAAAGCACTGGCCGTAGCAGAAGCGGTTC |
| Grem1_B2 | GACATGTCTGGGTATGTAGAAGGAGAAATCATCCAgTAAACCgCC |
| Grem1_B2 | CCTCgTAAATCCTCATCAAAGGACTGGAAGGAGCCTTCCTCCCGG |
| Grem1_B2 | GAACTTCTTGGGCTTGCAGAAGGAGAAATCATCCAgTAAACCgCC |
| Grem1_B2 | CCTCgTAAATCCTCATCAAAGCAATTGAGCGTCACTGTCATGGTG |
| Grem1_B2 | CTTCTTGGTGGGCGGCTGCAGCTCTAAATCATCCAgTAAACCgCC |
| Grem1_B2 | CCTCgTAAATCCTCATCAAAACATTGCTTGACTCTGGTGATTCTT |
| Grem1_B2 | CTACTCCAAGTCAATGGAGATGCAAAAATCATCCAgTAAACCgCC |
| Grem1_B2 | CCTCgTAAATCCTCATCAAAGACCATTGGCCTGTCACTTGTCGAG |
| Grem1_B2 | AAGCTAAACTGTCCTCTGCATTCTGAAATCATCCAgTAAACCgCC |
| Grem1_B2 | CCTCgTAAATCCTCATCAAACTCTTCACCCATTTTCAGAGAGGTG |
| Grem1_B2 | GGGCCGTGATTCAGGCATGTGTGGTAAATCATCCAgTAAACCgCC |
| Grem1_B2 | CCTCgTAAATCCTCATCAAACACTGAAGACTTTACCCTTCCAAGG |
| Grem1_B2 | CCATCAGGGCAGGGGTTATCCATCTAAATCATCCAgTAAACCgCC |
| Grem1_B2 | CCTCgTAAATCCTCATCAAAAAAATTGAACTACAGTTCCCAGACA |
| Grem1_B2 | GACAAGTCACCCTGATTTTCTGTTTAAATCATCCAgTAAACCgCC |
| Grem1_B2 | CCTCgTAAATCCTCATCAAAGGGGTTCATGGTGAGATTATTTCCC |
| Grem1_B2 | AGAGATGTGCAAAAGTTCAGAGATGAAATCATCCAgTAAACCgCC |
| Grem1_B2 | CCTCgTAAATCCTCATCAAACGTAGGCAAGAGGCTCGGGGTGGGG |
| Grem1_B2 | ACCGGGATCTTTCAAGGATGACAAGAAATCATCCAgTAAACCgCC |
| Grem1_B2 | CCTCgTAAATCCTCATCAAAGGTGCAGAAGCAGCGGGAAATCCAT |
| Grem1_B2 | TCCAGCATGTTGCTCAGAATTCAGAAAATCATCCAgTAAACCgCC |
| Grem1_B2 | CCTCgTAAATCCTCATCAAACGTCCTCGTGCGTTGCGTTCTAAGA |
| Grem1_B2 | ATTGGCTGTTGCTATAGCAGAGCGTAAATCATCCAgTAAACCgCC |
| **Shh** | |

| | |
|---|---|
| Shh_B1 | gAggAgggCAgCAAACggAATTATAGGCGAGCGGGGTCAGCTTTT |
| Shh_B1 | TTCTCGGCCACGTTGGGGATGAACTTAgAAgAgTCTTCCTTTACg |
| Shh_B1 | gAggAgggCAgCAAACggAAATCTTGCCTTCGTATCGCCCGCTGG |
| Shh_B1 | TCTTTGAAGCGGTCGGAGTTGCGGGTAgAAgAgTCTTCCTTTACg |
| Shh_B1 | gAggAgggCAgCAAACggAAAATGATGTCCGGGTTGTAATTGGGG |
| Shh_B1 | CGCTCCGGTGTTCTCTTCGTCCTTATAgAAgAgTCTTCCTTTACg |
| Shh_B1 | gAggAgggCAgCAAACggAACTTACACCTCTGGGTCATTAGCCTG |
| Shh_B1 | TGAGATGGCTAGGGCGTTCAGCTTATAgAAgAgTCTTCCTTTACg |
| Shh_B1 | gAggAgggCAgCAAACggAACTTCACCCCAGGCCACTGGTTCATC |
| Shh_B1 | TTCATCCCAGCCTTCAGTCACCCGCTAgAAgAgTCTTCCTTTACg |
| Shh_B1 | gAggAgggCAgCAAACggAACTGAGGTTGTGATGTCCACGGCCCG |
| Shh_B1 | GCATGCCGTATTTGCTTCGGTCTCGTAgAAgAgTCTTCCTTTACg |
| Shh_B1 | gAggAgggCAgCAAACggAAAACCCAATCAAAGCCGGCCTCCACG |
| Shh_B1 | ATGAATGTGGGCCTTGGACTCGAAGTAgAAgAgTCTTCCTTTACg |
| Shh_B1 | gAggAgggCAgCAAACggAATACCTTGGCGGATGCGGGAAAACAG |
| Shh_B1 | CGGTCTGGTCACTCCATGCTCCAGGTAgAAgAgTCTTCCTTTACg |
| Shh_B1 | gAggAgggCAgCAAACggAAAAGTCGCTGTAGACCAGCCTGCCCT |
| Shh_B1 | GCCTCCTCCTTGTCCAGGAACATGATAgAAgAgTCTTCCTTTACg |
| Shh_B1 | gAggAgggCAgCAAACggAAGTCACAGGGGCGTATGCACCTGCAC |
| Shh_B1 | TTGTCTATGACCACGGTGCCGTGTGTAgAAgAgTCTTCCTTTACg |
| Shh_B1 | gAggAgggCAgCAAACggAATCTCAAAGGGGCAAAGGCCCAGTGC |
| Shh_B1 | AAAGATGGACAGGATGCCATAGCCCTAgAAgAgTCTTCCTTTACg |
| Shh_B1 | gAggAgggCAgCAAACggAACGAGTACCAGTGCACGCCCTCTTTT |
| Shh_B1 | CCATGTCCCTATGTGGTAGAGGATTTAgAAgAgTCTTCCTTTACg |
| Shh_B1 | gAggAgggCAgCAAACggAAAGGGTGGATGGTGTCCGAGTGTAAC |
| Shh_B1 | TCAGCTGGACTTTGCAGCCATTCCCTAgAAgAgTCTTCCTTTACg |
| Shh_B1 | gAggAgggCAgCAAACggAACTTTGGACAGTCCTACTTTTGTTTT |
| Shh_B1 | CCGGGGTCTTTGTTTCTTTGAAGTCTAgAAgAgTCTTCCTTTACg |
| Shh_B1 | gAggAgggCAgCAAACggAACCAAGTTGCCGAGTGGAGAAGAAAA |
| Shh_B1 | GTTTGCCAGTCCCATACCAATAAATTAgAAgAgTCTTCCTTTACg |
| Shh_B1 | gAggAgggCAgCAAACggAATGAAAAGCCTGGTTTTCAGCTTGAC |
| Shh_B1 | ATTTTCATTTCTTTTGTTTCGGAGTTAgAAgAgTCTTCCTTTACg |
| Shh_B1 | gAggAgggCAgCAAACggAATTTGTCTGCTTTGCTTTAGAGTGTC |
| Shh_B1 | AAGAAAACGTCTGGGATCTATGTTTTAgAAgAgTCTTCCTTTACg |
| Shh_B1 | gAggAgggCAgCAAACggAAATAATATAGGTCTACAACACAAAGG |
| Shh_B1 | CAACAAAGAGAATACAAACAAAGGCTAgAAgAgTCTTCCTTTACg |
| Shh_B1 | gAggAgggCAgCAAACggAAATATCCAGTAGGCTGCAGAAAACGG |
| Shh_B1 | ACATCAACTCAAACGCCAAACAAATTAgAAgAgTCTTCCTTTACg |
| Shh_B1 | gAggAgggCAgCAAACggAATTGAAAGTCCCATTCACATCGTTAA |

| | |
|---|---|
| Shh_B1 | ATTATCAAAACAAGTTATTAAAGGCTAgAAgAgTCTTCCTTTACg |
| Shh_B1 | gAggAgggCAgCAAACggAAGTTCCCGATGCGGCCGGAGAGAAAA |
| Shh_B1 | GCCTTCCAAACCCACCGCTTTCTGCTAgAAgAgTCTTCCTTTACg |
| Shh_B1 | gAggAgggCAgCAAACggAACGGACACAAATGATACATAGAAAA |
| Shh_B1 | GAAAATTGTTTGTTCTACTGAAAGTTAgAAgAgTCTTCCTTTACg |
| Shh_B1 | gAggAgggCAgCAAACggAAACTAATTTACAATTCTGTGTATAAA |
| Shh_B1 | TTCGGTAAGTATTGATCTCGCAAAATAgAAgAgTCTTCCTTTACg |
| Shh_B1 | gAggAgggCAgCAAACggAAGGAAAATATTAAAACACAAATAGAA |
| Shh_B1 | ATAGGAAGGGGTCGAAACATTTAGATAgAAgAgTCTTCCTTTACg |
| **Fgf8** | |
| Fgf8_B3 | gTCCCTgCCTCTATATCTTTAGCCCTCTTCCTCCCCTCCTCTTGT |
| Fgf8_B3 | AGTAGATTTGAGGCAGTGGTGGTGATTCCACTCAACTTTAACCCg |
| Fgf8_B3 | gTCCCTgCCTCTATATCTTTTCTCTTGCCTTTTAACCCCGGGCGC |
| Fgf8_B3 | ATAATATGTCCTTTTCTTCATTTAGTTCCACTCAACTTTAACCCg |
| Fgf8_B3 | gTCCCTgCCTCTATATCTTTAAAGTTCGCTTTTCTGGATCGCTAG |
| Fgf8_B3 | GTAGTGGAGGTGAGGCACGTGAAGCTTCCACTCAACTTTAACCCg |
| Fgf8_B3 | gTCCCTgCCTCTATATCTTTACATTGCCACCTTGGATCACGGGCC |
| Fgf8_B3 | CCTCTGCGAAGCACTAGCTTAGTGGTTCCACTCAACTTTAACCCg |
| Fgf8_B3 | gTCCCTgCCTCTATATCTTTAAAGCGCCTGTCCTCCAGTAGCACC |
| Fgf8_B3 | GGCGCTTTACGGATCGTTACCCGGCTTCCACTCAACTTTAACCCg |
| Fgf8_B3 | gTCCCTgCCTCTATATCTTTTTGTCTGCAAATGGGAAGTCCCCGA |
| Fgf8_B3 | GAATAAACCCCTTGCTGGAGCGATGTTCCACTCAACTTTAACCCg |
| Fgf8_B3 | gTCCCTgCCTCTATATCTTTGATGAGTTCGATGGCAGGGGCTCGG |
| Fgf8_B3 | TTATCGCTCGCAGGGCTTAACCCGTTTCCACTCAACTTTAACCCg |
| Fgf8_B3 | gTCCCTgCCTCTATATCTTTTTCTGAACCCTTCGGACCGCTCTGG |
| Fgf8_B3 | CGACGACTGACGAGCTCGCGTTGTTTTCCACTCAACTTTAACCCg |
| Fgf8_B3 | gTCCCTgCCTCTATATCTTTCTTTGCTTGCATGCAGAGGACAAAA |
| Fgf8_B3 | TGTCACCAGGCTCTGCTCCCTCACATTCCACTCAACTTTAACCCg |
| Fgf8_B3 | gTCCCTgCCTCTATATCTTTGACCTGCACGTGCTTCCCACTGGTG |
| Fgf8_B3 | CATGGCGTTGATCTTCTTGTTGCCATTCCACTCAACTTTAACCCg |
| Fgf8_B3 | gTCCCTgCCTCTATATCTTTTAATTTGGCGTGCGAGTCGCCGTCG |
| Fgf8_B3 | GCTTCCGAAGGTGTCCGTTTCCACATTCCACTCAACTTTAACCCg |
| Fgf8_B3 | gTCCCTgCCTCTATATCTTTCGTCTCGGCTCCTTTAATGCGGACG |
| Fgf8_B3 | CTTCTTGTTCATGCAGATGTAATAGTTCCACTCAACTTTAACCCg |
| Fgf8_B3 | gTCCCTgCCTCTATATCTTTGCCATTGCTCTTGCCAATCAGCTTT |
| Fgf8_B3 | CTCGGAGAAGACGCAGTCTTTGCCTTTCCACTCAACTTTAACCCg |
| Fgf8_B3 | gTCCCTgCCTCTATATCTTTTGCCGTGTAGTTGTTCTCGAGCACT |
| Fgf8_B3 | CCATCCTTCGTATTTGGCGTTTTGTTTCCACTCAACTTTAACCCg |
| Fgf8_B3 | gTCCCTgCCTCTATATCTTTGCGGCCTTTCCTTGTGAAGGCCATG |

| | |
|---|---|
| Fgf8_B3 | CTGCCTGGTCTTGGAGCCTTTCCTCTTCCACTCAACTTTAACCCg |
| Fgf8_B3 | gTCCCTgCCTCTATATCTTTCTTCATGAAGTGGACCTCCCGCTGG |
| Fgf8_B3 | CGTTGTTTGGTGGCCTTTCGGCAGTTTCCACTCAACTTTAACCCg |
| Fgf8_B3 | gTCCCTgCCTCTATATCTTTTACAAACTCGAAGCGTCTATGAGGC |
| Fgf8_B3 | TTTGCTCCGCCTGTTGAAAGGATAATTCCACTCAACTTTAACCCg |
| Fgf8_B3 | gTCCCTgCCTCTATATCTTTCTATCGCGGTACGGAATGTCGAGTT |
| Fgf8_B3 | GTGGATCGCTGAGGAGTGTTTTGGTTTCCACTCAACTTTAACCCg |
| Fgf8_B3 | gTCCCTgCCTCTATATCTTTCATTTCATCCTTGTTTTGTATAAAA |
| Fgf8_B3 | TAAGAGCATAGCTGAAAAGTCTTTGTTCCACTCAACTTTAACCCg |
| Fgf8_B3 | gTCCCTgCCTCTATATCTTTAACTCTTGATTCTGTGTTTGATTTT |
| Fgf8_B3 | CATCACATCCTTTCTCAGTTTTGCATTCCACTCAACTTTAACCCg |
| Fgf8_B3 | gTCCCTgCCTCTATATCTTTTAACATGTATATGTACAAAAGAAAA |
| Fgf8_B3 | AGCTTTTAAAATGAATCAACGCGTCTTCCACTCAACTTTAACCCg |
| Fgf8_B3 | gTCCCTgCCTCTATATCTTTTACAGGTAGGAAACTCTGCAGAAAA |
| Fgf8_B3 | GCCTGTAAACCAGGCAAACTCCATGTTCCACTCAACTTTAACCCg |
| **Frem3** | |
| Frem3_B4 | CCTCAACCTACCTCCAACAAGTGCAGATGAGCAGCAGCAGAGCCG |
| Frem3_B4 | TTCATGAGTAACACAACAATGACCAATTCTCACCATATTCGCTTC |
| Frem3_B4 | CCTCAACCTACCTCCAACAATTCCCATCAGATGATCGCCTCCTCC |
| Frem3_B4 | CTGCAATGGCTGTTGGTTGAGGACTATTCTCACCATATTCGCTTC |
| Frem3_B4 | CCTCAACCTACCTCCAACAATGCCTTGTGGTCATAGTTTCGTTGC |
| Frem3_B4 | TAGACCTCCGAGCTGTCACTGCCGCATTCTCACCATATTCGCTTC |
| Frem3_B4 | CCTCAACCTACCTCCAACAAGGTTCCCATATTCATGAAGAGTTTT |
| Frem3_B4 | ATTTCAGGAACCCCTCAAAGCTCTTATTCTCACCATATTCGCTTC |
| Frem3_B4 | CCTCAACCTACCTCCAACAACCGAGAAGGGCAGCACCAACATATC |
| Frem3_B4 | GGTAGGCCAGGAGAGCGGATGTGAGATTCTCACCATATTCGCTTC |
| Frem3_B4 | CCTCAACCTACCTCCAACAAAATAAGGCACGTCTGGGCACAGGGC |
| Frem3_B4 | TTGTCCCTGTGGAAGTTTTGTTCATATTCTCACCATATTCGCTTC |
| Frem3_B4 | CCTCAACCTACCTCCAACAACTTATCCGCGTAAAGTCATCTTGGG |
| Frem3_B4 | ATATGGTGGTCAAGAGGCCGCCAAAATTCTCACCATATTCGCTTC |
| Frem3_B4 | CCTCAACCTACCTCCAACAAAAAACCAAAATAATACATACAAATC |
| Frem3_B4 | AGCCTTATTGGACACACCCATTTATATTCTCACCATATTCGCTTC |
| Frem3_B4 | CCTCAACCTACCTCCAACAAGTCATTCTAGTTTGACCTGAATACA |
| Frem3_B4 | GGTCTGAGCATGGAGCTACATAAATATTCTCACCATATTCGCTTC |
| Frem3_B4 | CCTCAACCTACCTCCAACAAGTTGTCCCGCTACCCTCCCAGGGAA |
| Frem3_B4 | GGGAATATAAAACATCTGAGCATAGATTCTCACCATATTCGCTTC |
| Frem3_B4 | CCTCAACCTACCTCCAACAAGTACGTTTCGCAATGGCATACGGAA |
| Frem3_B4 | TAGCTGGTAAGTAAGGCACACAGTCATTCTCACCATATTCGCTTC |
| Frem3_B4 | CCTCAACCTACCTCCAACAACGATTCCACAGAAGAGCGCCAGGGT |

| | |
|---|---|
| Frem3_B4 | ATTCCAAAATACAGAAAGAGAACAGATTCTCACCATATTCGCTTC |
| Frem3_B4 | CCTCAACCTACCTCCAACAACAGTTGCAGGATTGTCCTCAGCCCC |
| Frem3_B4 | CAATGCTTGAACGGGATGTGCTCCTATTCTCACCATATTCGCTTC |
| Frem3_B4 | CCTCAACCTACCTCCAACAAAATACTTGCCGGCTTGGGGTATATA |
| Frem3_B4 | CAAGTTTTACCAGGCAACGCCTTTCATTCTCACCATATTCGCTTC |
| Frem3_B4 | CCTCAACCTACCTCCAACAATTCAGGGCAATGCCATGCCAGCACT |
| Frem3_B4 | CACTCACAGACTTTACTTCAGTGGTATTCTCACCATATTCGCTTC |
| Frem3_B4 | CCTCAACCTACCTCCAACAAGAAGGCAGCGTGGAGCGGGTAAGTG |
| Frem3_B4 | GAGCCTGTCCCTGCTCGTCTTTATGATTCTCACCATATTCGCTTC |
| Frem3_B4 | CCTCAACCTACCTCCAACAACAATCTAGGGAATATGGACTTATGT |
| Frem3_B4 | ACACGGGAAAACAGTAGTGTGAGGCATTCTCACCATATTCGCTTC |
| Frem3_B4 | CCTCAACCTACCTCCAACAAAATTACTCTTCACATACTTTCAAAA |
| Frem3_B4 | ACCAGGAAGGACTTTTAAAGTCGAAATTCTCACCATATTCGCTTC |
| Frem3_B4 | CCTCAACCTACCTCCAACAAATTTGAGGTATAGGGGAAGAGGATA |
| Frem3_B4 | ATACAAAGCTGGTTTCATTTAGCGTATTCTCACCATATTCGCTTC |
| Frem3_B4 | CCTCAACCTACCTCCAACAACCAAGCGAGCCAGGCACTGAACAGA |
| Frem3_B4 | GAGGGGACATCATTCACAGGAGCAGATTCTCACCATATTCGCTTC |
| Frem3_B4 | CCTCAACCTACCTCCAACAACAAGAACCAAACCCAGTGTAAATAG |
| Frem3_B4 | CCAGCACCTTAATCATGGCTCCTGAATTCTCACCATATTCGCTTC |
| Frem3_B4 | CCTCAACCTACCTCCAACAACTTTACAGCCAAACTGGTAGCTGCC |
| Frem3_B4 | TATTTACACTGTCCAGAAACTCATGATTCTCACCATATTCGCTTC |
| Frem3_B4 | CCTCAACCTACCTCCAACAAAAACGGGCCTGCTAATTGAGTGACA |
| Frem3_B4 | CTGAATAGTTGTGGGGCAACAACTGATTCTCACCATATTCGCTTC |
| Frem3_B4 | CCTCAACCTACCTCCAACAAATAAATAGCTATCAGAGCGCATTTT |
| Frem3_B4 | AGGGAAAATAAAACTGCTCGTTCATATTCTCACCATATTCGCTTC |
| Frem3_B4 | CCTCAACCTACCTCCAACAACATGGGAAAGTGGAATAGAGACTAT |
| Frem3_B4 | ATTGCTCATGGACTGAAGCTATCTGATTCTCACCATATTCGCTTC |
| Frem3_B4 | CCTCAACCTACCTCCAACAAGGAGTTGTTCATTCGGGATCATCTT |
| Frem3_B4 | AATTGCGTGTATCCAAAATGGCCGAATTCTCACCATATTCGCTTC |
| **Hand2** | |
| Hand2_B3 | gTCCCTgCCTCTATATCTTTAGCAAAACCCTCGAGGATCACAGAC |
| Hand2_B3 | CTTCTGGCCCGGCACTCTTCCACCCTTCCACTCAACTTTAACCCg |
| Hand2_B3 | gTCCCTgCCTCTATATCTTTTGTGTTGGAGTTCACTCAAATCCCC |
| Hand2_B3 | TGGACGACTCTTCTTGAAGTCCTGGTTCCACTCAACTTTAACCCg |
| Hand2_B3 | gTCCCTgCCTCTATATCTTTACCGTCGCGCCTGGGAGCCCAGCGC |
| Hand2_B3 | TCGACCTCCGGCCTCCCCGGCACCATTCCACTCAACTTTAACCCg |
| Hand2_B3 | gTCCCTgCCTCTATATCTTTGGACGCTCCCCGTTGCTCCCCGGGG |
| Hand2_B3 | CCTCAGAGTTGTGTCCAGTTCCCCATTCCACTCAACTTTAACCCg |
| Hand2_B3 | gTCCCTgCCTCTATATCTTTCCAGGGTGGTGGTGAGGGAAGCCCC |

| | |
|---|---|
| Hand2_B3 | GGGTAACCCTCGTGGTGGTGGTGCATTCCACTCAACTTTAACCCg |
| Hand2_B3 | gTCCCTgCCTCTATATCTTTATGGCAGCGGCCCGTCGCCGCTGCG |
| Hand2_B3 | CCAGCCGTGGAAGTAGGGGCTGTCTTTCCACTCAACTTTAACCCg |
| Hand2_B3 | gTCCCTgCCTCTATATCTTTGTGCTCAGCAGCGCTCCCGCGCCCC |
| Hand2_B3 | GCCGTGCCCCTGCGCTTCACCGGCCTTCCACTCAACTTTAACCCg |
| Hand2_B3 | gTCCCTgCCTCTATATCTTTTGCGTGCGGCGCCGCTCCTTGCGGT |
| Hand2_B3 | AGCTCCGCGAAGGCGCTGTTGATGCTTCCACTCAACTTTAACCCg |
| Hand2_B3 | gTCCCTgCCTCTATATCTTTGCGGGCACGTTGGGGATGCACTCGC |
| Hand2_B3 | GTCTTGATCTTGGAGAGCTTGGTGTTTCCACTCAACTTTAACCCg |
| Hand2_B3 | gTCCCTgCCTCTATATCTTTGCGATGTAGCTGGTGGCTAGGCGCA |
| Hand2_B3 | TCCTTGGCCAGCAGGTCCATGAGGTTTCCACTCAACTTTAACCCg |
| Hand2_B3 | gTCCCTgCCTCTATATCTTTCGGAAGGCCTCGGCCTCGCTCTGCT |
| Hand2_B3 | CCGCCGCCGCGCTGCTTGAGGTCGGTTCCACTCAACTTTAACCCg |
| Hand2_B3 | gTCCCTgCCTCTATATCTTTCGCTTGTCCTCCTTGCACTCTGCGC |
| Hand2_B3 | TTCAAGAGTTCATTCAGCTCTTTCTTTCCACTCAACTTTAACCCg |
| Hand2_B3 | gTCCCTgCCTCTATATCTTTTTCTTGTCGTTGCTGCCGACGGTAC |
| Hand2_B3 | TGAGGCCAGCCAGTCCTGCCCTTGCTTCCACTCAACTTTAACCCg |
| Hand2_B3 | gTCCCTgCCTCTATATCTTTTGCTTGAGCTCCAAGGCCCAGACGT |
| Hand2_B3 | CTCACTCGCGTGCTCGGCCACGCTCTTCCACTCAACTTTAACCCg |
| Hand2_B3 | gTCCCTgCCTCTATATCTTTCTCCTCGGACACTGTCGCCCAGGGG |
| Hand2_B3 | CGTGGGCTCCCTCTGCCAGCCCCGGTTCCACTCAACTTTAACCCg |
| Hand2_B3 | gTCCCTgCCTCTATATCTTTCCCGGACCCTGTAGCATCGTAGGGG |
| Hand2_B3 | CTGTGACCTTGACCCGTTCCGGGACTTCCACTCAACTTTAACCCg |
| Hand2_B3 | gTCCCTgCCTCTATATCTTTACGGGCCCTGTATAGGCCCGAGGGC |
| Hand2_B3 | AGAGCCCGTGTGCCGGTATGGGGTCTTCCACTCAACTTTAACCCg |
| Hand2_B3 | gTCCCTgCCTCTATATCTTTGGGCACAGGCTAGTCCCAGGCTGCC |
| Hand2_B3 | AATACTGAAAGTCTTGAGGGTGGAGTTCCACTCAACTTTAACCCg |
| Hand2_B3 | gTCCCTgCCTCTATATCTTTTGCATATAAATAAAGCGGGAGGTTC |
| Hand2_B3 | CTGGGGCTTCGGTTTGTCTGCGGACTTCCACTCAACTTTAACCCg |
| Hand2_B3 | gTCCCTgCCTCTATATCTTTCTGCCCGGGCCTCCTCTCCATGGGG |
| Hand2_B3 | CGAACTGAAAGGCGGAGTGCTCCTTTTCCACTCAACTTTAACCCg |
| Hand2_B3 | gTCCCTgCCTCTATATCTTTGAGGTGCGCTTCACAATGCCTGGGG |
| Hand2_B3 | CATCTGCAGGAGGCAGTGCGGCCCCTTCCACTCAACTTTAACCCg |
| Hand2_B3 | gTCCCTgCCTCTATATCTTTGGTGCAGTTCCACAATCCAGGACCA |
| Hand2_B3 | TAAACATTGGCACAGCACTCCCGTGTTCCACTCAACTTTAACCCg |
| Hand2_B3 | gTCCCTgCCTCTATATCTTTCTTCAAAGACAGACCGACCACAAAA |
| Hand2_B3 | AAGACGGAAGTGCACAAGATAGATCTTCCACTCAACTTTAACCCg |
| Hand2_B3 | gTCCCTgCCTCTATATCTTTAGCACTGGCCCAAACTGCCGAGCCC |
| Hand2_B3 | GCGCCGTGCAGGCAGCGGTGGAGACTTCCACTCAACTTTAACCCg |

| Gli3 | |
|---|---|
| Gli3_B2 | CCTCgTAAATCCTCATCAAAGGAGGTTGTGCTGGAGGCAACCGCC |
| Gli3_B2 | GGGCTGTCCAGGACTTTCATCCTCAAAATCATCCAgTAAACCgCC |
| Gli3_B2 | CCTCgTAAATCCTCATCAAAAATGGCGTTCCTTCTCTCTCTGTGA |
| Gli3_B2 | GCCCTGCCCTCCTTGTGGCTGCATGAAATCATCCAgTAAACCgCC |
| Gli3_B2 | CCTCgTAAATCCTCATCAAAAGAGGGCTCTTCGCTGATTTTACTG |
| Gli3_B2 | TAATGATGCCCTTTCCTCACTCGATAAATCATCCAgTAAACCgCC |
| Gli3_B2 | CCTCgTAAATCCTCATCAAATATGGATGCATGGATCTCCTTCTTT |
| Gli3_B2 | CGGCAGGGCATGCTCGTGAAGGTGCAAATCATCCAgTAAACCgCC |
| Gli3_B2 | CCTCgTAAATCCTCATCAAAGTCCATCGCAAAGAGCGCACCTCGG |
| Gli3_B2 | ATGAGGTTCCATGTAGCCATTCCTTAAATCATCCAgTAAACCgCC |
| Gli3_B2 | CCTCgTAAATCCTCATCAAATGTCTTGCGTCAATTGGTACTGGGG |
| Gli3_B2 | GGTTCGTAATGGTATCGTCCCTCGTAAATCATCCAgTAAACCgCC |
| Gli3_B2 | CCTCgTAAATCCTCATCAAAGGCATATGCAGCGGTGGAATGGGCG |
| Gli3_B2 | TAGGTTGGGCTACTCGACAATGCTGAAATCATCCAgTAAACCgCC |
| Gli3_B2 | CCTCgTAAATCCTCATCAAAGAAATCCGGATGAAGGGAAGGTCTG |
| Gli3_B2 | TCGGAGGCCCCGTGAGTTCTGTGAGAAATCATCCAgTAAACCgCC |
| Gli3_B2 | CCTCgTAAATCCTCATCAAATACGGATGAGGAGGGCTGAAAGGAG |
| Gli3_B2 | CGAATGTAGTCCATGTACGGGTTGAAAATCATCCAgTAAACCgCC |
| Gli3_B2 | CCTCgTAAATCCTCATCAAACTGTGATCGGAGAGTGGCGAGATGG |
| Gli3_B2 | GTCCGAATCATCGTCTGCAGGTCAAAAATCATCCAgTAAACCgCC |
| Gli3_B2 | CCTCgTAAATCCTCATCAAAAGAAGGGTGACCAAGGAGTTTGGAG |
| Gli3_B2 | GCCGAGGAGCTGCTGCGTGAGGTATAAATCATCCAgTAAACCgCC |
| Gli3_B2 | CCTCgTAAATCCTCATCAAAGCAGATAAGTGACCGTATGACCCAC |
| Gli3_B2 | AAGCTGAGAGCTGGGCTGATTGCACAAATCATCCAgTAAACCgCC |
| Gli3_B2 | CCTCgTAAATCCTCATCAAACAACCTTCAAAAGTGCATTTGTGGG |
| Gli3_B2 | TTTTCCAGTCTGGAGTAAGCCTTTGAAATCATCCAgTAAACCgCC |
| Gli3_B2 | CCTCgTAAATCCTCATCAAAGTGTGTGATCTCAAGTGGGTTTTCA |
| Gli3_B2 | TGCTCACAGACATATGGTTTCTCTCAAATCATCCAgTAAACCgCC |
| Gli3_B2 | CCTCgTAAATCCTCATCAAATTGGAGAAAGCTTTGTTGCAGGTTT |
| Gli3_B2 | TTTTGGTGCTTGGCCCTGTCCGATGAAATCATCCAgTAAACCgCC |
| Gli3_B2 | CCTCgTAAATCCTCATCAAATATGGTTTCTCATTTGAATGAGTTC |
| Gli3_B2 | TTTGTGCAGCCCGGGATCTTGCAAAAAATCATCCAgTAAACCgCC |
| Gli3_B2 | CCTCgTAAATCCTCATCAAAGGCCCCGCCATCGCTGGACCTTCTT |
| Gli3_B2 | TAACATATGCCTTCTGCTAAAGCTGAAATCATCCAgTAAACCgCC |
| Gli3_B2 | CCTCgTAAATCCTCATCAAATGTCGTGTTCCCTTGCATCTCATGC |
| Gli3_B2 | CATTCTTACAGGATCACTAGCTCTTAAATCATCCAgTAAACCgCC |
| Gli3_B2 | CCTCgTAAATCCTCATCAAATACTCTAGTGGAAGAAAGGTTAGAC |
| Gli3_B2 | GACATTGTTTAGGCTGTTGAAACGCAAATCATCCAgTAAACCgCC |

| | |
|---|---|
| Gli3_B2 | CCTCgTAAATCCTCATCAAAAGGATCATGTTGTTGTACAGGCCAA |
| Gli3_B2 | GCATTTCGTGATTGACTTTGCTGCTAAATCATCCAgTAAACCgCC |
| Gli3_B2 | CCTCgTAAATCCTCATCAAATGCATGTAGCCTTGCTGTTGGACAA |
| Gli3_B2 | AAATTGTATGCACTGTTTGCTGAGTAAATCATCCAgTAAACCgCC |
| Gli3_B2 | CCTCgTAAATCCTCATCAAAGGGTGGTTAATGGACGAATTCTGCT |
| Gli3_B2 | GGTTGCATGCTGTTGAAGGAATTACAAATCATCCAgTAAACCgCC |
| Gli3_B2 | CCTCgTAAATCCTCATCAAACCTGGGGAGAGGAGTTCAGAAGAAT |
| Gli3_B2 | TCAACTGTGCTGGTTACTTGATTGGAAATCATCCAgTAAACCgCC |
| Gli3_B2 | CCTCgTAAATCCTCATCAAACCCAACAGACCACTGCTGTCAATGT |
| Gli3_B2 | AAGATGGCGTCAAAATCAATCTGCAAAATCATCCAgTAAACCgCC |
| **Cyp26b1** | |
| Cyp26b1_B1 | gAggAgggCAgCAAACggAAAGGACCACTGACACTAGGCAGGCGG |
| Cyp26b1_B1 | CACAGCTGTTGGGAGACGGCCAGCATAgAAgAgTCTTCCTTTACg |
| Cyp26b1_B1 | gAggAgggCAgCAAACggAATCCCGGGTGGCAGCCCAGCGGAGTT |
| Cyp26b1_B1 | TTGGGGATGGGCAGCTTGCAGCTCTTAgAAgAgTCTTCCTTTACg |
| Cyp26b1_B1 | gAggAgggCAgCAAACggAACGTTGCCATATTTCTCCCTTCTGGA |
| Cyp26b1_B1 | GCCGCCCCAACAAGTGTGTCTTGAATAgAAgAgTCTTCCTTTACg |
| Cyp26b1_B1 | gAggAgggCAgCAAACggAATCTCTGCCCCGGTCACTCTGATTAA |
| Cyp26b1_B1 | GCTCACCCATCAGAATCTTGCGGACTAgAAgAgTCTTCCTTTACg |
| Cyp26b1_B1 | gAggAgggCAgCAAACggAAGAGGCCACTCTGTGCTCACCAGGCT |
| Cyp26b1_B1 | TGGGGCCTAGTAGGGTCCTTGTGCTTAgAAgAgTCTTCCTTTACg |
| Cyp26b1_B1 | gAggAgggCAgCAAACggAACCAGGGCCTCGTGGCTGAAGATCTT |
| Cyp26b1_B1 | CTAGCTGGATCTTGGGAAGGTAACTTAgAAgAgTCTTCCTTTACg |
| Cyp26b1_B1 | gAggAgggCAgCAAACggAATCCACATCCTCAATGTGTCCTGGAT |
| Cyp26b1_B1 | AGACATTAATGGGATCAGGGTTACTTAgAAgAgTCTTCCTTTACg |
| Cyp26b1_B1 | gAggAgggCAgCAAACggAAGGAAGGTTAACTTCTGGGCCTCGAA |
| Cyp26b1_B1 | ATCCCAGAAGGACCCGGATGGCCATTAgAAgAgTCTTCCTTTACg |
| Cyp26b1_B1 | gAggAgggCAgCAAACggAAGATTGAGTTCCTCATCAGAGAGGCG |
| Cyp26b1_B1 | CAAACTGCTGAAAGACCTGGAAGAGTAgAAgAgTCTTCCTTTACg |
| Cyp26b1_B1 | gAggAgggCAgCAAACggAACCACAGGCAGGGAGAACACGTTCTC |
| Cyp26b1_B1 | CCCTTCTGTAGCCACTGAATGGCATTAgAAgAgTCTTCCTTTACg |
| Cyp26b1_B1 | gAggAgggCAgCAAACggAATCTGCAGGGTCTCGCGAGCCCGAAT |
| Cyp26b1_B1 | TCTCTCGAATGGCCTTCTCCAGACTTAgAAgAgTCTTCCTTTACg |
| Cyp26b1_B1 | gAggAgggCAgCAAACggAAAGTCCTTCCCCTGCGAATTCTGGAA |
| Cyp26b1_B1 | CAATCAAAATATCGAGCGCGTCTGCTAgAAgAgTCTTCCTTTACg |
| Cyp26b1_B1 | gAggAgggCAgCAAACggAAGGCTGTTGTAGCATAGGCAGCAAAA |
| Cyp26b1_B1 | CTGCATTATGAGTGAGGTGCTGGAGTAgAAgAgTCTTCCTTTACg |
| Cyp26b1_B1 | gAggAgggCAgCAAACggAATTCGAAGACAGCAGGATGCTTCAAG |

| | |
|---|---|
| Cyp26b1_B1 | GTTGCCCCGGAGTTCCTCCCGCAGCTAgAAgAgTCTTCCTTTACg |
| Cyp26b1_B1 | gAggAgggCAgCAAACggAAACAGATGCATCCATTGTGGAGGATA |
| Cyp26b1_B1 | GATGTTGTCTACCCGGAACGCTCCGTAgAAgAgTCTTCCTTTACg |
| Cyp26b1_B1 | gAggAgggCAgCAAACggAACACACAGTCCAAGTAATGGAGGCTG |
| Cyp26b1_B1 | ACTAAAGAGCCGGAGGACTTCCTTGTAgAAgAgTCTTCCTTTACg |
| Cyp26b1_B1 | gAggAgggCAgCAAACggAACATTACACTCCAGCCTTTTGGGATT |
| Cyp26b1_B1 | TGTGTCATGTGTATCCCGTATACTATAgAAgAgTCTTCCTTTACg |
| Cyp26b1_B1 | gAggAgggCAgCAAACggAAAACGTCCACATCCTTGAAGACCGGT |
| Cyp26b1_B1 | GTCTTGACCGAAGCGGTCTGGGTCATAgAAgAgTCTTCCTTTACg |
| Cyp26b1_B1 | gAggAgggCAgCAAACggAAGAACCTTCCATCCTTGTCTTCAGTG |
| Cyp26b1_B1 | TATTCCACCGCCAAATGGGAGATAGTAgAAgAgTCTTCCTTTACg |
| Cyp26b1_B1 | gAggAgggCAgCAAACggAATTGAGGAAAAGCTTTGCCAGATTTT |
| Cyp26b1_B1 | GTGCTGGCCAGTTCGATGGCTAGGGTAgAAgAgTCTTCCTTTACg |
| Cyp26b1_B1 | gAggAgggCAgCAAACggAAGTCCGTGTGGCCAGCTCAAATCGGC |
| Cyp26b1_B1 | ACAGGAACCGGCATCACACGTGGAATAgAAgAgTCTTCCTTTACg |
| Cyp26b1_B1 | gAggAgggCAgCAAACggAAGTTCTGGTTGGAATCAAGTCCAAAA |
| Cyp26b1_B1 | CATTGTTTCTGTCTCTGTTATGATTTAgAAgAgTCTTCCTTTACg |
| Cyp26b1_B1 | gAggAgggCAgCAAACggAAGGAAGAATGGAAACATACACTGGGG |
| Cyp26b1_B1 | TTTGCATTATTCCCCTGACAGGGGCTAgAAgAgTCTTCCTTTACg |
| Cyp26b1_B1 | gAggAgggCAgCAAACggAAGTTTCTGTACTTGATTTGTATTGTT |
| Cyp26b1_B1 | TGATGTCGCTGTTTAGCTTCTCTGTTAgAAgAgTCTTCCTTTACg |
| **Pdgfra** | |
| Pdgfra_B1 | GAGACACCCCAGGGCCAGGAATGTCTAgAAgAgTCTTCCTTTACg |
| Pdgfra_B1 | gAggAgggCAgCAAACggAACAAAGACTTCGCCGGACCTGTCACC |
| Pdgfra_B1 | GAAAATAGAGGGGATGGGGTACTGGTAgAAgAgTCTTCCTTTACg |
| Pdgfra_B1 | gAggAgggCAgCAAACggAAGCCTGTGCATTTCAGGGTGAAGGGG |
| Pdgfra_B1 | CGGGTGCTGCCAACTCAGCTCACTCTAgAAgAgTCTTCCTTTACg |
| Pdgfra_B1 | gAggAgggCAgCAAACggAAAATCTCCACATCTTCGTCCATGCTA |
| Pdgfra_B1 | GAGGCCACTGTTGTTTTCTTCCGTGTAgAAgAgTCTTCCTTTACg |
| Pdgfra_B1 | gAggAgggCAgCAAACggAAATTCTTTACGCTGAGCAAAGACACA |
| Pdgfra_B1 | GTACATCCCCGTGCCCGCAGCCGTGTAgAAgAgTCTTCCTTTACg |
| Pdgfra_B1 | gAggAgggCAgCAAACggAACAATTGCGTGTGGTTGTGAAAGCAC |
| Pdgfra_B1 | ATCCTTGCCTTCGATCTCATTGTCTTAgAAgAgTCTTCCTTTACg |
| Pdgfra_B1 | gAggAgggCAgCAAACggAAGTCTGGGTCTGGCACATAGATGTAA |
| Pdgfra_B1 | CAGCATCATGGATGGCACGAAGGCCTAgAAgAgTCTTCCTTTACg |
| Pdgfra_B1 | gAggAgggCAgCAAACggAAATCTTCCTCCACCGCAATGAAGTGG |
| Pdgfra_B1 | GGTACGACAAGGGATGAGCGATGAGTAgAAgAgTCTTCCTTTACg |
| Pdgfra_B1 | gAggAgggCAgCAAACggAAAAGAGAGACCTGCGCGTTCGGGGAG |
| Pdgfra_B1 | CACCACCTTGGAATCCTCCTTGTTATAgAAgAgTCTTCCTTTACg |

| | |
|---|---|
| Pdgfra_B1 | gAggAgggCAgCAAACggAACCCTTGCTTGTTGTCATAGTAGGCA |
| Pdgfra_B1 | GGCTCCCGATGGAAAGTTGCCAAAGTAgAAgAgTCTTCCTTTACg |
| Pdgfra_B1 | gAggAgggCAgCAAACggAATCCGTTGGCGATGGTTTCGCACACG |
| Pdgfra_B1 | GATGTAAGTGTCTGTGTGGATCGCGTAgAAgAgTCTTCCTTTACg |
| Pdgfra_B1 | gAggAgggCAgCAAACggAACTCAGATGTGGCTTTCCATTTGTGC |
| Pdgfra_B1 | TTTGCTGGCCTCGATATTAACAGGATAgAAgAgTCTTCCTTTACg |
| Pdgfra_B1 | gAggAgggCAgCAAACggAAGATGTTTTCTCCCGTCTTGAGCACA |
| Pdgfra_B1 | ATTATCGAGCACCACACAGGTGACTTAgAAgAgTCTTCCTTTACg |
| Pdgfra_B1 | gAggAgggCAgCAAACggAAGTAGGTCCATTCGAGATCCACCACC |
| Pdgfra_B1 | GACGCCTCTTTCTTTCGCTTTGCCGTAgAAgAgTCTTCCTTTACg |
| Pdgfra_B1 | gAggAgggCAgCAAACggAAGGGCGACTTGGATTCCACCATCTTG |
| Pdgfra_B1 | GGTCAAGGTGCAAGTCAACTTCAAATAgAAgAgTCTTCCTTTACg |
| Pdgfra_B1 | gAggAgggCAgCAAACggAAGGAGTCCTTCACCGTGGCATTGAGA |
| Pdgfra_B1 | GTGTCTGACAGCACACTCGTACTCGTAgAAgAgTCTTCCTTTACg |
| Pdgfra_B1 | gAggAgggCAgCAAACggAACTTCACTTTGTCGGAGTCCTGCGTT |
| Pdgfra_B1 | TTTATCATGAACTGCAATGTCGATTTAgAAgAgTCTTCCTTTACg |
| Pdgfra_B1 | gAggAgggCAgCAAACggAAGAACTTGGGCTCCAAGTGAATGAAT |
| Pdgfra_B1 | ATGAAGATTTGCCGACTCCACTAATTAgAAgAgTCTTCCTTTACg |
| Pdgfra_B1 | gAggAgggCAgCAAACggAACACATCCACAACAAAGCTTTTCACT |
| Pdgfra_B1 | GGTGATTTTAGGCGGAGGGTAGGCCTAgAAgAgTCTTCCTTTACg |
| Pdgfra_B1 | gAggAgggCAgCAAACggAAGATCAGAGTCACGTTGTCCTTCAGC |
| Pdgfra_B1 | GCTGGTGAGCATCTCAGTGAAATTCTAgAAgAgTCTTCCTTTACg |
| Pdgfra_B1 | gAggAgggCAgCAAACggAAATAGCTGGCTTCTTGCGTCTGGATT |
| Pdgfra_B1 | GGCACGAATCAGTTTCAATATGCATTAgAAgAgTCTTCCTTTACg |
| Pdgfra_B1 | gAggAgggCAgCAAACggAAGGTGTAGTATCCGCTGTCTTCTTCT |
| Pdgfra_B1 | AACGTCCCCATCATTCTGCGCGATATAgAAgAgTCTTCCTTTACg |
| Pdgfra_B1 | gAggAgggCAgCAAACggAAAATTTGCAATGAAAATGTGTAGTTT |
| Pdgfra_B1 | CAGATCCAAAATGGCTGCCTTGACTTAgAAgAgTCTTCCTTTACg |
| Pdgfra_B1 | gAggAgggCAgCAAACggAAACCAGAAGAACCGTGGTGCTCATCC |
| Pdgfra_B1 | TTTAGTCAAACACTCCACGGTCTGCTAgAAgAgTCTTCCTTTACg |
| Pdgfra_B1 | gAggAgggCAgCAAACggAACAGTCGGAACTAGAATTGCACTTTT |
| Pdgfra_B1 | TCAGAGCCATTGGCATTCAAAGGGGTAgAAgAgTCTTCCTTTACg |
| Pdgfra_B1 | gAggAgggCAgCAAACggAATCATCCAGGTAAGTGTGTGTGGTGA |
| Pdgfra_B1 | GTCACCTGACTCTCTACGACATTCTTAgAAgAgTCTTCCTTTACg |
| Pdgfra_B1 | gAggAgggCAgCAAACggAAGCCAGCGTCTCTTCCAATTTTGGAA |
| Pdgfra_B1 | AGTTCGTTCCTCGCAACACATCGAATAgAAgAgTCTTCCTTTACg |
| Pdgfra_B1 | gAggAgggCAgCAAACggAAAATTTCATTTCCCGAGAAGAAAATC |
| Pdgfra_B1 | AGTTCAGAATGTAATGCTTGCGCCATAgAAgAgTCTTCCTTTACg |
| Pdgfra_B1 | gAggAgggCAgCAAACggAAAGCACCAGAACTGCGGCCGCCACTG |

| | |
|---|---|
| Pdgfra_B1 | ATGAGCGAGATGATGACAATCACAATAgAAgAgTCTTCCTTTACg |
| Pdgfra_B1 | gAggAgggCAgCAAACggAATTCTGCTTCCATATGATGACCAGAA |
| Pdgfra_B1 | ACCCGCCACCTGATCTCATACCTGGTAgAAgAgTCTTCCTTTACg |
| Pdgfra_B1 | gAggAgggCAgCAAACggAATGACCGTCTGGGCTGATGGACTCAA |
| Pdgfra_B1 | TGCATCGGATCCACATAGATGTACTTAgAAgAgTCTTCCTTTACg |
| Pdgfra_B1 | gAggAgggCAgCAAACggAAAACTCCCATCTGGAATCGTAGGGTA |
| Pdgfra_B1 | CGACCGAGCACTAGTCCATCGCGAGTAgAAgAgTCTTCCTTTACg |
| Pdgfra_B1 | gAggAgggCAgCAAACggAATTCCCAAAAGCGCCAGAGCCAAGGA |
| Pdgfra_B1 | AAGCCGTACGCCGTTCCTTCAACAATAgAAgAgTCTTCCTTTACg |
| Pdgfra_B1 | gAggAgggCAgCAAACggAAACCTTCATCACTGGCTGGGAGTGGC |
| Pdgfra_B1 | GCAGTGGGTTTTAACATTTTCACAGTAgAAgAgTCTTCCTTTACg |
| Pdgfra_B1 | gAggAgggCAgCAAACggAACAACTCTGACATCAGGGCCTGTTTT |
| Pdgfra_B1 | ATGGGGCCCGAGATGGGTCATTATCTAgAAgAgTCTTCCTTTACg |
| Pdgfra_B1 | gAggAgggCAgCAAACggAAGGCTCCCAGCAAGTTCACAATGTTC |
| Pdgfra_B1 | GATGTAAATGGGTCCTGATTTCGTGTAgAAgAgTCTTCCTTTACg |
| Pdgfra_B1 | gAggAgggCAgCAAACggAAATCTCCATAGAAGCAGTACTCGGTA |
| Pdgfra_B1 | CCTGTTCTTGTGCAGGTAATTTACCTAgAAgAgTCTTCCTTTACg |
| Pdgfra_B1 | gAggAgggCAgCAAACggAACATCCCAAAGATGTCTAAGTCTTTT |
| **Gdf5** | |
| Gdf5_B2 | CCTCgTAAATCCTCATCAAACCCCTCTGCAGGACTGTCTGAAACC |
| Gdf5_B2 | CTGCAAGGTGATGCCCACCTCCCCTAAATCATCCAgTAAACCgCC |
| Gdf5_B2 | CCTCgTAAATCCTCATCAAACAGGCCACGGCTGAAGGGAACCTGT |
| Gdf5_B2 | GTTCAGCCAGCTCCTGTCCCCGTCCAAATCATCCAgTAAACCgCC |
| Gdf5_B2 | CCTCgTAAATCCTCATCAAAGTGTCCAGTCCCATAGTGGAAGTGA |
| Gdf5_B2 | CTCTTGAAATCGGTTGTCTTAATGTAAATCATCCAgTAAACCgCC |
| Gdf5_B2 | CCTCgTAAATCCTCATCAAACGAGAGAATGTCTCTTGGAGCGTTA |
| Gdf5_B2 | TGAAAGAAGTGGAACGCCGAGGCCAAAATCATCCAgTAAACCgCC |
| Gdf5_B2 | CCTCgTAAATCCTCATCAAACTTCCACCAGAGTGACAAGCGAAAA |
| Gdf5_B2 | AGGGCACCGAGGGAAATTTCAAGAGAAATCATCCAgTAAACCgCC |
| Gdf5_B2 | CCTCgTAAATCCTCATCAAAAACTTTCATCCTCTGATCTTCCTTT |
| Gdf5_B2 | CCAAAGCAGTAAAGGGACGTATCTGAAATCATCCAgTAAACCgCC |
| Gdf5_B2 | CCTCgTAAATCCTCATCAAATAGATCCAGGGAGAGGCAGGTCCAG |
| Gdf5_B2 | TTCTGGGTGGCTCAACACAGCGGGGAAATCATCCAgTAAACCgCC |
| Gdf5_B2 | CCTCgTAAATCCTCATCAAACCTAGCTCCAGGAGGGATCTCCCCG |
| Gdf5_B2 | CTTGCTGTCTGCCTCCAACAGCCCCAAATCATCCAgTAAACCgCC |
| Gdf5_B2 | CCTCgTAAATCCTCATCAAAACCCGCCCTGGGCAGTGGGCTTCGC |
| Gdf5_B2 | AGGCCCATGGTGCGCTGTCGTAGAGAAATCATCCAgTAAACCgCC |
| Gdf5_B2 | CCTCgTAAATCCTCATCAAAGGCTCTGGCCTTCGGAGCGCCCGCG |
| Gdf5_B2 | GTTAGGGTCATCGCTCTGGGTCAGGAAATCATCCAgTAAACCgCC |

| | |
|---|---|
| Gdf5_B2 | CCTCgTAAATCCTCATCAAATGCCTCCGTGGCCCCAGCCCGAGGG |
| Gdf5_B2 | TTGCCTACTCAAGAGGTGGCCTTTCAAATCATCCAgTAAACCgCC |
| Gdf5_B2 | CCTCgTAAATCCTCATCAAATCTCGGAGCGACAGTCCTGGCCGCT |
| Gdf5_B2 | TGCGGCCTTGCTGCTTGAGTGCTGGAAATCATCCAgTAAACCgCC |
| Gdf5_B2 | CCTCgTAAATCCTCATCAAATGGGCTGCTGGGCTCTTTGCCTTTT |
| Gdf5_B2 | CTGAAAGTGTCTTTGGGGTCTCTTTAAATCATCCAgTAAACCgCC |
| Gdf5_B2 | CCTCgTAAATCCTCATCAAATACTCGTGCGGCGTTATCACCGGCG |
| Gdf5_B2 | GAAAGCGTGCGGTACAGAGATAGCAAAATCATCCAgTAAACCgCC |
| Gdf5_B2 | CCTCgTAAATCCTCATCAAATCTAGCTTGACGCTACCGTTGCCCC |
| Gdf5_B2 | CTGGTGATGGTGTTGGCTAGTCCTGAAATCATCCAgTAAACCgCC |
| Gdf5_B2 | CCTCgTAAATCCTCATCAAAAGCACTAATGTCGAAAATGTATTTT |
| Gdf5_B2 | AGCCCCTAGTAAGCCATCCTTCTCTAAATCATCCAgTAAACCgCC |
| Gdf5_B2 | CCTCgTAAATCCTCATCAAAGGGCTTCTTCCTAAGAATACGTAGT |
| Gdf5_B2 | TCCTGGCAGACCTCGCCAAGCATCCAAATCATCCAgTAAACCgCC |
| Gdf5_B2 | CCTCgTAAATCCTCATCAAAGGATAGCTTCAACTGCACAATCTTA |
| Gdf5_B2 | GGCCGCCTGTCGGTTGGTTGGGCAGAAATCATCCAgTAAACCgCC |
| Gdf5_B2 | CCTCgTAAATCCTCATCAAACCGCAGGTCAATGGGCCGGCCTTTT |
| Gdf5_B2 | CCTCCCTGTTCGGTCAAACCCTGCAAAATCATCCAgTAAACCgCC |
| Gdf5_B2 | CCTCgTAAATCCTCATCAAATGACCTGGCCTTGATTTCGTTAAAA |
| Gdf5_B2 | TTCATAAACAGTTTTGTCATCTTGGAAATCATCCAgTAAACCgCC |
| Gdf5_B2 | CCTCgTAAATCCTCATCAAACCTCTTCCTCCTCTGGTTGAATAAA |
| Gdf5_B2 | CCCTTGACGCGTTGACAGCGGAGCCAAATCATCCAgTAAACCgCC |
| Gdf5_B2 | CCTCgTAAATCCTCATCAAATTAAAGTTCACGTGGAGGGGCTTTT |
| Gdf5_B2 | ATGATCCAGTCATCCCAGCCCATATAAATCATCCAgTAAACCgCC |
| Gdf5_B2 | CCTCgTAAATCCTCATCAAATGATACGCCTCATACTCCAACGGAG |
| Gdf5_B2 | AGAGGAAATTCACAGAGTCCTTCACAAATCATCCAgTAAACCgCC |
| Gdf5_B2 | CCTCgTAAATCCTCATCAAATGGTTGGTGGGCTCTAGGTGTGACC |
| Gdf5_B2 | GAGTTCATCAAGGTTTGGATGACTGAAATCATCCAgTAAACCgCC |
| Gdf5_B2 | CCTCgTAAATCCTCATCAAACGCGTCGGAATGCAACATGTAGGGG |
| Gdf5_B2 | ATGTACAGGATGCTGATGGGGCTTAAAATCATCCAgTAAACCgCC |
| Gdf5_B2 | CCTCgTAAATCCTCATCAAATTGTAGACCACGTTATTCGCTGAGT |
| Gdf5_B2 | GACTCCACAACCATGTCCTCGTATTAAATCATCCAgTAAACCgCC |
| Gdf5_B2 | CCTCgTAAATCCTCATCAAAGGCTGATCCTGCTACCTACAACCAC |
| Gdf5_B2 | TTTGATACAGTTTGTCATTCATCTGAAATCATCCAgTAAACCgCC |
| Gdf5_B2 | CCTCgTAAATCCTCATCAAATGCGGTGAGTGACATTTCACGCATA |
| Gdf5_B2 | GTTCCATCCCGCCGCTGCCCATGCGAAATCATCCAgTAAACCgCC |
| Gdf5_B2 | CCTCgTAAATCCTCATCAAAGCCAGCAGTCTGATGCTATTAGGTT |
| Gdf5_B2 | GGCTATGCCCATGGCAAAGTCTGTCAAATCATCCAgTAAACCgCC |
| Gdf5_B2 | CCTCgTAAATCCTCATCAAATGGCTGTAGCAGTCTCACATTCCTT |

| | |
|---|---|
| Gdf5_B2 | TCGTCAAAAGTAATACTCGTGTCTGAAATCATCCAgTAAACCgCC |
| Gdf5_B2 | CCTCgTAAATCCTCATCAAAACAGTATCCTTCGAATGTGCAGTTC |
| Gdf5_B2 | CTGTTTGGGCTCTGACAAGTTTAGTAAATCATCCAgTAAACCgCC |
| Gdf5_B2 | CCTCgTAAATCCTCATCAAACCCTCTCACCCTTAATATACATCAG |
| Gdf5_B2 | ATTGCATAAAGTCCCAAGCGAAGAAAAATCATCCAgTAAACCgCC |
| Gdf5_B2 | CCTCgTAAATCCTCATCAAACAATGAATGAAGACATGAAACATCT |
| Gdf5_B2 | TAGAGACATTCTAGCATGTCTATGCAAATCATCCAgTAAACCgCC |
| Gdf5_B2 | CCTCgTAAATCCTCATCAAACGAGTAGAGCTATACAATATCAAGA |
| Gdf5_B2 | CAAAGAAAGCTCTTTGTTTTCAGTCAAATCATCCAgTAAACCgCC |
| Gdf5_B2 | CCTCgTAAATCCTCATCAAACCTTCTCATGGAACAAGCTGTGGGC |
| Gdf5_B2 | TATTTGCCATTCAACGTTGACCTCAAAATCATCCAgTAAACCgCC |
| Gdf5_B2 | CCTCgTAAATCCTCATCAAAGTCCATTTGGCAGCAACCTCCCAAA |
| Gdf5_B2 | ATGCAGTAGGTAGAATCGTGCCCCTAAATCATCCAgTAAACCgCC |
| Gdf5_B2 | CCTCgTAAATCCTCATCAAACTTCGCAAAAGCACAAGCCAGGACA |
| Gdf5_B2 | AAGAGGCATTGCTAATGCCCCTAGCAAATCATCCAgTAAACCgCC |
| **Crabp2** | |
| Crabp2_B2 | CCTCgTAAATCCTCATCAAATCCAGTTTCCGGTGAAGTTGGGCAT |
| Crabp2_B2 | CCTCGAAGTTCTCCGAGTGCTTCATAAATCATCCAgTAAACCgCC |
| Crabp2_B2 | CCTCgTAAATCCTCATCAAACGTTGACATTCATTGCCTTCAGTAG |
| Crabp2_B2 | CGGCCACGGCAATCTTTCGCAGCATAAATCATCCAgTAAACCgCC |
| Crabp2_B2 | CCTCgTAAATCCTCATCAAATGATCTCCACTGCCGGCTTAGAAGC |
| Crabp2_B2 | TTATGTAGAAATTCTCACCATCTTGAAATCATCCAgTAAACCgCC |
| Crabp2_B2 | CCTCgTAAATCCTCATCAAACGGTAGTGCGGACAGTTGTGAAGGT |
| Crabp2_B2 | ATTCCTCGTTGATTTTGAAGTTGATAAATCATCCAgTAAACCgCC |
| Crabp2_B2 | CCTCgTAAATCCTCATCAAAGCCTGCCATCCACAGTCTGTTCCTC |
| Crabp2_B2 | TCTCCCATTTGACCAGGCTCTTGCAAAATCATCCAgTAAACCgCC |
| Crabp2_B2 | CCTCgTAAATCCTCATCAAAACCAGGAGGTTTTGGGGCCCTCCCC |
| Crabp2_B2 | GCTCCCCGTCGTTGGTCATTTCCCTAAATCATCCAgTAAACCgCC |
| Crabp2_B2 | CCTCgTAAATCCTCATCAAACGTTGTCTGCCTTCATGGTCAGGAT |
| Crabp2_B2 | CCCGCACGTAGATCCTCGTGCAGACAAATCATCCAgTAAACCgCC |
| Crabp2_B2 | CCTCgTAAATCCTCATCAAATTTCTGTGAGTTTCATATGCCTTCA |
| Crabp2_B2 | AAGATGGTTGCTGGAGAGTGTGTTGAAATCATCCAgTAAACCgCC |
| Crabp2_B2 | CCTCgTAAATCCTCATCAAACGTGGGCGTGCAAGGACTGGCAGGA |
| Crabp2_B2 | TTAAGAGGACAGGAATTGTCTGCAGAAATCATCCAgTAAACCgCC |
| Crabp2_B2 | CCTCgTAAATCCTCATCAAATCCAGGCTGTAAACGGGTGGTGTCT |
| Crabp2_B2 | CGGGTCAGAGCAGGCAGGATCGAGTAAATCATCCAgTAAACCgCC |
| Crabp2_B2 | CCTCgTAAATCCTCATCAAAGGGAGATCCACATATGAAGATCAGA |
| Crabp2_B2 | TCAGTCTGTTTCCCTATCGAAGGCGAAATCATCCAgTAAACCgCC |
| Crabp2_B2 | CCTCgTAAATCCTCATCAAACAGCAGTGTCCTGTGTGCCCCGGGG |

| | |
|---|---|
| Crabp2_B2 | ATGGTTAGATTTCGAATTGTGTCAGAAATCATCCAgTAAACCgCC |
| Crabp2_B2 | CCTCgTAAATCCTCATCAAACTCTAATTATGTACCCTTGAAAATC |
| Crabp2_B2 | TCCAACATTTTCTGTACAACATCATAAATCATCCAgTAAACCgCC |
| Crabp2_B2 | CCTCgTAAATCCTCATCAAACAAAGATCTCACAGCTTTAGAGGGG |
| Crabp2_B2 | ACATGTCAGACATGTGAGCAGACCGAAATCATCCAgTAAACCgCC |
| Crabp2_B2 | CCTCgTAAATCCTCATCAAAATGGGGTGAGCTATGACAGAGCCCG |
| Crabp2_B2 | TTCAGGTGGCGGCTGGTGCCCTAAAAAATCATCCAgTAAACCgCC |
| Crabp2_B2 | CCTCgTAAATCCTCATCAAAAAAGGGGCGGAGAAAAGTTAGTCAC |
| Crabp2_B2 | ATTTTGTATTGTGTCATATTGAGGTAAATCATCCAgTAAACCgCC |
| Crabp2_B2 | CCTCgTAAATCCTCATCAAAATGCTTTGACTGTCATGCCAATGAA |
| Crabp2_B2 | GTGAAATCCGAGCAAGGGAATTCATAAATCATCCAgTAAACCgCC |
| Crabp2_B2 | CCTCgTAAATCCTCATCAAAAAGAAGACACACAGATCCACAGACT |
| Crabp2_B2 | CCTGGGCATACAGAATAACACCATAAAATCATCCAgTAAACCgCC |
| Crabp2_B2 | CCTCgTAAATCCTCATCAAAGACCCGTCACTGACAATGTGTGGTC |
| Crabp2_B2 | AGCTCAGAGTGAGGTTGCAAAATGGAAATCATCCAgTAAACCgCC |
| Crabp2_B2 | CCTCgTAAATCCTCATCAAAACAAATAGAACCAGCGAAATGAAAA |
| Crabp2_B2 | TTCCTGCTTCATAGGAGGAGTTTGGAAATCATCCAgTAAACCgCC |
| Crabp2_B2 | CCTCgTAAATCCTCATCAAAGCATTTTCAGTACCTTTTGTGAAAA |
| Crabp2_B2 | CAAAGCTGAATGCTTGACAACATATAAATCATCCAgTAAACCgCC |
| Crabp2_B2 | CCTCgTAAATCCTCATCAAAGGCTAGTCAAGGTACCATATTCAAG |
| Crabp2_B2 | TATTCTATTCGTTCACAAGTGAGTAAAATCATCCAgTAAACCgCC |
| Crabp2_B2 | CCTCgTAAATCCTCATCAAAGGTAATCGCATTCTTCACTTGATTT |
| Crabp2_B2 | AATACTACCTAGCTTGTGAATTTGGAAATCATCCAgTAAACCgCC |
| Crabp2_B2 | CCTCgTAAATCCTCATCAAAAATTGCTCCCCATTCTGGACATGTC |
| Crabp2_B2 | GGAAAAGTGCCCTATTCAATGGGTGAAATCATCCAgTAAACCgCC |
| Crabp2_B2 | CCTCgTAAATCCTCATCAAACAGTTCCAGGGCCTGGCTGTCTTGC |
| Crabp2_B2 | GCTGGACAATGCTGGCCTGATTTTGAAATCATCCAgTAAACCgCC |
| Crabp2_B2 | CCTCgTAAATCCTCATCAAAAGCGGATGGGGAATTACATCCCCTG |
| Crabp2_B2 | TCGGCTATGAGGACTTACTCACTGTAAATCATCCAgTAAACCgCC |
| Crabp2_B2 | CCTCgTAAATCCTCATCAAACCACATACAGGAAATGGGGAACGGC |
| Crabp2_B2 | TTTCATGCCAGTGAAGGCAGACTGAAAATCATCCAgTAAACCgCC |
| Crabp2_B2 | CCTCgTAAATCCTCATCAAATTTAGAATGATGACGTTCAAGAAAA |
| Crabp2_B2 | TTAAGCAACCCAAATAAAATATAGGAAATCATCCAgTAAACCgCC |
| Crabp2_B2 | CCTCgTAAATCCTCATCAAAACATCATGTGCATGCAGCCTCCTGG |
| Crabp2_B2 | TTGAAATGACACCCTTCAATGTGCGAAATCATCCAgTAAACCgCC |
| Crabp2_B2 | CCTCgTAAATCCTCATCAAAAAAAGGCATTCTTAATTATTGTAAT |
| Crabp2_B2 | TGGATTTTCTCCCTGGAAGGTATACAAATCATCCAgTAAACCgCC |
| Crabp2_B2 | CCTCgTAAATCCTCATCAAAACAGGATCTTAATAAGGTAAGGAAG |
| Crabp2_B2 | GACGTGACTGAAATCAAACGCTGAAAAATCATCCAgTAAACCgCC |

| | |
|---|---|
| Crabp2_B2 | CCTCgTAAATCCTCATCAAAGTTAATGACTATCTCACATGAATCT |
| Crabp2_B2 | TATCTCCATAATTGCACCCCAGGGCAAATCATCCAgTAAACCgCC |
| Crabp2_B2 | CCTCgTAAATCCTCATCAAAGGGTTGCTGGGCGCTGACAACAAAA |
| Crabp2_B2 | CTTCCTGTATGGACTGTTTTGTGTTAAATCATCCAgTAAACCgCC |
| Crabp2_B2 | CCTCgTAAATCCTCATCAAATTTGGGTAGTGCTGCTGCTGTCCCC |
| Crabp2_B2 | ATTTAAGGCTTTTAGTTTGAAAACGAAATCATCCAgTAAACCgCC |
| Crabp2_B2 | CCTCgTAAATCCTCATCAAAAGGTTTGGATTGCAAATGTGATGGT |
| Crabp2_B2 | CAAGTAAAACAAAATACAACAGAAAAAATCATCCAgTAAACCgCC |
| Crabp2_B2 | CCTCgTAAATCCTCATCAAAACCACTTGGGTTGGTCAGTTCTATA |
| Crabp2_B2 | ATTATCCGTGTGTTACGGCGCTGTGAAATCATCCAgTAAACCgCC |
| Crabp2_B2 | CCTCgTAAATCCTCATCAAAGAAACCGTGTTTTACCAGCTTTTGG |
| Crabp2_B2 | CTGAGAATGAAGGCCCCAAGCCCGCAAATCATCCAgTAAACCgCC |
| **Sox9** | |
| Sox9_B3 | TCTCCTGCTCCTCGGTCATCTTCATTTCCACTCAACTTTAACCCg |
| Sox9_B3 | gTCCCTgCCTCTATATCTTTAGGGGCTGGGCGCTCCGGACAGGCA |
| Sox9_B3 | GCGAGCCCGCCGAGTCCTCGGACATTTCCACTCAACTTTAACCCg |
| Sox9_B3 | gTCCCTgCCTCTATATCTTTCGTCCGAGCTGGAGCCCGACGGGCA |
| Sox9_B3 | CGTTCTCCAGGGGCCGGGTGTTCTCTTCCACTCAACTTTAACCCg |
| Sox9_B3 | gTCCCTgCCTCTATATCTTTGCTCCTGCAGCTCGCCCTTCGGGAA |
| Sox9_B3 | ACTTGTCCTCCTCGCTCTCCTTCTTTTCCACTCAACTTTAACCCg |
| Sox9_B3 | gTCCCTgCCTCTATATCTTTTGACGGCCTCGCGGATGCAGACGGG |
| Sox9_B3 | TCCAGTCGTAGCCCTTGAGCACCTGTTCCACTCAACTTTAACCCg |
| Sox9_B3 | gTCCCTgCCTCTATATCTTTTCACCCGCACGGGCATGGGCACCAG |
| Sox9_B3 | CGTGCGGCTTGCTCTTGCTGGAGCCTTCCACTCAACTTTAACCCg |
| Sox9_B3 | gTCCCTgCCTCTATATCTTTCCATGAAGGCGTTCATGGGCCGCTT |
| Sox9_B3 | GCTTCCTGCGCGCCGCCTGCGCCCATTCCACTCAACTTTAACCCg |
| Sox9_B3 | gTCCCTgCCTCTATATCTTTTGTGCAGGTGCGGGTACTGGTCGGC |
| Sox9_B3 | TGCCCAGGGTCTTGCTGAGCTCGGCTTCCACTCAACTTTAACCCg |
| Sox9_B3 | gTCCCTgCCTCTATATCTTTCGCCCTCGTTCAGCAGCCTCCAGAG |
| Sox9_B3 | CGGCCTCCTCCACGAAGGGACGTTTTTCCACTCAACTTTAACCCg |
| Sox9_B3 | gTCCCTgCCTCTATATCTTTCTTTCTTGTGCTGCACCCGCAGCCT |
| Sox9_B3 | TGGGCTGGTACTTGTAGTCGGGGTGTTCCACTCAACTTTAACCCg |
| Sox9_B3 | gTCCCTgCCTCTATATCTTTGGCCGTTCTTCACCGACTTCCTCCG |
| Sox9_B3 | GTTCGGCGCCCTCCTCCTGGTCGGCTTCCACTCAACTTTAACCCg |
| Sox9_B3 | gTCCCTgCCTCTATATCTTTAGATGGCGTTGGGCGAGATGTGCGT |
| Sox9_B3 | GCGGCGAGTCGGCCTGTAGCGCCTTTTCCACTCAACTTTAACCCg |
| Sox9_B3 | gTCCCTgCCTCTATATCTTTGCACCTCGCTCATGCCGGAGGAGGA |
| Sox9_B3 | ACTGACCGGAGTGTTCGCCAGGGGATTCCACTCAACTTTAACCCg |
| Sox9_B3 | gTCCCTgCCTCTATATCTTTTGGTGGGAGGGGTTGGTGGCCCCTG |

| | |
|---|---|
| Sox9_B3 | TGCCAGGTTGCACGTCGGTTTTGGGTTCCACTCAACTTTAACCCg |
| Sox9_B3 | gTCCCTgCCTCTATATCTTTGGCGCCCTTCTCTCTTAAGGTCTGG |
| Sox9_B3 | GAGGCTGCCTGCCTTCCTCCTGGAGTTCCACTCAACTTTAACCCg |
| Sox9_B3 | gTCCCTgCCTCTATATCTTTTGTCCACGTCCCCAAAGTCAATATG |
| Sox9_B3 | AGATGACGTCGCTACTTAGCACTGATTCCACTCAACTTTAACCCg |
| Sox9_B3 | gTCCCTgCCTCTATATCTTTGTCGAACTCCTGGATGTCGATGGGG |
| Sox9_B3 | AGGGTGACTATTGGGAGGAAGGTACTTCCACTCAACTTTAACCCg |
| Sox9_B3 | gTCCCTgCCTCTATATCTTTGGCCTGTCCATGGGTGACGGGCACC |
| Sox9_B3 | GTAGCTGCTGGTGTAGGTGCCGGGGTTCCACTCAACTTTAACCCg |
| Sox9_B3 | gTCCCTgCCTCTATATCTTTGCAAGCCAGGCATGGGCGGGACCCC |
| Sox9_B3 | AGGGTGGTCAGCGTGTGCTGCTGCTTTCCACTCAACTTTAACCCg |
| Sox9_B3 | gTCCCTgCCTCTATATCTTTTGCTGGCCCTGCCCCTGCTCGTTGC |
| Sox9_B3 | AGCTGCTCCGTCTTGATGTGTGTCCTTCCACTCAACTTTAACCCg |
| Sox9_B3 | gTCCCTgCCTCTATATCTTTTGCTGCTCGCTGTAGTGTGTGGGGC |
| Sox9_B3 | TAGCTCAGCTGCTGCGGGGAATGTTTTCCACTCAACTTTAACCCg |
| Sox9_B3 | gTCCCTgCCTCTATATCTTTTAGTGCTGCTGCAGGTTGAAAGGGC |
| Sox9_B3 | CGCGTGATGGTGGGGTAGGTGGCGTTTCCACTCAACTTTAACCCg |
| Sox9_B3 | gTCCCTgCCTCTATATCTTTTGGTGGTCGGTGTAGTCGTACTGGG |
| Sox9_B3 | GCGTGGCTGTAGTAGCTGTTAGAGCTTCCACTCAACTTTAACCCg |
| Sox9_B3 | gTCCCTgCCTCTATATCTTTGAGTACAGGCTTGAGCTCTGACCCG |
| Sox9_B3 | TGTGTGGGGTTCATGTAGGAGAAGGTTCCACTCAACTTTAACCCg |
| Sox9_B3 | gTCCCTgCCTCTATATCTTTTCTGCGATAGGGGTGTACATGGGGC |
| Sox9_B3 | TGCGGGATGGAGGGGACGCCCGTCGTTCCACTCAACTTTAACCCg |
| Sox9_B3 | gTCCCTgCCTCTATATCTTTAGACTGGCTGCTCCCAGTGCTGGGG |
| Sox9_B3 | GCCTCTAGGGCCGGGTGAGCTGTGTTTCCACTCAACTTTAACCCg |
| Sox9_B3 | gTCCCTgCCTCTATATCTTTTCTTCAAAGTCTGCAGGGCAGCTTG |
| Sox9_B3 | CGGTAACCAAGATGGCCGACTTAAGTTCCACTCAACTTTAACCCg |
| Sox9_B3 | gTCCCTgCCTCTATATCTTTACAAGCTCTTGGTCCTTCCAACATG |
| Sox9_B3 | GTTCGAGCCCAAGCTGTAGTGCAGATTCCACTCAACTTTAACCCg |
| Sox9_B3 | gTCCCTgCCTCTATATCTTTGGGTTCACTGTCCATCCGTTTCCCC |
| Sox9_B3 | AATGTACAGGGTTTCTGGGCCCCGATTCCACTCAACTTTAACCCg |
| Sox9_B3 | gTCCCTgCCTCTATATCTTTAGTGTTCGGTTCTGAGATGTCCTCT |
| Sox9_B3 | TTTCCAGTCTTTGTGAGAACAGAGGTTCCACTCAACTTTAACCCg |
| Sox9_B3 | gTCCCTgCCTCTATATCTTTCTGTTCCTTAATTACCGTTATAAAA |
| Sox9_B3 | TATTCCTCACAGAGGATTTTCTATTTTCCACTCAACTTTAACCCg |
| Sox9_B3 | gTCCCTgCCTCTATATCTTTTTCCACCCATAGGTTTAAATTGGAA |
| Sox9_B3 | GCACCAGTCGTGCCTTTGTCTGCAGTTCCACTCAACTTTAACCCg |
| Sox9_B3 | gTCCCTgCCTCTATATCTTTGTGTCTGCTGCGTTTCGTGTTCCTC |
| Sox9_B3 | GCATCCCGATTGGCTGGCTTCGAGGTTCCACTCAACTTTAACCCg |

| | |
|---|---|
| Sox9_B3 | gTCCCTgCCTCTATATCTTTCCCCTCAGGTGCAAGCGTGCACCCC |
| Sox9_B3 | AGAATTGTGCAGGCAAAGGCGGCGTTTCCACTCAACTTTAACCCg |
| Sox9_B3 | gTCCCTgCCTCTATATCTTTACACAGTACATACTACAACGTGTGG |
| Sox9_B3 | TCTCCACAATAAAAGTCGATGAAACTTCCACTCAACTTTAACCCg |
| Sox9_B3 | gTCCCTgCCTCTATATCTTTGGAGAAACAGTTTTGCTTTCTGGGG |
| Sox9_B3 | TGCAAGAAAACTATGGGAAACAGTGTTCCACTCAACTTTAACCCg |
| Sox9_B3 | gTCCCTgCCTCTATATCTTTAATGAAGCAAAGCTGAACTGGAAAA |
| Sox9_B3 | TGTATCACTCCACGTCACAAGTTACTTCCACTCAACTTTAACCCg |
| Sox9_B3 | gTCCCTgCCTCTATATCTTTGCACAAACAAACGGCCCACAATTTT |
| **Ihh** | |
| Ihh_B3 | gTCCCTgCCTCTATATCTTTCAGCAGCAGAGCGCAGCCCACGGCC |
| Ihh_B3 | CCCACAGCCGAGCACTCCAGGGGCCTTCCACTCAACTTTAACCCg |
| Ihh_B3 | gTCCCTgCCTCTATATCTTTTAGGAGAGTGGCGTTAGCTTTGGGG |
| Ihh_B3 | TCGGGCACGTTGGGGCTGAACTGCTTTCCACTCAACTTTAACCCg |
| Ihh_B3 | gTCCCTgCCTCTATATCTTTTAACGCCCGCTGGCTCCTAGGGTCT |
| Ihh_B3 | TCCGAGTTGCGGGCTATCTTGCCCTTTCCACTCAACTTTAACCCg |
| Ihh_B3 | gTCCCTgCCTCTATATCTTTTAATTGGGCGTCAGCTCCTTGAAGC |
| Ihh_B3 | TCGTCCTTGAAGATAATGTCCGGGTTTCCACTCAACTTTAACCCg |
| Ihh_B3 | gTCCCTgCCTCTATATCTTTGCTGGGTCATGATCCGGTCCGCCCC |
| Ihh_B3 | CTAAGGAGTTAAGCCTGTCTTTGCATTCCACTCAACTTTAACCCg |
| Ihh_B3 | gTCCCTgCCTCTATATCTTTCGGGCCACTGGTTCATGACCGAGAT |
| Ihh_B3 | ACCCCTCGGTCACCCGAAGTTTCACTTCCACTCAACTTTAACCCg |
| Ihh_B3 | gTCCCTgCCTCTATATCTTTCCTCGGAGTGGTGACCGTCTTCGTC |
| Ihh_B3 | CGGCCCGCCCTTCGTAGTGCAGTGATTCCACTCAACTTTAACCCg |
| Ihh_B3 | gTCCCTgCCTCTATATCTTTGGTCCCGGTCCGATGTGGTGATGTC |
| Ihh_B3 | GCCGGGCCAGCATGCCGTACTTGTTTTCCACTCAACTTTAACCCg |
| Ihh_B3 | gTCCCTgCCTCTATATCTTTCCCAGTCGAACCCGGCTTCCACGGC |
| Ihh_B3 | GAATGTGAGCCTTGGACTCGTAGTATTCCACTCAACTTTAACCCg |
| Ihh_B3 | gTCCCTgCCTCTATATCTTTCTGAGTGTTCGGACTTCACTGAGCA |
| Ihh_B3 | GGAAGCAGCCACCAGTCTTTGCTGCTTCCACTCAACTTTAACCCg |
| Ihh_B3 | gTCCCTgCCTCTATATCTTTTCTCCAGCATTGCCAGGGCTGTGGC |
| Ihh_B3 | ACTCCGGTATGGGAATCTTCTGACCTTCCACTCAACTTTAACCCg |
| Ihh_B3 | gTCCCTgCCTCTATATCTTTTGGACAGCACACGGTCGCCTGGCCG |
| Ihh_B3 | TGTACGTTCTTCGGCCCTCTCCATCTTCCACTCAACTTTAACCCg |
| Ihh_B3 | gTCCCTgCCTCTATATCTTTCTTTGTCCAGGAAGGTTAGGAAATT |
| Ihh_B3 | CATAGAAGTCTTTCACTGCCGAAGGTTCCACTCAACTTTAACCCg |
| Ihh_B3 | gTCCCTgCCTCTATATCTTTGATGGGGTGGATCCTGCGTCTCCAC |
| Ihh_B3 | GCAGGTGGGCTGCAGTGATTGCCAGTTCCACTCAACTTTAACCCg |
| Ihh_B3 | gTCCCTgCCTCTATATCTTTGTACAGTGAAGTTATCCGCCACAAA |

| | |
|---|---|
| Ihh_B3 | CAAACACTGCACTGAAGTCGGTGAGTTCCACTCAACTTTAACCCg |
| Ihh_B3 | gTCCCTgCCTCTATATCTTTTGTACTGTCCTGGCTTCACGTGGCT |
| Ihh_B3 | GCAACCCGAGTACCCCTTCTGAAAGTTCCACTCAACTTTAACCCg |
| Ihh_B3 | gTCCCTgCCTCTATATCTTTTGGTGACAGAAACTACCTTAGCTGG |
| Ihh_B3 | GAGCATAGGCACCTATGTCAGTTCGTTCCACTCAACTTTAACCCg |
| Ihh_B3 | gTCCCTgCCTCTATATCTTTCCAACAACGTGCCATGGCTTGTGAG |
| Ihh_B3 | CAAAGCAGGAGACAACTACATTGTCTTCCACTCAACTTTAACCCg |
| Ihh_B3 | gTCCCTgCCTCTATATCTTTGGCTAGCTGTGCCAGGTGATGTTTT |
| Ihh_B3 | ACCATAGTAAAGCCTCAGTGGCCAGTTCCACTCAACTTTAACCCg |
| Ihh_B3 | gTCCCTgCCTCTATATCTTTCTCTGTATGTGTCTCCAGCCTTCCC |
| Ihh_B3 | AAGGACTTCTGAGTACCAGTGAATGTTCCACTCAACTTTAACCCg |
| Ihh_B3 | gTCCCTgCCTCTATATCTTTGTGGAACTCTTCCTTGGCTAGAAAA |
| Ihh_B3 | GCTCCTCGTTAAGGGCATCCCAAGATTCCACTCAACTTTAACCCg |
| Ihh_B3 | gTCCCTgCCTCTATATCTTTGGTTTCGTTTTCCTTCCGTGGCCCC |
| Ihh_B3 | CTGGGGCTGCTTGGCAGGCTAGTCTTTCCACTCAACTTTAACCCg |
| Ihh_B3 | gTCCCTgCCTCTATATCTTTACATTGCCTTGGCCGACAGCACTTG |
| Ihh_B3 | TGCACGGCGTCCCTACATCTCCGTGTTCCACTCAACTTTAACCCg |
| Ihh_B3 | gTCCCTgCCTCTATATCTTTTGGCGGTGGTCTCTCCTGTACCCTC |
| Ihh_B3 | GCCTTTGAGGATTATTGAAGAGGTGTTCCACTCAACTTTAACCCg |
| Ihh_B3 | gTCCCTgCCTCTATATCTTTCTGTCCTTGGATCTCCTGTTGGATG |
| Ihh_B3 | GCCAGGAAAGCAAGGCAGTGTAGTGTTCCACTCAACTTTAACCCg |
| Ihh_B3 | gTCCCTgCCTCTATATCTTTGAGGCGTCCCACTTGTTTCCATTTT |
| Ihh_B3 | ATTATTTCTCCTCTTATTATATCTTTTCCACTCAACTTTAACCCg |
| Ihh_B3 | gTCCCTgCCTCTATATCTTTGTCAATATTTCACAGTCCATTTGGT |
| Ihh_B3 | TCCCACTTTCTTCCCGCAATACATCTTCCACTCAACTTTAACCCg |
| Ihh_B3 | gTCCCTgCCTCTATATCTTTAAGGCCTTTGGTTTTGAAGGGAGAA |
| Ihh_B3 | GTTTGGTTCTGTGGTCAGAGGTGTGTTCCACTCAACTTTAACCCg |
| Ihh_B3 | gTCCCTgCCTCTATATCTTTTTTACATCAGGAATGTTTCGATGTA |
| Ihh_B3 | TGTCGCATCTCACGCTTGTAGTGCCTTCCACTCAACTTTAACCCg |
| Ihh_B3 | gTCCCTgCCTCTATATCTTTGGTATCCGAGGTGGCCTGATTCAAG |
| Ihh_B3 | AATAAGTCATCCCTCCCAGCTTCTTTTCCACTCAACTTTAACCCg |
| Ihh_B3 | gTCCCTgCCTCTATATCTTTCAACCCATCCCTGAATTAATGCTGG |
| Ihh_B3 | TTAGATTTTCTCCTCTCTCCCTCCCTTCCACTCAACTTTAACCCg |
| Ihh_B3 | gTCCCTgCCTCTATATCTTTGTATTCTGGTCTTGCATAGTTATGA |
| Ihh_B3 | TAACAATGTCTAAGGAGCTTCAAAATTCCACTCAACTTTAACCCg |
| Ihh_B3 | gTCCCTgCCTCTATATCTTTGATTGGCAAAGATCCGGTAAGTTTT |
| Ihh_B3 | GGTTTCCCTCTACCTACACTATATATTCCACTCAACTTTAACCCg |
| Ihh_B3 | gTCCCTgCCTCTATATCTTTTGCATGGACTCACTAGATGGGCCAG |
| Ihh_B3 | TGGGGCCATCTGGGGCAGTAGGCATTTCCACTCAACTTTAACCCg |

| | |
|---|---|
| Ihh_B3 | gTCCCTgCCTCTATATCTTTGACCATCAAGAAGCACAGAGCAAAA |
| Ihh_B3 | CCCCTGGGAATGTCGTGCGGCATAATTCCACTCAACTTTAACCCg |
| Ihh_B3 | gTCCCTgCCTCTATATCTTTAGCTTTGACGATGCAAGGAACTATC |
| Ihh_B3 | GTGACGGAACCTTCACATTCGATGATTCCACTCAACTTTAACCCg |
| Ihh_B3 | gTCCCTgCCTCTATATCTTTAAGATAGCTTGCAAAGGCTCAAGGT |
| Ihh_B3 | TCCCACAATAAGTCAACGTCTCTGGTTCCACTCAACTTTAACCCg |
| **Noggin** | |
| Noggin_B1 | gAggAgggCAgCAAACggAACGGAGTCCGAGGAGCACCAGCAGCG |
| Noggin_B1 | TAGTGCTGGCCGGAGCCCTGCTCGATAgAAgAgTCTTCCTTTACg |
| Noggin_B1 | gAggAgggCAgCAAACggAATCGCTGGGAGAGGGGCGGATGTGCA |
| Noggin_B1 | TCGATCAGGTCCACCAGGGGCAGGTTAgAAgAgTCTTCCTTTACg |
| Noggin_B1 | gAggAgggCAgCAAACggAATTGGGGTCGTACACCGGGTCCGGGT |
| Noggin_B1 | AGCAGAGTCTCGTTCAGGTCCCGCTTAgAAgAgTCTTCCTTTACg |
| Noggin_B1 | gAggAgggCAgCAAACggAATCGAAGTGAGCGCCCAGGAGGCCCC |
| Noggin_B1 | GGCAGCAAGACGGCCATGAACGTGGTAgAAgAgTCTTCCTTTACg |
| Noggin_B1 | gAggAgggCAgCAAACggAACCTGCTGCTGCGGCTGCTGCGCCCC |
| Noggin_B1 | TCTTGCTGAGCCGCTGCCGCTTGCCTAgAAgAgTCTTCCTTTACg |
| Noggin_B1 | gAggAgggCAgCAAACggAAGCCACATCTGCAGCTTCCTGCGCAG |
| Noggin_B1 | GCACGGGGCAGAAGGTCTGCGACCATAgAAgAgTCTTCCTTTACg |
| Noggin_B1 | gAggAgggCAgCAAACggAAGGCTGCCCAGGTCGTTCCACGTGTG |
| Noggin_B1 | CCACCTTGACGTAGCGCGGCCAGAATAgAAgAgTCTTCCTTTACg |
| Noggin_B1 | gAggAgggCAgCAAACggAAAGCACGAGCGCTTGCTGTAGCAGCT |
| Noggin_B1 | CCTTGCACACCATGCCCTCGGGCACTAgAAgAgTCTTCCTTTACg |
| Noggin_B1 | gAggAgggCAgCAAACggAAGGATGGTCAGGTGCACCGACTTGGC |
| Noggin_B1 | TCTGCCGCCGCTGGCACCTCCAGCGTAgAAgAgTCTTCCTTTACg |
| Noggin_B1 | gAggAgggCAgCAAACggAAGGTAGTGGATGGGGATCCAGTGGCA |
| Noggin_B1 | AGGAGCACTTGCACTCCGAGATGATTAgAAgAgTCTTCCTTTACg |
| Noggin_B1 | gAggAgggCAgCAAACggAACCAGGCGGGAAGGCGCCCGGCCTCA |
| Noggin_B1 | TAGTCTCTGGTGTGGGCGGCACCGGTAgAAgAgTCTTCCTTTACg |
| Noggin_B1 | gAggAgggCAgCAAACggAACTGTGCCACCCGCCTAGGCCCTCAG |
| Noggin_B1 | AGGAAGTCCGTTCCGTTCTTAAGAATAgAAgAgTCTTCCTTTACg |
| Noggin_B1 | gAggAgggCAgCAAACggAACTGCTGCTCAGAGCCGGTTTTGCAG |
| Noggin_B1 | CAATGTACATTCCTACAACACTGGGTAgAAgAgTCTTCCTTTACg |
| Noggin_B1 | gAggAgggCAgCAAACggAATTAAACTTGGCACCATAAATACACA |
| Noggin_B1 | GAATAATACTGACCTTTTAATAGCATAgAAgAgTCTTCCTTTACg |
| Noggin_B1 | gAggAgggCAgCAAACggAAAAGTACAGTAGGCGCTGGTTATTTA |
| Noggin_B1 | AGCATCAACCAGGATTAAAGGTGTCTAgAAgAgTCTTCCTTTACg |
| **Ptch1** | |
| Ptch1_B3 | gTCCCTgCCTCTATATCTTTCGCTGCAAGTACTCCGGGTCAGGGG |

| | |
|---|---|
| Ptch1_B3 | GCAAACGCGGCGTCGCAGTAGCTCGTTCCACTCAACTTTAACCCg |
| Ptch1_B3 | gTCCCTgCCTCTATATCTTTGCCTTGCCCTTGTAGATCTGCTCCA |
| Ptch1_B3 | AGCCAGAGGGGCGCTTTCCGGCCCGTTCCACTCAACTTTAACCCg |
| Ptch1_B3 | gTCCCTgCCTCTATATCTTTAAGAGCAATCTCTGGAATTTCGCTC |
| Ptch1_B3 | TTCCTCTGTATGTAGCATCCCAGTCTTCCACTCAACTTTAACCCg |
| Ptch1_B3 | gTCCCTgCCTCTATATCTTTAGGCCCACCACCAAGAATTTTCCGC |
| Ptch1_B3 | CCCACGGCGAAAGCGCCGAAGATAATTCCACTCAACTTTAACCCg |
| Ptch1_B3 | gTCCCTgCCTCTATATCTTTTTAGTCTCCAAGTTGGCGGCTCGGA |
| Ptch1_B3 | CCAACTTCTACCCAGAGCTCCTCCATTCCACTCAACTTTAACCCg |
| Ptch1_B3 | gTCCCTgCCTCTATATCTTTTAATTTAATTCCCGACTCACTCGTC |
| Ptch1_B3 | GCCTCTTCTCCAATCTTCTGGCGCGTTCCACTCAACTTTAACCCg |
| Ptch1_B3 | gTCCCTgCCTCTATATCTTTTGGATCATGAGTTGGGGATTAAACA |
| Ptch1_B3 | ACATTAGCACCATCTTCTCTAGGTGTTCCACTCAACTTTAACCCg |
| Ptch1_B3 | gTCCCTgCCTCTATATCTTTTGCTGCCGCAGCGCATCAGCCGTGA |
| Ptch1_B3 | CTGCTGGCATCCAGTGCAGATTGGATTCCACTCAACTTTAACCCg |
| Ptch1_B3 | gTCCCTgCCTCTATATCTTTTGTCTGTTGTACATGTAGACTTGGA |
| Ptch1_B3 | TTATAGCATAACTGTTCCAATTTCCTTCCACTCAACTTTAACCCg |
| Ptch1_B3 | gTCCCTgCCTCTATATCTTTCCAGCTTCTGTGATCAGTTCTCCCG |
| Ptch1_B3 | AGAGATTCAATAATGTGGTTTACATTTCCACTCAACTTTAACCCg |
| Ptch1_B3 | gTCCCTgCCTCTATATCTTTAAAGGTGTGATAACCAAACACGGAT |
| Ptch1_B3 | AGTTTAGCTCCCTCCCAGAAACAGTTTCCACTCAACTTTAACCCg |
| Ptch1_B3 | gTCCCTgCCTCTATATCTTTCCTGGGAGATACGCTGTGCCCGTTT |
| Ptch1_B3 | TGTGTCCACTGCAAGGGAAGATGCTTTCCACTCAACTTTAACCCg |
| Ptch1_B3 | gTCCCTgCCTCTATATCTTTAGCTGTCCGTCTGATAGAATATTTT |
| Ptch1_B3 | CCTCAGCCTTGTTCAGCATTTCCTCTTCCACTCAACTTTAACCCg |
| Ptch1_B3 | gTCCCTgCCTCTATATCTTTAAGGCCGGTCCATATAACCCTGGCC |
| Ptch1_B3 | GACATTCGGGGTCTGCGGGATTCAATTCCACTCAACTTTAACCCg |
| Ptch1_B3 | gTCCCTgCCTCTATATCTTTACGTCAAGAGGTCTGGTGGAGTTTT |
| Ptch1_B3 | TAGCATCCACCGCTCAAAATAAGTGTTCCACTCAACTTTAACCCg |
| Ptch1_B3 | gTCCCTgCCTCTATATCTTTTCTTCCTGCCAGCGCATGTACTTTT |
| Ptch1_B3 | TGCTTGACTGTACCACCCACAATCATTCCACTCAACTTTAACCCg |
| Ptch1_B3 | gTCCCTgCCTCTATATCTTTTGGGCGCTGAGAAGCGTTCCATTGC |
| Ptch1_B3 | ATTAACTGGAACATGGTCTGCAAAGTTCCACTCAACTTTAACCCg |
| Ptch1_B3 | gTCCCTgCCTCTATATCTTTAAGTGCTCGTACATTTGCTTGGGAG |
| Ptch1_B3 | ATGTTCACAACGTCCTCATATCCCTTTCCACTCAACTTTAACCCg |
| Ptch1_B3 | gTCCCTgCCTCTATATCTTTGCAGCAGCCTTGTCCTCATTCCAGT |
| Ptch1_B3 | TACATCCTCTGCCAGGCCTCAAGAATTCCACTCAACTTTAACCCg |
| Ptch1_B3 | gTCCCTgCCTCTATATCTTTGCTACACTTTGATGAACAACCTCAA |
| Ptch1_B3 | GAAACCACCTTCTGAGTAGAGTTTGTTCCACTCAACTTTAACCCg |

| | |
|---|---|
| Ptch1_B3 | gTCCCTgCCTCTATATCTTTATGTCGTCCAAAGTAGTCGTGGTGA |
| Ptch1_B3 | ACACTAATATCAGAAAACGATTTGATTCCACTCAACTTTAACCCg |
| Ptch1_B3 | gTCCCTgCCTCTATATCTTTAATAAGTAGCCGCTGGCCACTCGGA |
| Ptch1_B3 | ATGGTTAAACAAGCATAGGCAAGCATTCCACTCAACTTTAACCCg |
| Ptch1_B3 | gTCCCTgCCTCTATATCTTTTGGGACTTGGCGCAGTCCCACCGCA |
| Ptch1_B3 | AGCACTCCTGCAAGCCCCACGGCACTTCCACTCAACTTTAACCCg |
| Ptch1_B3 | gTCCCTgCCTCTATATCTTTCCTGCAGCCACTGAGAGCGCAACCA |
| Ptch1_B3 | ATTCCAATCAATGAACACAGGCCCATTCCACTCAACTTTAACCCg |
| Ptch1_B3 | gTCCCTgCCTCTATATCTTTACCTGAGTAGTGGCAGCGTTAAAGG |
| Ptch1_B3 | CCAACGCCAAGCGCAAGGAACGGCATTCCACTCAACTTTAACCCg |
| Ptch1_B3 | gTCCCTgCCTCTATATCTTTTGTGCCAGAAGGAAAACGTCGTCCA |
| Ptch1_B3 | TTATTCTGCCCAGTCTCACTGAATGTTCCACTCAACTTTAACCCg |
| Ptch1_B3 | gTCCCTgCCTCTATATCTTTTGGAGGTCAGGGCCACACTGGCCCC |
| Ptch1_B3 | CCATAAAGAAGGCAGTGACATTGCTTTCCACTCAACTTTAACCCg |
| Ptch1_B3 | gTCCCTgCCTCTATATCTTTGGAGGGCTGGGATGGGGATAAGGGC |
| Ptch1_B3 | CAACCGCGGCTTGGAGCGAAAAGGCTTCCACTCAACTTTAACCCg |
| Ptch1_B3 | gTCCCTgCCTCTATATCTTTAGACCATGGCAAAGTTGAAGACCAC |
| Ptch1_B3 | TGCTAAGGATGGCTGGAAAGATGAGTTCCACTCAACTTTAACCCg |
| Ptch1_B3 | gTCCCTgCCTCTATATCTTTTGCAATCTTCGCGCCGGTAGAGATC |
| Ptch1_B3 | GTGTGAAGCAGCAGAAAATATCAAGTTCCACTCAACTTTAACCCg |
| Ptch1_B3 | gTCCCTgCCTCTATATCTTTCCTGGATCACTCGACTGGCACAGGG |
| Ptch1_B3 | TACGGTCTCCATAGGCCTGAGGTTCTTCCACTCAACTTTAACCCg |
| Ptch1_B3 | gTCCCTgCCTCTATATCTTTGTGGAGGGCTGTATTGTGAACGCTC |
| Ptch1_B3 | GTGCAAAGCTGTGACTGCTGTAGGGTTCCACTCAACTTTAACCCg |
| Ptch1_B3 | gTCCCTgCCTCTATATCTTTTGGACTGCATTGTGATTTGGGTTTC |
| Ptch1_B3 | GATCATACTCCGTGCGCAGCTGTACTTCCACTCAACTTTAACCCg |
| Ptch1_B3 | gTCCCTgCCTCTATATCTTTCAGTGGTGTAGTAAGCTTGTGTGCG |
| Ptch1_B3 | GCACAGATATTTCGGAGCGAGGTTCTTCCACTCAACTTTAACCCg |
| Ptch1_B3 | gTCCCTgCCTCTATATCTTTTCTCCTGTGGCACTGCAACTGCCGG |
| Ptch1_B3 | AGCTCGTGCTTTCGGAACAGTTCAGTTCCACTCAACTTTAACCCg |
| Ptch1_B3 | gTCCCTgCCTCTATATCTTTAGATATGGGAGAGTAAATCCCTTGT |
| Ptch1_B3 | GCTCAAGGCAACGCATTCCAGAGTCTTCCACTCAACTTTAACCCg |
| Ptch1_B3 | gTCCCTgCCTCTATATCTTTACGACAGTGTCCACTTTGTACATGG |
| Ptch1_B3 | ACGGGGCGTAATGTTTCTCAGCAAATTCCACTCAACTTTAACCCg |
| **Alx4** | |
| Alx4_B2 | CCTCgTAAATCCTCATCAAATTCATCCCTGAAAGATGAGGAGGGG |
| Alx4_B2 | CAGTACGAGGACACGCACGTGTCCGAAATCATCCAgTAAACCgCC |
| Alx4_B2 | CCTCgTAAATCCTCATCAAAGGGCTGTAGTAAGGGTCCATGCCCC |
| Alx4_B2 | GGGTGGTCCCTGCCCTGCTGGGCGGAAATCATCCAgTAAACCgCC |

| | |
|---|---|
| Alx4_B2 | CCTCgTAAATCCTCATCAAAGCCTGGAAGGCCGTATAAGGAGGCG |
| Alx4_B2 | AAGGCTGGGCCGAACTTGTCACCTGAAATCATCCAgTAAACCgCC |
| Alx4_B2 | CCTCgTAAATCCTCATCAAACCATAGCCTGTCCCTTTGCTGGCCA |
| Alx4_B2 | TGCGAGTAGGAAGCCCGCGACCTGTAAATCATCCAgTAAACCgCC |
| Alx4_B2 | CCTCgTAAATCCTCATCAAAGCGGTCCTCTCCTCCCCGGGGCTGT |
| Alx4_B2 | AGGTATTGCTGCTGGTACTTGCTGTAAATCATCCAgTAAACCgCC |
| Alx4_B2 | CCTCgTAAATCCTCATCAAAGGCGGGGTCTTGCAGGAGCCCCTCT |
| Alx4_B2 | CTGCCCTCGTGCAGCTTCGGGTTCTAAATCATCCAgTAAACCgCC |
| Alx4_B2 | CCTCgTAAATCCTCATCAAATAGCAGGGGCCCAGGCCGTCGCCGT |
| Alx4_B2 | TCTGATTCCAATGAGTTCTCTTTAGAAATCATCCAgTAAACCgCC |
| Alx4_B2 | CCTCgTAAATCCTCATCAAACCTGAAAGGTCACTACTCGGGGTCA |
| Alx4_B2 | TTCACAGTGAGGTAACTATTGTCCAAAATCATCCAgTAAACCgCC |
| Alx4_B2 | CCTCgTAAATCCTCATCAAATCCTGAGACCCCTTCACACCAGACT |
| Alx4_B2 | GGGCTTGGTAGGTCACCGCTGCTCCAAATCATCCAgTAAACCgCC |
| Alx4_B2 | CCTCgTAAATCCTCATCAAATTACTTTCTGAATCGCCCTTCTCCA |
| Alx4_B2 | GTCCTATTTCTCCTCTTCTTGCCCTAAATCATCCAgTAAACCgCC |
| Alx4_B2 | CCTCgTAAATCCTCATCAAATCCTCTAGTTGGTAACTGGTGAATG |
| Alx4_B2 | TGTGTTTTCTGGAAGACCTTCTCCAAAATCATCCAgTAAACCgCC |
| Alx4_B2 | CCTCgTAAATCCTCATCAAATGCTCCCGGGCGTAGACATCAGGAT |
| Alx4_B2 | TCCGTGAGGTCAGTGCGCATAGCGAAAATCATCCAgTAAACCgCC |
| Alx4_B2 | CCTCgTAAATCCTCATCAAATTCTGGAACCACACCTGGACCCGTG |
| Alx4_B2 | TCTCGCTTCCTCCACTTCGCACGTCAAATCATCCAgTAAACCgCC |
| Alx4_B2 | CCTCgTAAATCCTCATCAAACTCACTTGCTGCATCTGACCAAAGC |
| Alx4_B2 | AGCTCGTAGGCGCTTGAAAAGTGTGAAATCATCCAgTAAACCgCC |
| Alx4_B2 | CCTCgTAAATCCTCATCAAATAATTCTCAGCCCGGGTTAGGAGCG |
| Alx4_B2 | ATCCATGTTGGGTTCTGGATCTGTGAAATCATCCAgTAAACCgCC |
| Alx4_B2 | CCTCgTAAATCCTCATCAAAACACAGGCTGCCACCGGAGAGCCCC |
| Alx4_B2 | CAGGATGGGACGGTGTCACAGGGGAAAATCATCCAgTAAACCgCC |
| Alx4_B2 | CCTCgTAAATCCTCATCAAACAGAAACACCTCCACTTGCATGGGG |
| Alx4_B2 | TTCCTGAGCTAGGCACGCTCAGGAAAAATCATCCAgTAAACCgCC |
| Alx4_B2 | CCTCgTAAATCCTCATCAAATACCCATGTGTGCCTGACCCATGTG |
| Alx4_B2 | TGCTGAGCCCAGCAGTGCCAAAGAGAAATCATCCAgTAAACCgCC |
| Alx4_B2 | CCTCgTAAATCCTCATCAAACGTTGAGGTCGTAGCCATTGATGCC |
| Alx4_B2 | TGCTGGAGGTCTTGCGGTCTGGCTCAAATCATCCAgTAAACCgCC |
| Alx4_B2 | CCTCgTAAATCCTCATCAAACCTTGGCCTTCATCCGCAGGGCTGC |
| Alx4_B2 | TGGCCCAGGAGATGGCGGCACTGTGAAATCATCCAgTAAACCgCC |
| Alx4_B2 | CCTCgTAAATCCTCATCAAATGCATGCTCTTTAAGTGCCCTGTCA |
| Alx4_B2 | TATCCAATCTCATGCTATGTCAGTGAAATCATCCAgTAAACCgCC |
| Alx4_B2 | CCTCgTAAATCCTCATCAAAGATGTCTGTTGGCCAATGGGTTGGC |

| | |
|---|---|
| Alx4_B2 | CAGCACACACATTCTAAGGTTGTGTAAATCATCCAgTAAACCgCC |
| Alx4_B2 | CCTCgTAAATCCTCATCAAATCTTAGAACTATTTCTTGCTAATAG |
| Alx4_B2 | ACGCACGCAGTTATGGTTATGTCAAAAATCATCCAgTAAACCgCC |
| Alx4_B2 | CCTCgTAAATCCTCATCAAATACTGCATGTGGGAACCTTTCCTCT |
| Alx4_B2 | ACTTGGTTAGCTGGCTGTGCCGAAGAAATCATCCAgTAAACCgCC |
| Alx4_B2 | CCTCgTAAATCCTCATCAAATGATGGTTTTCTCATTACAAATCCA |
| Alx4_B2 | AAATAGTCTACTTGGAATTCACAAGAAATCATCCAgTAAACCgCC |
| Alx4_B2 | CCTCgTAAATCCTCATCAAATCATCAGCAAATGGAACAAACAAAA |
| Alx4_B2 | GGTGTCTGCCAAGAATAAAATCCCAAAATCATCCAgTAAACCgCC |
| Alx4_B2 | CCTCgTAAATCCTCATCAAAAATTTTAAGCACTGAACTGTTCAAT |
| Alx4_B2 | CTCTGAACTGGCTTTGAGTATTAAAAAATCATCCAgTAAACCgCC |
| Alx4_B2 | CCTCgTAAATCCTCATCAAATTCAGCTGTGGTCACTCACATGCAC |
| Alx4_B2 | CTCCAATGTGGTCACTGCCCTCTGTAAATCATCCAgTAAACCgCC |
| Alx4_B2 | CCTCgTAAATCCTCATCAAATAGCTTGTATCAAGAGCTCTTCAAC |
| Alx4_B2 | GGAAGATTTAATGTGGGTATTATGAAAATCATCCAgTAAACCgCC |
| Alx4_B2 | CCTCgTAAATCCTCATCAAAAGCTGTGCTTTCTTAAGGTGGCAAA |
| Alx4_B2 | AGTGAGACTGCATTTGAGTGCTTCCAAATCATCCAgTAAACCgCC |
| Alx4_B2 | CCTCgTAAATCCTCATCAAAATTGAGTCTGTTATGTGGACTATTT |
| Alx4_B2 | TGTTGTTGGACAAGTCCACCATAGTAAATCATCCAgTAAACCgCC |
| Alx4_B2 | CCTCgTAAATCCTCATCAAATGACACGAGGGAGAGAAGTGTTATG |
| Alx4_B2 | GGAGATGGCTTCCAGGTGGGATATGAAATCATCCAgTAAACCgCC |
| Alx4_B2 | CCTCgTAAATCCTCATCAAAAAATCCAACACATGCTGCATGGATC |
| Alx4_B2 | CATTTAAGGGAATTGGCATCCCTGGAAATCATCCAgTAAACCgCC |
| Alx4_B2 | CCTCgTAAATCCTCATCAAACAAAAGGTTCTTGAGATCCTGCTAG |
| Alx4_B2 | TATCTTGCAAAGTTTCTAATCTAATAAATCATCCAgTAAACCgCC |
| Alx4_B2 | CCTCgTAAATCCTCATCAAATCACCTTGAAACAGCACATGCTGAA |
| Alx4_B2 | AAAGGAATTTTAGATGTCCTGCATTAAATCATCCAgTAAACCgCC |
| Alx4_B2 | CCTCgTAAATCCTCATCAAATTGGTTTCTTCATGACCAATCAGAA |
| Alx4_B2 | AAGTTTGATAAACCGAGTTTGGCAGAAATCATCCAgTAAACCgCC |
| Alx4_B2 | CCTCgTAAATCCTCATCAAAAATGAGAAGAGTTGGCAATGGCCAA |
| Alx4_B2 | ATCAGAAAGGACATTGGTACAGATAAAATCATCCAgTAAACCgCC |
| **Smo** | |
| Smo_B1 | gAggAgggCAgCAAACggAATACAAGGTGGTACAAAGACCTTTCA |
| Smo_B1 | CATTCGTTGCCCCATCCTCAAAGACTAgAAgAgTCTTCCTTTACg |
| Smo_B1 | gAggAgggCAgCAAACggAACTTATTGGAGGGTGTAGACAACCCC |
| Smo_B1 | GCGGCTGGACGTTTTGCCACGCTGTTAgAAgAgTCTTCCTTTACg |
| Smo_B1 | gAggAgggCAgCAAACggAAACAAGGTCTGAGCAACCCTCAGCAT |
| Smo_B1 | TCCAGCCGCCTTAACTTGCTCTCCATAgAAgAgTCTTCCTTTACg |
| Smo_B1 | gAggAgggCAgCAAACggAACTTGCCGCATCGCAAACCGGAACAG |

| | |
|---|---|
| Smo_B1 | CCAAATACAACTGTTTCTAGATGGTTAgAAgAgTCTTCCTTTACg |
| Smo_B1 | gAggAgggCAgCAAACggAAATCCGTGCAGGAGAGTCTGCTGAGC |
| Smo_B1 | ACCAGAGACATCTTGGAACGCCCCATAgAAgAgTCTTCCTTTACg |
| Smo_B1 | gAggAgggCAgCAAACggAACAGAGCCACTGAATCAGGCCTTGCT |
| Smo_B1 | AGTAGAAGCCACAGTACCAACAACATAgAAgAgTCTTCCTTTACg |
| Smo_B1 | gAggAgggCAgCAAACggAAAGCGCCTCGCAGGTGGCTGTCTTTT |
| Smo_B1 | GTGGACCCCAGGCAGACATTGTACTTAgAAgAgTCTTCCTTTACg |
| Smo_B1 | gAggAgggCAgCAAACggAAGCAATGGAGGTGAGGCCATAGGGCA |
| Smo_B1 | TCCTGAGAGCTTGAGTCTTCGGCCATAgAAgAgTCTTCCTTTACg |
| Smo_B1 | gAggAgggCAgCAAACggAACAGAGGAGCAGCTTCTGGTTGGCAT |
| Smo_B1 | CAGCGGGGAGCATTGCGGAGACCTGTAgAAgAgTCTTCCTTTACg |
| Smo_B1 | gAggAgggCAgCAAACggAACACAGGAGGGGCTGAATAACATCCC |
| Smo_B1 | CTTTCGCACTTTGGCATGTAGACAGTAgAAgAgTCTTCCTTTACg |
| Smo_B1 | gAggAgggCAgCAAACggAACCTTGGCTGGGGAGCTCCACCTTTC |
| Smo_B1 | CACGGGCCCCTTGTGGCCTGGCACATAgAAgAgTCTTCCTTTACg |
| Smo_B1 | gAggAgggCAgCAAACggAACATCCTCGTTCACGCTCTACGATGG |
| Smo_B1 | GGGTTACTGCACTTCAGGAAGTCTGTAgAAgAgTCTTCCTTTACg |
| Smo_B1 | gAggAgggCAgCAAACggAATCATTCATGCACCCTTCTGGAAAGC |
| Smo_B1 | GAGCTGTTGAACTTGATGTTCTGGATAgAAgAgTCTTCCTTTACg |
| Smo_B1 | gAggAgggCAgCAAACggAACGCACGAGGGGCGGCTCGCACTGGC |
| Smo_B1 | TCGTACCAGCTCTTGGGGTTGTCTGTAgAAgAgTCTTCCTTTACg |
| Smo_B1 | gAggAgggCAgCAAACggAACACTGGATCCCGCAGCCCTCCACGT |
| Smo_B1 | TCCCGCTCCGTGAACAAGGGGTTCTTAgAAgAgTCTTCCTTTACg |
| Smo_B1 | gAggAgggCAgCAAACggAAGCGATATAGCCATGCATCTCCCGGT |
| Smo_B1 | CAGAAGACGGTGACGGAGCTGAAGGTAgAAgAgTCTTCCTTTACg |
| Smo_B1 | gAggAgggCAgCAAACggAAACTGCCGGGTAGCGGTTGGAGTTTT |
| Smo_B1 | AAGCAGGCGTTGACGTAGAAGAGGATAgAAgAgTCTTCCTTTACg |
| Smo_B1 | gAggAgggCAgCAAACggAAGCCAGCCAGCCAATGCTGCCCACGA |
| Smo_B1 | TCCAATCGTGCCCCATCCATGAACTTAgAAgAgTCTTCCTTTACg |
| Smo_B1 | gAggAgggCAgCAAACggAAATGGTCCCGTCTGCCTTGCAGACTA |
| Smo_B1 | TCGTTCGAGGTCGGCTCTCCCAGCCTAgAAgAgTCTTCCTTTACg |
| Smo_B1 | gAggAgggCAgCAAACggAAACGAAGATGATCACGCAGGACAGTG |
| Smo_B1 | CCAGACATCATGGAATAATAGACAATAgAAgAgTCTTCCTTTACg |
| Smo_B1 | gAggAgggCAgCAAACggAATAGGTCAGCATAACAAACCAGATGA |
| Smo_B1 | AGTGCCCTGAACGAGGCGTGCCAGGTAgAAgAgTCTTCCTTTACg |
| Smo_B1 | gAggAgggCAgCAAACggAACCTGATAGTGGCTGGTATGTGGTCC |
| Smo_B1 | GTAACCAGGTGGAAATAGGAGGTCCTAgAAgAgTCTTCCTTTACg |
| Smo_B1 | gAggAgggCAgCAAACggAAACCGTGAGGACAAAGGGGATAGACC |
| Smo_B1 | TCCACCTGAGCAACCGCCAAAATGGTAgAAgAgTCTTCCTTTACg |

| | |
|---|---|
| Smo_B1 | gAggAgggCAgCAAACggAAAAGCAAATTCCACTGACTGAATCCC |
| Smo_B1 | CGGTAACGGTAGTTCTTGTATCCCATAgAAgAgTCTTCCTTTACg |
| Smo_B1 | gAggAgggCAgCAAACggAACCGATGGGTGCCAGGACAAAGCCTG |
| Smo_B1 | AAATAACCTCCAACAATCAGCACCATAgAAgAgTCTTCCTTTACg |
| Smo_B1 | gAggAgggCAgCAAACggAAGAAGCCAAAGGCCAGGAATCCAAAA |
| Smo_B1 | GAAGTGGCAGCTGAACGTGATCAGATAgAAgAgTCTTCCTTTACg |
| Smo_B1 | gAggAgggCAgCAAACggAACCATTCCGCCTGGTGGAAGAAGTCA |
| Smo_B1 | GAGAACATATTCTCGGAAGCTGCGCTAgAAgAgTCTTCCTTTACg |
| Smo_B1 | gAggAgggCAgCAAACggAACAGCGCAATGGTTACATTGGCCTCA |
| Smo_B1 | GCAGTCTGGGACTGGCTTGTTGGCCTAgAAgAgTCTTCCTTTACg |
| Smo_B1 | gAggAgggCAgCAAACggAAGATGCCAGTTCCAAACATCGCAAAA |
| Smo_B1 | CTTGGTCCACACCCAGGTGCTCATGTAgAAgAgTCTTCCTTTACg |
| Smo_B1 | gAggAgggCAgCAAACggAAGGTGCGCTTCCAGATAAGTACAGTG |
| Smo_B1 | GTCGCTGCGCCCAATGATCCGGCACTAgAAgAgTCTTCCTTTACg |
| Smo_B1 | gAggAgggCAgCAAACggAAGCCTTGGCGATCATCTTGCTCTTTT |
| Smo_B1 | TTCAGCAGCTCCTTCCTCTTAGAGATAgAAgAgTCTTCCTTTACg |
| Smo_B1 | gAggAgggCAgCAAACggAATGCTGAAGGACAGCTCGCGTTGGGG |
| Smo_B1 | CAGGCCCCTCGTGCGACACTGTGTGTAgAAgAgTCTTCCTTTACg |
| Smo_B1 | gAggAgggCAgCAAACggAACATTCAGGTCGATGTTCAGACCAGC |
| Smo_B1 | ACGCAGAAGACACGTCAGCAGAGGGTAgAAgAgTCTTCCTTTACg |
| Smo_B1 | gAggAgggCAgCAAACggAACCACCATCTTGGTAACATGCTGGGC |
| Smo_B1 | CCTGGGGTAGGATAGCGCCCCTCCTTAgAAgAgTCTTCCTTTACg |
| Smo_B1 | gAggAgggCAgCAAACggAAGTGTTGCTACTGGTGTGAGGGACAG |
| Smo_B1 | ATTTGGCCCTTTCCTCCGGTGGTACTAgAAgAgTCTTCCTTTACg |
| Smo_B1 | gAggAgggCAgCAAACggAAGTGGAAGGCTGGCGTCCACCAGCCA |
| Smo_B1 | TCTTGAATCCATGCTTATCAAGCTCTAgAAgAgTCTTCCTTTACg |
| Smo_B1 | gAggAgggCAgCAAACggAACCATCTCATCCGTAGCTCCAAGGGG |
| Smo_B1 | CCCGGGCTACCCTGTAAGCCCCATCTAgAAgAgTCTTCCTTTACg |
| **Apoeb** | |
| Apoeb_B1 | gAggAgggCAgCAAACggAACTCTGTCTCCTCCAGTGCTTGCTGT |
| Apoeb_B1 | TCTCCAAGGAAACGCGCGCCTGGATTAgAAgAgTCTTCCTTTACg |
| Apoeb_B1 | gAggAgggCAgCAAACggAATTCATGGTGATGTCGTATCTTCCGC |
| Apoeb_B1 | ACAAGAGCCAGCACCAGGCCCAGGATAgAAgAgTCTTCCTTTACg |
| Apoeb_B1 | gAggAgggCAgCAAACggAAAGGTTGGCTTGACACCCTGTCAGCG |
| Apoeb_B1 | TTGGTCTGCGGCTCATCGTTGAAGATAgAAgAgTCTTCCTTTACg |
| Apoeb_B1 | gAggAgggCAgCAAACggAACAGAAGGTGTCCACTGCCGTTTCCC |
| Apoeb_B1 | ACCTGCCCCACTTGGGACACGTAATTAgAAgAgTCTTCCTTTACg |
| Apoeb_B1 | gAggAgggCAgCAAACggAACTGACCTTGGCCGTGGCGTCATCTG |
| Apoeb_B1 | TCCAGTTCTTTGCTGATCTGCGACTTAgAAgAgTCTTCCTTTACg |

| | |
|---|---|
| Apoeb_B1 | gAggAgggCAgCAAACggAATCGTCCATGGTGTCGCTGATAAGAC |
| Apoeb_B1 | TTGATGCCTTCTGAGTAGGTATTGATAgAAgAgTCTTCCTTTACg |
| Apoeb_B1 | gAggAgggCAgCAAACggAATCTTGAGTGTAAGGGCCCAGCTTGG |
| Apoeb_B1 | ACTTCAGTGCTAAACCTCCGCTGCGTAgAAgAgTCTTCCTTTACg |
| Apoeb_B1 | gAggAgggCAgCAAACggAAGTCTTCAGCTTCTCAGAGATTGCTG |
| Apoeb_B1 | ACTTTGCTTTTGGTGTCCTCCATGTTAgAAgAgTCTTCCTTTACg |
| Apoeb_B1 | gAggAgggCAgCAAACggAAGGTCGAACATCATGCGGATGTCCCC |
| Apoeb_B1 | CTCGGGTGCGGACCTCCTCGAGGTTTAgAAgAgTCTTCCTTTACg |
| Apoeb_B1 | gAggAgggCAgCAAACggAATCTTCATCTTCCTCAGGTACATGCT |
| Apoeb_B1 | GCTCCTCTGTGTCCTTGTTGAGCCTTAgAAgAgTCTTCCTTTACg |
| Apoeb_B1 | gAggAgggCAgCAAACggAACGGCATAGGTGCTCATCTTCCTCTT |
| Apoeb_B1 | TCTGGTTCGTTTGGCTGCGCACCTCTAgAAgAgTCTTCCTTTACg |
| Apoeb_B1 | gAggAgggCAgCAAACggAACTACTGAGTCGCGGATGGCCTCCGC |
| Apoeb_B1 | TGTCCCGGACATTGCTGATAATGGGTAgAAgAgTCTTCCTTTACg |
| Apoeb_B1 | gAggAgggCAgCAAACggAATAAGGGCCTGAAGGCGCTGCTGGCT |
| Apoeb_B1 | TCTTGCCCTGCTCGTCCATCGCCTGTAgAAgAgTCTTCCTTTACg |
| Apoeb_B1 | gAggAgggCAgCAAACggAACTCTGGAGTTCAGCTGGTCGCGCAC |
| Apoeb_B1 | CCTTCACCTTGTCCTGGATCTCCCCTAgAAgAgTCTTCCTTTACg |
| Apoeb_B1 | gAggAgggCAgCAAACggAACAGAGTTCCGCACGATGTCCATCTT |
| Apoeb_B1 | TTCTCATCATCTCGGTGTTCTCCTCTAgAAgAgTCTTCCTTTACg |
| Apoeb_B1 | gAggAgggCAgCAAACggAATGTTCTCCAAGTAGGGTGCAAACCA |
| Apoeb_B1 | CCAGGAAGGACTCAAACTGGTTGCGTAgAAgAgTCTTCCTTTACg |
| Apoeb_B1 | gAggAgggCAgCAAACggAATTTACTGCTTCTTCCCCTGCAGGGT |
| Apoeb_B1 | AGTATAATAAATGCAGAAAGAAACATAgAAgAgTCTTCCTTTACg |
| Apoeb_B1 | gAggAgggCAgCAAACggAACATTGTCTGTAGAACTTGTTTGGGG |
| Apoeb_B1 | GACGGCAGGAAGGGTTAGAGATCATTAgAAgAgTCTTCCTTTACg |
| Apoeb_B1 | gAggAgggCAgCAAACggAAAGCAGCTTTCCTGGTTTCGCGTTGT |
| Apoeb_B1 | CAGATATTTAAGTTCGCCGTGGCCCTAgAAgAgTCTTCCTTTACg |
| Apoeb_B1 | gAggAgggCAgCAAACggAAGCATGTTACCTGGAGTCAATGAGTC |
| Apoeb_B1 | GGCGAATGGGTCCCCAGACTTCATGTAgAAgAgTCTTCCTTTACg |
| Apoeb_B1 | gAggAgggCAgCAAACggAAACCCCACACAGTACATGAGTAGAGC |
| Apoeb_B1 | TGACGGAATCCAGACAGGAAGCCAATAgAAgAgTCTTCCTTTACg |
| Apoeb_B1 | gAggAgggCAgCAAACggAAGAGGCTCTAACAACATCTTCCATCG |
| Apoeb_B1 | ATGCACTCGAAGTAGAGCAGCGAATTAgAAgAgTCTTCCTTTACg |
| Apoeb_B1 | gAggAgggCAgCAAACggAAAGAGGGTGGAGGCAACATCTTCAGA |
| Apoeb_B1 | CATGCAAAGGTCAAGACCCCAATGCTAgAAgAgTCTTCCTTTACg |
| Apoeb_B1 | gAggAgggCAgCAAACggAAGCCCATGCGGAGTGTCTTATGAAGA |
| Apoeb_B1 | TTTCAAATATTAGGTGATGGCTTGCTAgAAgAgTCTTCCTTTACg |
| Apoeb_B1 | gAggAgggCAgCAAACggAAGAAGCATCAGTCCACGTATATGCAT |

| | |
|---|---|
| Apoeb_B1 | AGAGGTGGAAGGAGGAGCGATAGGCTAgAAgAgTCTTCCTTTACg |
| Apoeb_B1 | gAggAgggCAgCAAACggAAATTCTGTTCACTGATGATATCAGAG |
| Apoeb_B1 | GAAGCACTGAAGTAATCATTGCATGTAgAAgAgTCTTCCTTTACg |
| Apoeb_B1 | gAggAgggCAgCAAACggAATGAAACTAGCACCAGGTCAGCAAAG |
| Apoeb_B1 | GAAAGCAGCATTTTATTTGGTTTCATAgAAgAgTCTTCCTTTACg |
| Apoeb_B1 | gAggAgggCAgCAAACggAAAAACACATACGTGCACGCCTGGGAG |
| Apoeb_B1 | CAGCAAAACATGTACACACATGCATTAgAAgAgTCTTCCTTTACg |
| **Kazald1** | |
| Kazald1_B3 | gTCCCTgCCTCTATATCTTTACTATGTGACTTGTGTTTAACACAG |
| Kazald1_B3 | CTCCAGTCTCACGCACGATCCTTGCTTCCACTCAACTTTAACCCg |
| Kazald1_B3 | gTCCCTgCCTCTATATCTTTAATGTGACTCCTCGACCCCATCCCC |
| Kazald1_B3 | TTGAATGAGGGAGTGATGTCAGAGTTTCCACTCAACTTTAACCCg |
| Kazald1_B3 | gTCCCTgCCTCTATATCTTTCTATTCCGGCAGCACCCCGCCTGCG |
| Kazald1_B3 | CTCGGCTCCGAGAGAGCGGATTCTGTTCCACTCAACTTTAACCCg |
| Kazald1_B3 | gTCCCTgCCTCTATATCTTTTGGTGCATGCGGGAAGCCTTCTCAC |
| Kazald1_B3 | GGAGGTCCTGCGCGGCACCGGCTCGTTCCACTCAACTTTAACCCg |
| Kazald1_B3 | gTCCCTgCCTCTATATCTTTCTGTCACCACGATCCTGAGAGTGCT |
| Kazald1_B3 | CCGCCTTCCCTCTGCAGCCAAAGAGTTCCACTCAACTTTAACCCg |
| Kazald1_B3 | gTCCCTgCCTCTATATCTTTCACCGGGTTCTTGAGCCTGCTCTGC |
| Kazald1_B3 | AGCGGGACACCATTGCTCCTGCCCTTTCCACTCAACTTTAACCCg |
| Kazald1_B3 | gTCCCTgCCTCTATATCTTTTTTGCTCCCAAATGCCATCACTTTT |
| Kazald1_B3 | TTGGCATTTTCAGAAGCTCTCCGGTTTCCACTCAACTTTAACCCg |
| Kazald1_B3 | gTCCCTgCCTCTATATCTTTACACCAGTTCAAATACAGCAGAAAA |
| Kazald1_B3 | TGGCAGCCCTAACCCTGACCATATCTTCCACTCAACTTTAACCCg |
| Kazald1_B3 | gTCCCTgCCTCTATATCTTTGCCACGCTGTAGGTAGTTGGGTGCG |
| Kazald1_B3 | CTCGCCCTCCTCCAGTAGTCTTTGCTTCCACTCAACTTTAACCCg |
| Kazald1_B3 | gTCCCTgCCTCTATATCTTTTTCCTCCGGATCACAGTCTGCACAG |
| Kazald1_B3 | AGCCAGGCAACCCCGGGGCGCCAGGTTCCACTCAACTTTAACCCg |
| Kazald1_B3 | gTCCCTgCCTCTATATCTTTACAGTCACAGGTGTCCTGCACCACC |
| Kazald1_B3 | GCCCTCTAAGTTGGCGCACTCTTGATTCCACTCAACTTTAACCCg |
| Kazald1_B3 | gTCCCTgCCTCTATATCTTTGTTGGTATGGTCCAAGTCACAGATT |
| Kazald1_B3 | GTTATCCCCGCATTTCCCGTAGAAGTTCCACTCAACTTTAACCCg |
| Kazald1_B3 | gTCCCTgCCTCTATATCTTTGGACCTCTCCGTGCCTGAGATCCCC |
| Kazald1_B3 | TGGACAAGCAAACACACTGGGGTTCTTCCACTCAACTTTAACCCg |
| Kazald1_B3 | gTCCCTgCCTCTATATCTTTTCCCATTGGATCCACAGACCGCGGT |
| Kazald1_B3 | GAAATTTGCAGATCTGGGTGTATGTTTCCACTCAACTTTAACCCg |
| Kazald1_B3 | gTCCCTgCCTCTATATCTTTCCTCTGGATGAGCATTTGATGCCTC |
| Kazald1_B3 | GCCCTTCATGGAGCACGGTGAGGTTTTCCACTCAACTTTAACCCg |
| Kazald1_B3 | gTCCCTgCCTCTATATCTTTACAAGATCTGTGGTCCTGATTCACA |

| | |
|---|---|
| Kazald1_B3 | TGACATTCCAGATGTTGTAGGGTGGTTCCACTCAACTTTAACCCg |
| Kazald1_B3 | gTCCCTgCCTCTATATCTTTCGCAGCCAAATATCACATCTTGCCC |
| Kazald1_B3 | TGGATGCCATTGGATAGGCGAAGACTTCCACTCAACTTTAACCCg |
| Kazald1_B3 | gTCCCTgCCTCTATATCTTTTGTCCGTGCCGTCCTTCCTCCACTC |
| Kazald1_B3 | TGTGAGGATCATCCCCAGGTAGTAGTTCCACTCAACTTTAACCCg |
| Kazald1_B3 | gTCCCTgCCTCTATATCTTTGCGGGCCTCCTCTGAACTGAACAGA |
| Kazald1_B3 | GTAACCATCCACTCACTTCATATTTTTCCACTCAACTTTAACCCg |
| Kazald1_B3 | gTCCCTgCCTCTATATCTTTCATCAGTGATCCGGACGGCTTGAAT |
| Kazald1_B3 | TCCTAGCAAAGCAATGGTAGGTTCCTTCCACTCAACTTTAACCCg |
| Kazald1_B3 | gTCCCTgCCTCTATATCTTTCAGTAGCTGTGACCTCTCCCACCTT |
| Kazald1_B3 | GATCTGGAGTGAGCACTGTAAGGCTTTCCACTCAACTTTAACCCg |
| Kazald1_B3 | gTCCCTgCCTCTATATCTTTGTAAAGATAGTGCTGTCGTGTTGAG |
| Kazald1_B3 | CATCGTATATTTCATGCCGAGGCTTTTCCACTCAACTTTAACCCg |
| Kazald1_B3 | gTCCCTgCCTCTATATCTTTAGTCATCTGAGTCCTCGCCTTCTTC |
| Kazald1_B3 | TGGGCCAGTGTTATACGTGTTAGTATTCCACTCAACTTTAACCCg |
| Kazald1_B3 | gTCCCTgCCTCTATATCTTTCTGAAATCTGGATAAAGTGTGTGGT |
| Kazald1_B3 | GCTGATGCTGATTGGATCACGTTGCTTCCACTCAACTTTAACCCg |
| Kazald1_B3 | gTCCCTgCCTCTATATCTTTAACATGTCTGTCCATTTTCTGAGTG |
| Kazald1_B3 | ATCTCCATGAAGACTTTATGAGATGTTCCACTCAACTTTAACCCg |
| Kazald1_B3 | gTCCCTgCCTCTATATCTTTATGTTGGGAATGTGGCAGTAGTTTT |
| Kazald1_B3 | CACTGAATTAACCACAGTTGTAGGATTCCACTCAACTTTAACCCg |
| Kazald1_B3 | gTCCCTgCCTCTATATCTTTAATAATGTGAGGGCTAAACAAATGA |
| Kazald1_B3 | CCTTGTGTCCTCTTTGTCTTCTCTCTTCCACTCAACTTTAACCCg |
| Kazald1_B3 | gTCCCTgCCTCTATATCTTTTTCCCTTTGGCTTTCAAACCTAAAA |
| Kazald1_B3 | TTAGAGAGTAGCATTTGGAATCTGATTCCACTCAACTTTAACCCg |
| Kazald1_B3 | gTCCCTgCCTCTATATCTTTGTCCCCTTGACAACAAGTACCAGGT |
| Kazald1_B3 | ATGAAGTCAATAAATTACTTGAGAATTCCACTCAACTTTAACCCg |
| Kazald1_B3 | gTCCCTgCCTCTATATCTTTAATGTATGTCTATAAACAAAGGCGT |
| Kazald1_B3 | AAAGCATTTACACAAGCATTGCAAGTTCCACTCAACTTTAACCCg |
| Kazald1_B3 | gTCCCTgCCTCTATATCTTTTAAAGGGTATTTAGAAAGAAACAAC |
| Kazald1_B3 | TACCAAAGAATGTTTACACATTATTTTCCACTCAACTTTAACCCg |
| Kazald1_B3 | gTCCCTgCCTCTATATCTTTCATGTGGATAATTTGATATCATTTT |
| Kazald1_B3 | AACATGGGTTTATTTTACAAAACATTTCCACTCAACTTTAACCCg |
| **Raldh2** | |
| Raldh2_B2 | CCTCgTAAATCCTCATCAAAGGGTCCGGCTTCACCTCCCCGGGCA |
| Raldh2_B2 | AGGTGCAGGGAGGCCATGAGAGCCGAAATCATCCAgTAAACCgCC |
| Raldh2_B2 | CCTCgTAAATCCTCATCAAAGTTCTGCCACTCGTTGTTGATAAAA |
| Raldh2_B2 | CACTGGGAATACTTTGCCGCTCACGAAATCATCCAgTAAACCgCC |
| Raldh2_B2 | CCTCgTAAATCCTCATCAAAAATCATCTCTGACGTGGCTGGATTG |

| | |
|---|---|
| Raldh2_B2 | AGCCTTGTCAGCTTCCTGGACTTCAAAATCATCCAgTAAACCgCC |
| Raldh2_B2 | CCTCgTAAATCCTCATCAAACGCAGCTCGCACTGCTTTGTCTATA |
| Raldh2_B2 | GACGGAGCCCATGGAGAAGGCAAGTAAATCATCCAgTAAACCgCC |
| Raldh2_B2 | CCTCgTAAATCCTCATCAAACCTCTCCGATGCGTCCATTCTCCGC |
| Raldh2_B2 | GTCAGCCAGTTTGCTCAAGAGTCGGAAATCATCCAgTAAACCgCC |
| Raldh2_B2 | CCTCgTAAATCCTCATCAAAAAGAAGGGCCAAGTCCCTTTCTACG |
| Raldh2_B2 | GCCGCTGTCCAGAGATTCAATGGTAAAATCATCCAgTAAACCgCC |
| Raldh2_B2 | CCTCgTAAATCCTCATCAAAAATATCTCAGTGTCTTTATCGCCCC |
| Raldh2_B2 | CATGAATCTTGTCGGCCCAGCCTGCAAATCATCCAgTAAACCgCC |
| Raldh2_B2 | CCTCgTAAATCCTCATCAAAAATCGCCATCTGCTGGGATGGTTAT |
| Raldh2_B2 | TGGGCTCGTGCCTCGTGAACGTGAAAAATCATCCAgTAAACCgCC |
| Raldh2_B2 | CCTCgTAAATCCTCATCAAATCTTCCAGGCAAACATCAGCAGGGG |
| Raldh2_B2 | TGTTACCACAGCACAGAGCTGGAGCAAATCATCCAgTAAACCgCC |
| Raldh2_B2 | CCTCgTAAATCCTCATCAAATTTGTTCGGCCGGTTTAATAACCAC |
| Raldh2_B2 | ATCCCATGTGGAGCGCGCTGAGCGGAAATCATCCAgTAAACCgCC |
| Raldh2_B2 | CCTCgTAAATCCTCATCAAAGCGGAAACCCAGCCTCTTTGATGAG |
| Raldh2_B2 | ATCCCGGCAAAATATTGACGACTCCAAATCATCCAgTAAACCgCC |
| Raldh2_B2 | CCTCgTAAATCCTCATCAAACCATTGCTGCGCCTGCCGTCGGACC |
| Raldh2_B2 | CTACTTTATCGATGCCGGAGTGAGAAAATCATCCAgTAAACCgCC |
| Raldh2_B2 | CCTCgTAAATCCTCATCAAATCCCAACTTCAGTGGATCCAGTAAA |
| Raldh2_B2 | TCCTTCCAGCTGCCTCCTGGATCAGAAATCATCCAgTAAACCgCC |
| Raldh2_B2 | CCTCgTAAATCCTCATCAAAGCTCCAGGGTAACTCTTTTCAAATT |
| Raldh2_B2 | AAATAATGTTGGGGCTCTTCCCACCAAATCATCCAgTAAACCgCC |
| Raldh2_B2 | CCTCgTAAATCCTCATCAAACTGCATAGTCCAAATCGGCATCTGC |
| Raldh2_B2 | AGAAAACGCCTTGGTGGGCTTGCTCAAATCATCCAgTAAACCgCC |
| Raldh2_B2 | CCTCgTAAATCCTCATCAAACTGCAGTGCAACACTGCCCTTGATT |
| Raldh2_B2 | TGGTCTCCTCAACAAATGTCCGGGAAAATCATCCAgTAAACCgCC |
| Raldh2_B2 | CCTCgTAAATCCTCATCAAACACTCCGCCGGACAAACTCATCGTA |
| Raldh2_B2 | CTACTATACGCCTCTTGGCACGCTCAAATCATCCAgTAAACCgCC |
| Raldh2_B2 | CCTCgTAAATCCTCATCAAAGTTCAGTGGAAGGATCAAAGGGACT |
| Raldh2_B2 | ACTGTTTCATATCCGTCTGTGGACCAAATCATCCAgTAAACCgCC |
| Raldh2_B2 | CCTCgTAAATCCTCATCAAATTTGGATGAGATCCAGAATTTTGTT |
| Raldh2_B2 | CTAGTTTAGCGCCTTCCATGACACCAAATCATCCAgTAAACCgCC |
| Raldh2_B2 | CCTCgTAAATCCTCATCAAACCCTTTCTTCCTAAGCCTTTGCCCC |
| Raldh2_B2 | GAAAACACAGTTGGTTCCACGAATAAAATCATCCAgTAAACCgCC |
| Raldh2_B2 | CCTCgTAAATCCTCATCAAAGAGTATTTGTTGGACGGGTCCAAAA |
| Raldh2_B2 | GATCACTTCTTCCATGGTTTTAAACAAATCATCCAgTAAACCgCC |
| Raldh2_B2 | CCTCgTAAATCCTCATCAAATCCAAACTCCGAGCTGTTCGCTCTC |
| Raldh2_B2 | GTCATTTGTGAAAACGGCTGCTACCAAATCATCCAgTAAACCgCC |

| | |
|---|---|
| Raldh2_B2 | CCTCgTAAATCCTCATCAAACGAGGAAACCGTCAGGGCTTTGCTG |
| Raldh2_B2 | AATCCAGACTGTTCCAGCTTGCATTAAATCATCCAgTAAACCgCC |
| Raldh2_B2 | CCTCgTAAATCCTCATCAAACTGGGCATTCAAGGCATTGTAGCAA |
| Raldh2_B2 | GGACATCTTATATCCTCCGAAAGGAAAATCATCCAgTAAACCgCC |
| Raldh2_B2 | CCTCgTAAATCCTCATCAAAGTACTCGCCCATTTCCCTCCCATTA |
| Raldh2_B2 | CTTGGCTTCTGTGTACTCCCGCAATAAATCATCCAgTAAACCgCC |
| Raldh2_B2 | CCTCgTAAATCCTCATCAAAAGCGCTCTTAGGAGTTCTTTTGGGG |
| Raldh2_B2 | GCGTGCTCTTCGCCCCACGTGGTCTAAATCATCCAgTAAACCgCC |
| Raldh2_B2 | CCTCgTAAATCCTCATCAAAGTTGCTGTAACGAATACTGATGGGG |
| Raldh2_B2 | CTGTATAACACTGCATTCCTGAGTGAAATCATCCAgTAAACCgCC |
| Raldh2_B2 | CCTCgTAAATCCTCATCAAATTTGCCAACTAGGATCTGAAATATA |
| Raldh2_B2 | ATTTACGGCAAGGCATCTGGTTCACAAATCATCCAgTAAACCgCC |
| Raldh2_B2 | CCTCgTAAATCCTCATCAAAGGACACCATGCGGAAACCCTTGATA |
| Raldh2_B2 | CGTACATTGTGTAACGGTGGGGTGGAAATCATCCAgTAAACCgCC |
| Raldh2_B2 | CCTCgTAAATCCTCATCAAAGTAAGACTTAGCATGTTATTTTGTG |
| Raldh2_B2 | TGTTGAACGTTCATTAAATGGTTTAAAATCATCCAgTAAACCgCC |
| Raldh2_B2 | CCTCgTAAATCCTCATCAAAGCACTGTATATGTTTAATAGCTTGA |
| Raldh2_B2 | TCTAGCATCCTTCAACTTACTGCACAAATCATCCAgTAAACCgCC |
| Raldh2_B2 | CCTCgTAAATCCTCATCAAATGGGAAACAGACCCCAACCTGCAGT |
| Raldh2_B2 | CAAAGTATACAGGGCTGTTCATCCAAAATCATCCAgTAAACCgCC |
| Raldh2_B2 | CCTCgTAAATCCTCATCAAATAGCGTGTACACGCACTCTTGTTTT |
| Raldh2_B2 | AGGGGCGTCGAAACACTGATGATATAAATCATCCAgTAAACCgCC |
| Raldh2_B2 | CCTCgTAAATCCTCATCAAAACTCTCAGTTTCCTGAAGAAGCATT |
| Raldh2_B2 | GTCAGAGCAGAGAAGCAATTTAATTAAATCATCCAgTAAACCgCC |
| Raldh2_B2 | CCTCgTAAATCCTCATCAAAAAGCACAACAGTCTGGATTGGTTGA |
| Raldh2_B2 | CATAAATATCCTAGGTTATGGTGACAAATCATCCAgTAAACCgCC |
| Raldh2_B2 | CCTCgTAAATCCTCATCAAATAAAGACAAAGTGGGTTTATATTTT |
| Raldh2_B2 | TAGACACCAAAGTTTATCATACAGAAAATCATCCAgTAAACCgCC |
| Raldh2_B2 | CCTCgTAAATCCTCATCAAACATGTTGACATGACAGTGTTATCTA |
| Raldh2_B2 | GATCAGCATCTCCTGCATGATGTGGAAATCATCCAgTAAACCgCC |
| **Bmp2** | |
| Bmp2_B3 | gTCCCTgCCTCTATATCTTTCAGGCTAGTGCTCGCCCGCCTCCCC |
| Bmp2_B3 | CCTCTTAAAGGGGCCGCGGCCGCTCTTCCACTCAACTTTAACCCg |
| Bmp2_B3 | gTCCCTgCCTCTATATCTTTCGAGTCCCGGCTCTCTTCCTGCCCC |
| Bmp2_B3 | TCCAATCAAGAGTCTGCGTAGCTCGTTCCACTCAACTTTAACCCg |
| Bmp2_B3 | gTCCCTgCCTCTATATCTTTTTGGGACGAGCCGAAGGCTGCTTTT |
| Bmp2_B3 | GTCTCCGCTCCCTCTCTGTATCATGTTCCACTCAACTTTAACCCg |
| Bmp2_B3 | gTCCCTgCCTCTATATCTTTTGCTGCTGCTGCGGGCGTGCGGGCG |
| Bmp2_B3 | CCCTGCGGCGTTCCTCGAGGCGGCTTTCCACTCAACTTTAACCCg |

| | |
|---|---|
| Bmp2_B3 | gTCCCTgCCTCTATATCTTTGGGCAGCCCAGTGTGTTGTGCGCTG |
| Bmp2_B3 | GTCCTCTGCCCTGTCACAGTGCAGGTTCCACTCAACTTTAACCCg |
| Bmp2_B3 | gTCCCTgCCTCTATATCTTTCCTCGGAATACTCCGGGCAGTGGGG |
| Bmp2_B3 | GTGTGCACAAGCGGCGCCCTGCCCGTTCCACTCAACTTTAACCCg |
| Bmp2_B3 | gTCCCTgCCTCTATATCTTTCTCATGGTGCCCAGCGGGTCACCCC |
| Bmp2_B3 | GTCCCGGCCACAGTCCTCCGCCTCCTTCCACTCAACTTTAACCCg |
| Bmp2_B3 | gTCCCTgCCTCTATATCTTTGCAGCCGTAAAGTCGAGCCTGGAAA |
| Bmp2_B3 | CGGGGTCACTCCAAACGTGTGCCGTTTCCACTCAACTTTAACCCg |
| Bmp2_B3 | gTCCCTgCCTCTATATCTTTCTCTCGGGCTGTCCTGTCCACATCC |
| Bmp2_B3 | ACCTGGGGTCCGGAGGAGGGCTGTCTTCCACTCAACTTTAACCCg |
| Bmp2_B3 | gTCCCTgCCTCTATATCTTTACGAGCGCATCCCGGCCACCATCGT |
| Bmp2_B3 | GCACTTGGTACAGGAGCAGCACCAGTTCCACTCAACTTTAACCCg |
| Bmp2_B3 | gTCCCTgCCTCTATATCTTTAGTGCTGCTGCTGCTGCTGCTGGGG |
| Bmp2_B3 | GCTCGAACTCGCGCAGCACGTCCTCTTCCACTCAACTTTAACCCg |
| Bmp2_B3 | gTCCCTgCCTCTATATCTTTGTCGCGCATGTAGGGCGGGATGGGG |
| Bmp2_B3 | GAGGTCCAGGCCCTGGCCCTGGTACTTCCACTCAACTTTAACCCg |
| Bmp2_B3 | gTCCCTgCCTCTATATCTTTTGGCGCGGCTGGCGGAGCGCTCCTG |
| Bmp2_B3 | CTTCATGGTGGAAGCTGCGGACGGTTTCCACTCAACTTTAACCCg |
| Bmp2_B3 | gTCCCTgCCTCTATATCTTTTTGCTTCAGGCAGGCCTTCCAAAAC |
| Bmp2_B3 | AGAAGAAACGCTGCAGCAGTTTTCCTTCCACTCAACTTTAACCCg |
| Bmp2_B3 | gTCCCTgCCTCTATATCTTTCCTCGTTAGGGATAGAAGTCAAGTT |
| Bmp2_B3 | TCCGGAGTTCAGCCGAGGTGATAAATTCCACTCAACTTTAACCCg |
| Bmp2_B3 | gTCCCTgCCTCTATATCTTTAGGTCTCCGTGACCAGTTCTCGAAA |
| Bmp2_B3 | TGCGGTGCGTGCTGCTGTTCCCGCCTTCCACTCAACTTTAACCCg |
| Bmp2_B3 | gTCCCTgCCTCTATATCTTTGGGGCTTGAGGATTTCATAAATATT |
| Bmp2_B3 | TGACCAGCTCCTGGGGCCCCGCGGCTTCCACTCAACTTTAACCCg |
| Bmp2_B3 | gTCCCTgCCTCTATATCTTTGCACCAGCCGGGTGTCCAGCAGCCG |
| Bmp2_B3 | AGCTCTCCCAGCTGCTGGCGTTGGGTTCCACTCAACTTTAACCCg |
| Bmp2_B3 | gTCCCTgCCTCTATATCTTTACCGCAGCACGGCCGGGGTCACGTC |
| Bmp2_B3 | CGTGGTTGGGCTGCCCGTGCACCTGTTCCACTCAACTTTAACCCg |
| Bmp2_B3 | gTCCCTgCCTCTATATCTTTCTAGGTGGGCCACCTCCACCACGAA |
| Bmp2_B3 | GCCGCTGGGCCAGGCCGCGCTCACTTTCCACTCAACTTTAACCCg |
| Bmp2_B3 | gTCCCTgCCTCTATATCTTTTGCTGCGGGGAGCGGCGGAGGCGGA |
| Bmp2_B3 | CGGAGCTGGGCCCAGCTGTGCGCGTTTCCACTCAACTTTAACCCg |
| Bmp2_B3 | gTCCCTgCCTCTATATCTTTTCGTGGCCAAACGTTACTAAGAGGG |
| Bmp2_B3 | CGCCGGTGCAGCGGGTGGCCCTTGCTTCCACTCAACTTTAACCCg |
| Bmp2_B3 | gTCCCTgCCTCTATATCTTTTGCCGCTGGCGCGCCTGGCGCTTTT |
| Bmp2_B3 | CGGCAGTTGGACTTGAGGCGCTTGCTTCCACTCAACTTTAACCCg |
| Bmp2_B3 | gTCCCTgCCTCTATATCTTTCGTCGCTGAAGTCCACGTAGAGGGG |

| | |
|---|---|
| Bmp2_B3 | GCGCCACGATCCAGTCATTCCAGCCTTCCACTCAACTTTAACCCg |
| Bmp2_B3 | gTCCCTgCCTCTATATCTTTGGCAATAGAAGGCGTGGTATCCCGG |
| Bmp2_B3 | CCGCCAGAGGGAAGGGGCAGTCCCCTTCCACTCAACTTTAACCCg |
| Bmp2_B3 | gTCCCTgCCTCTATATCTTTTGGCATGGTTCGTGGAGTTCATGTG |
| Bmp2_B3 | TCACAGAGTTGACCAAAGTCTGCACTTCCACTCAACTTTAACCCg |
| Bmp2_B3 | gTCCCTgCCTCTATATCTTTCGCAGCAAGCCTTGGGGATGTTGGC |
| Bmp2_B3 | TGGAGATGGCGCTCAGCTCGGTGGGTTCCACTCAACTTTAACCCg |
| Bmp2_B3 | gTCCCTgCCTCTATATCTTTCCTTTTCGTTCTCGTCCAGGTAGAG |
| Bmp2_B3 | CCATGTCCTGGTAATTCTTGAGCACTTCCACTCAACTTTAACCCg |
| Bmp2_B3 | gTCCCTgCCTCTATATCTTTCCTAATGACACCCACAACCCTCCAC |
| Bmp2_B3 | TTTGTTGGTCCATTGGGTGGTTGGCTTCCACTCAACTTTAACCCg |
| Bmp2_B3 | gTCCCTgCCTCTATATCTTTATTAAAGTGTCAGCTTTTGTGTTTT |
| Bmp2_B3 | CATAAATAAAAGTCTTCATCGGGAATTCCACTCAACTTTAACCCg |
| Bmp2_B3 | gTCCCTgCCTCTATATCTTTCCAACTTTGTAGACATAAATATTTT |
| Bmp2_B3 | TAATTCTCTGGATAAAATACATTTTTTCCACTCAACTTTAACCCg |
| Bmp2_B3 | gTCCCTgCCTCTATATCTTTGACCATTATACTTCATGTGCTGGGG |
| Bmp2_B3 | TTGTAAATAAATACAAAATAGGCATTTCCACTCAACTTTAACCCg |
| Bmp2_B3 | gTCCCTgCCTCTATATCTTTCAAAATTCAAACGGCTGCAGATTTT |
| Bmp2_B3 | AAACACATCCCTCACAACTTGCAAATTCCACTCAACTTTAACCCg |
| Bmp2_B3 | gTCCCTgCCTCTATATCTTTATGTCCCTGCGAACACATCCACTGT |
| Bmp2_B3 | AAAATAGGACCGGATCATCACCAGGTTCCACTCAACTTTAACCCg |
| Bmp2_B3 | gTCCCTgCCTCTATATCTTTGAGTAAAACAGAAAATCATCATTTT |
| Bmp2_B3 | ATTCCAAAAGGTACAACCCAGGCCATTCCACTCAACTTTAACCCg |
| Bmp2_B3 | gTCCCTgCCTCTATATCTTTTGGGATGGACATGGTACGTACGGGG |
| Bmp2_B3 | ACGTGGCCCACATATGCCTCACTGATTCCACTCAACTTTAACCCg |
| Bmp2_B3 | gTCCCTgCCTCTATATCTTTCATAGTAATAGAACCCAGCTTCATA |
| Bmp2_B3 | ACTTGTGGAGCGGTGGGTGGCAGGGTTCCACTCAACTTTAACCCg |
| **Bmp7** | |
| Bmp7_B1 | gAggAgggCAgCAAACggAAGGTCAGCAGGGCAGCCAGGTTCCCC |
| Bmp7_B1 | CGAGCCGACCAAGAGTGGCGCCCACTAgAAgAgTCTTCCTTTACg |
| Bmp7_B1 | gAggAgggCAgCAAACggAAGTTGTCCAAGGAGAAGTCCGAAAAG |
| Bmp7_B1 | CCGGTGGATGAAGCTCGAGTGCACCTAgAAgAgTCTTCCTTTACg |
| Bmp7_B1 | gAggAgggCAgCAAACggAACTCCCGCCGCTCCTGGCTGCGCAAC |
| Bmp7_B1 | CAAGATGGACAGGATTTCTCGCTGCTAgAAgAgTCTTCCTTTACg |
| Bmp7_B1 | gAggAgggCAgCAAACggAAAGGTGAGGCCGGGGCCGGTGCGGCA |
| Bmp7_B1 | ATGGGCGCCGAGTTCTGCTTGCCGTTAgAAgAgTCTTCCTTTACg |
| Bmp7_B1 | gAggAgggCAgCAAACggAAGGCTTGTAGGGGTAGGAGAAGCCCC |
| Bmp7_B1 | GCGGGCGGGCCCTGGGTGGTGAAGATAgAAgAgTCTTCCTTTACg |
| Bmp7_B1 | gAggAgggCAgCAAACggAAAGGAAGTTGTTGTCCTGCAGGCTGG |

| | |
|---|---|
| Bmp7_B1 | AAGCTCATGACCATGTCGGCGTCGTTAgAAgAgTCTTCCTTTACg |
| Bmp7_B1 | gAggAgggCAgCAAACggAATCCTTGTCATGCTCGACCAAGTTGA |
| Bmp7_B1 | CGGTGATGGCGCCTCTGGTGGAAGATAgAAgAgTCTTCCTTTACg |
| Bmp7_B1 | gAggAgggCAgCAAACggAAATTTTCGCGAGATCAAACCGAAACT |
| Bmp7_B1 | GCGGCAGTGACGGCTTCGCCCTCTGTAgAAgAgTCTTCCTTTACg |
| Bmp7_B1 | gAggAgggCAgCAAACggAAATGTAATCCTTATAGATCCGAAACT |
| Bmp7_B1 | AATGTCTCATTATCAAAGCGTTCCCTAgAAgAgTCTTCCTTTACg |
| Bmp7_B1 | gAggAgggCAgCAAACggAATGCAGTACCTGGTAGATGCTGATCT |
| Bmp7_B1 | AAGTCTGAATCCCTCCCTGGGTGTTTAgAAgAgTCTTCCTTTACg |
| Bmp7_B1 | gAggAgggCAgCAAACggAACAGATAACGCGGCTGTCTAATGGGA |
| Bmp7_B1 | AAGACGAGCCATCCCTCTTCGGCAGTAgAAgAgTCTTCCTTTACg |
| Bmp7_B1 | gAggAgggCAgCAAACggAACAGTGGTTACTGGTTACTGTGATGT |
| Bmp7_B1 | CCGAGATTGTACCGTGGATTCATCATAgAAgAgTCTTCCTTTACg |
| Bmp7_B1 | gAggAgggCAgCAAACggAATCAACGCTCTCGACAGAGAGCTGTA |
| Bmp7_B1 | GCCATCTTGGGATTGATGCTTTCTCTAgAAgAgTCTTCCTTTACg |
| Bmp7_B1 | gAggAgggCAgCAAACggAATGCGGCCCATGTCTTCCAATCAAAC |
| Bmp7_B1 | AAGGCCACCATGAAGGGCTGCTTATTAgAAgAgTCTTCCTTTACg |
| Bmp7_B1 | gAggAgggCAgCAAACggAACGATTTTGGTTTCTTTGTTTGCCCC |
| Bmp7_B1 | GCTTCCTGGTTCTTTGGTGCCTTGGTAgAAgAgTCTTCCTTTACg |
| Bmp7_B1 | gAggAgggCAgCAAACggAATTCTCTGCGACATTCGAAACTCGCA |
| Bmp7_B1 | CATGCTTGCCTCTGGTCGCTGCTACTAgAAgAgTCTTCCTTTACg |
| Bmp7_B1 | gAggAgggCAgCAAACggAAAAACTGACATACAGCTCATGTTTCT |
| Bmp7_B1 | ATCCAGTCCTGCCAGCCAAGGTCCCTAgAAgAgTCTTCCTTTACg |
| Bmp7_B1 | gAggAgggCAgCAAACggAATAGGCCGCATAACCTTCTGGGGCAA |
| Bmp7_B1 | GGGAAAGAACATTCTCCTTCGCAGTTAgAAgAgTCTTCCTTTACg |
| Bmp7_B1 | gAggAgggCAgCAAACggAATTTGTGGCATTCATGTAGGAATTCA |
| Bmp7_B1 | TGAACCAGCGTCTGAACAATGGCGTTAgAAgAgTCTTCCTTTACg |
| Bmp7_B1 | gAggAgggCAgCAAACggAACGAGATGGGGTTGAGTTGTGTGGGG |
| Bmp7_B1 | GTTGGAGCTGTCGTCAAAGTAGAGATAgAAgAgTCTTCCTTTACg |
| Bmp7_B1 | gAggAgggCAgCAAACggAACATATTCCTGTATTTCTTCAGGATG |
| Bmp7_B1 | CTAGTGGCAGCCGCATGCCCGCACCTAgAAgAgTCTTCCTTTACg |
| Bmp7_B1 | gAggAgggCAgCAAACggAAGACTTTGCGACCACGCCTCCGGCCT |
| Bmp7_B1 | TAAAAGCAGACTCCCGAGGGACAGCTAgAAgAgTCTTCCTTTACg |
| Bmp7_B1 | gAggAgggCAgCAAACggAAACATAAAACCTTTTGCGAATCCATA |
| Bmp7_B1 | TTTGTATTCATAAACCTGATGCAGGTAgAAgAgTCTTCCTTTACg |
| Bmp7_B1 | gAggAgggCAgCAAACggAAAAAGAGAAAACATACTGCCAGAAAA |
| Bmp7_B1 | TATGTTGGCGCAGGCAGCGACACTGTAgAAgAgTCTTCCTTTACg |
| Bmp7_B1 | gAggAgggCAgCAAACggAAAAATCACCCGGGAGATTCAGTCCTT |
| Bmp7_B1 | ATTCATAACATTCTGACACGGGTGGTAgAAgAgTCTTCCTTTACg |

| | |
|---|---|
| Bmp7_B1 | gAggAgggCAgCAAACggAAACGTTTTGTCACGTTCAGGCGGTTT |
| Bmp7_B1 | ACAAGAAAGCAAAGGGCTGTTTAAATAgAAgAgTCTTCCTTTACg |
| Bmp7_B1 | gAggAgggCAgCAAACggAAATGCTCCCACTTCGGAAGCAGCGGT |
| Bmp7_B1 | TAAGTCAATATTCAATTAGCCGACCTAgAAgAgTCTTCCTTTACg |
| Bmp7_B1 | gAggAgggCAgCAAACggAAACCATCGTGACTGCCAAAGTCAACA |
| Bmp7_B1 | AACATCCAAGTAATTAAACCCACATTAgAAgAgTCTTCCTTTACg |
| Bmp7_B1 | gAggAgggCAgCAAACggAAGTTCCCATCGCTCGCTTTTGCACGT |
| Bmp7_B1 | AGAAATAGCAAAACGTTAAGAACTCTAgAAgAgTCTTCCTTTACg |
| Bmp7_B1 | gAggAgggCAgCAAACggAAAATATTCAGAAGTAACTTATTTTGT |
| Bmp7_B1 | GAGCCTTCAAACTGTACAAACCTGTTAgAAgAgTCTTCCTTTACg |
| Bmp7_B1 | gAggAgggCAgCAAACggAACTTTGGACAATGGCATTTGTGTTTT |
| Bmp7_B1 | ATGGCTGACAGACACAAGCACACATTAgAAgAgTCTTCCTTTACg |
| Bmp7_B1 | gAggAgggCAgCAAACggAAGGGACCTAGTTAAAGATTTAAAACA |
| Bmp7_B1 | CCCAATGAGAAGTGCAAGACTGCTGTAgAAgAgTCTTCCTTTACg |
| Bmp7_B1 | gAggAgggCAgCAAACggAAGTTGCTCCCAAGCCCAGGTCAAGGA |
| Bmp7_B1 | TGAGCGGGTTACCACCACACATGTGTAgAAgAgTCTTCCTTTACg |
| Bmp7_B1 | gAggAgggCAgCAAACggAAGGGCTGAAGTTGCCACCACAATGTT |
| Bmp7_B1 | CCTCTGGGGAGCTCAGCCTAGTACATAgAAgAgTCTTCCTTTACg |
| Bmp7_B1 | gAggAgggCAgCAAACggAATCTCTGGTGTGCCATGGGCAAAATG |
| Bmp7_B1 | CTTTATAGCAGTGCTTCTGGCATTTTAgAAgAgTCTTCCTTTACg |
| Bmp7_B1 | gAggAgggCAgCAAACggAAAGGGGTAGGGTTAGCCCCTCAGATA |
| Bmp7_B1 | CACAGCATTCTAGCAAAGCAAACACTAgAAgAgTCTTCCTTTACg |
| Bmp7_B1 | gAggAgggCAgCAAACggAAATTCCCGCCGTCAGTCAAACAGAAT |
| Bmp7_B1 | AGAATTGCTGGCCACCTAAAATGAGTAgAAgAgTCTTCCTTTACg |
| **Prrx1** | |
| Prrx1_B2 | CCTCGTAAATCCTCATCAAACTTCTTGGCCTGCAGGGTGTCCTGC |
| Prrx1_B2 | GTCCAGCAGGTGACTGACCGAGAAGAAATCATCCAGTAAACCGCC |
| Prrx1_B2 | CCTCGTAAATCCTCATCAAATCCTGTTTCGCCGCTGCTTTCTTTT |
| Prrx1_B2 | CCTGGAGTTGACTGCTGTTGAAGGTAAATCATCCAGTAAACCGCC |
| Prrx1_B2 | CCTCGTAAATCCTCATCAAAGTCTGGGTAATGCGTTCTCTCAAAA |
| Prrx1_B2 | TCGTGCAAGATCCTCTCTTACAAAGAAATCATCCAGTAAACCGCC |
| Prrx1_B2 | CCTCGTAAATCCTCATCAAAAACTCTAGCCTCTGTGAGATTAACT |
| Prrx1_B2 | TGCTCTTCGGTTCTGGAACCACACCAAATCATCCAGTAAACCGCC |
| Prrx1_B2 | CCTCGTAAATCCTCATCAAATAGGATTTCAGCAGAGAGGCGTTTT |
| Prrx1_B2 | TGTTCCACAGCTGTCATGTCCCCAGAAATCATCCAGTAAACCGCC |
| Prrx1_B2 | CCTCGTAAATCCTCATCAAAATGGCACTGTATGGAGAGGCCGTGC |
| Prrx1_B2 | TTGGCACAAGGAGGAGAATAAGTAGAAATCATCCAGTAAACCGCC |
| Prrx1_B2 | CCTCGTAAATCCTCATCAAAATGTTCATACCCTGTTCGGGGCCAG |
| Prrx1_B2 | AGTCGCAGGTTTGCAATGCTGTTGGAAATCATCCAGTAAACCGCC |

| | |
|---|---|
| Prrx1_B2 | CCTCGTAAATCCTCATCAAACTTTGTAAACTATATTCTTTTGCCT |
| Prrx1_B2 | TCTTAGTTGACCGTTGGCACTTGGTAAATCATCCAGTAAACCGCC |
| Prrx1_B2 | CCTCGTAAATCCTCATCAAAAATGCAGGAAGTGCACAAGTGTTTT |
| Prrx1_B2 | CCTTAGTCCACTTTCTGCTATTTTCAAATCATCCAGTAAACCGCC |
| Prrx1_B2 | CCTCGTAAATCCTCATCAAAATTCACCCTGCGTTTCTCCATTTTG |
| Prrx1_B2 | GAAGATGAACTGTGTTTCCGTGATTAAATCATCCAGTAAACCGCC |
| Prrx1_B2 | CCTCGTAAATCCTCATCAAAGGGGAAGGTGGTGGCATTGTCCCAG |
| Prrx1_B2 | ATCTAGCAAAGCATGAAGAGAACAAAAATCATCCAGTAAACCGCC |
| Prrx1_B2 | CCTCGTAAATCCTCATCAAAAACCAAAAGAATTTCTTGTGATTTT |
| Prrx1_B2 | CTGTCTAGGCACTTGGTGTTGGTAAAAATCATCCAGTAAACCGCC |
| Prrx1_B2 | CCTCGTAAATCCTCATCAAAGCGAAAGACGCTGGAAAGCGGAATT |
| Prrx1_B2 | AGCCCTTGACATGTATTGCTTTTGCAAATCATCCAGTAAACCGCC |
| Prrx1_B2 | CCTCGTAAATCCTCATCAAATATGATACCTGACAACATTTTAAAA |
| Prrx1_B2 | CACACATACAATCACAACATGTGCTAAATCATCCAGTAAACCGCC |
| Prrx1_B2 | CCTCGTAAATCCTCATCAAAATATCTTTGGCAAGGAAGCGCCGCA |
| Prrx1_B2 | TGCTTGTCTGTAGTTAGAGAAAAGTAAATCATCCAGTAAACCGCC |
| Prrx1_B2 | CCTCGTAAATCCTCATCAAACCATTACTCTTTATTGGCATAGGTG |
| Prrx1_B2 | TGGTCTCATAGTCCGCTGCTGCGAGAAATCATCCAGTAAACCGCC |
| Prrx1_B2 | CCTCGTAAATCCTCATCAAATGCAACTTGAAGATACACTTCGAAT |
| Prrx1_B2 | TTGAACCGAAGATCACACCAGCCTCAAATCATCCAGTAAACCGCC |
| Prrx1_B2 | CCTCGTAAATCCTCATCAAAAGTTCCCTTACTGCAGACCAGGGCA |
| Prrx1_B2 | AGGGAATTGCCCTTATTTTGCACATAAATCATCCAGTAAACCGCC |
| Prrx1_B2 | CCTCGTAAATCCTCATCAAAATAAACCTTGTGCATCTGTCATTCT |
| Prrx1_B2 | TTAAACAATTATGAATCACAACTATAAATCATCCAGTAAACCGCC |
| Prrx1_B2 | CCTCGTAAATCCTCATCAAAAAAAGGCAGGGCAATGAGGGACCGG |
| Prrx1_B2 | TTTCTGTATCCCACGCAAACCAACAAAATCATCCAGTAAACCGCC |
| Prrx1_B2 | CCTCGTAAATCCTCATCAAAATAGAAGAAAGCAATTTTCGGATAG |
| Prrx1_B2 | GTGTCTGTAAGCTCTAGCTTTACATAAATCATCCAGTAAACCGCC |
| Prrx1_B2 | CCTCGTAAATCCTCATCAAATTCACCGGATGCTTGCCTCACTTTG |
| Prrx1_B2 | TGTCTTATTGTCTCTATGATCCTTCAAATCATCCAGTAAACCGCC |
| Prrx1_B2 | CCTCGTAAATCCTCATCAAAAACAGCAGCTCCTGTACTGCCAAAA |
| Prrx1_B2 | ACCTCTGAGTTTTAGCACACTTTCAAAATCATCCAGTAAACCGCC |
| Prrx1_B2 | CCTCGTAAATCCTCATCAAAGCGTAGAGATGCTCTGTTTGGATAT |
| Prrx1_B2 | CCTGGTGAGTGTAGTTCATTAATATAAATCATCCAGTAAACCGCC |
| Prrx1_B2 | CCTCGTAAATCCTCATCAAAGACTGTAATCGCAGCCTTGCTACCT |
| Prrx1_B2 | TGACATGGGATCTGGAAACGAGATCAAATCATCCAGTAAACCGCC |
| Prrx1_B2 | CCTCGTAAATCCTCATCAAATGAGCAGTTTACGGCTGTTTTGGCT |
| Prrx1_B2 | AGCCTTCCTGGAAAACCCTGGGCAAAAATCATCCAGTAAACCGCC |
| Prrx1_B2 | CCTCGTAAATCCTCATCAAAGTCCTGCTGCAGGTGCAATGGGAGG |

| | |
|---|---|
| Prrx1_B2 | TCTGCTATCCCTTGTCATGATAAGCAAATCATCCAGTAAACCGCC |

**Table S2.**

| Gene | Primer L | Primer R |
|---|---|---|
| Shh | GAGTCCAAGGCCCACATTCA | CTGGTCACTCCATGCTCCAG |
| Fgf8 | CCGCATTAAAGGAGCCGAGA | GGCCATGTACCATCCTTCGT |
| Grem1 | GAGCTAGCTGTTCCACCCAC | GCTCAGAGTCATTGGGCTGT |
| Hand2 | AGAGTGCAAGGAGGACAAGC | TCCTGCCCTTGCTCTTCTTG |
| Alx4 | CTTTTCAAGCGCCTACGAGC | GGAGACATGCAGGATGGGAC |
| Alx1 | ACTGCATCCCGAGAAACGAG | AGGGTGCAACAGTTCTCCAG |
| Ptch1 | TGAGCCGCCATGTACAAAGT | ACTGACACCAAGCAAACCCA |
| Gli3 | CGGCGGCAACCTAAATGATG | GTTCTGCAAGTGCTGTTGGG |
| Gli2 | CAGAAACACCAAGCTGCCAC | GTGTAGGCCGAGCTCATTGT |
| Gli1 | TGTACGAGACCAACTGCCAC | CAGCATGTACTGGGCCTTGA |
| Smo | CCTCACCTCCATTGCTCTGG | TTTCCGCTTTCGCACTTTGG |
| Bmp2 | ATGTTTCCAAGCGCACTTCG | CACTTGGTACAGGAGCAGCA |
| Bmp7 | CCCATGTTCATGCTGGACCT | GGGCTTGTAGGGGTAGGAGA |
| Pdgfra | AGAGATTGAGTGCCGACAGC | TCCTCGCGCTTCATGAAAGT |
| Cyp26b1 | AAAGAGCATGGCAAGGAGCT | TTGTGGAGGATACCGTTGCC |

**Table S3:**

| Cell Cycle State | Fluorescent Signature |
|---|---|
| G0 | pHH3-, mCherry-, mAG-, EdU- |
| G1 | pHH3-, mCherry+, mAG-, EdU- |
| S | pHH3-, mCherry-, mAG+, EdU+ |
| | pHH3-, mCherry+, mAG-, EdU+ |
| | pHH3-, mCherry-, mAG-, EdU+ |
| | pHH3-, mCherry+, mAG+, EdU+ |
| G2 | pHH3-, mCherry-, mAG+, EdU- |
| M | pHH3+, mCherry-, mAG+, EdU- |
| | pHH3+, mCherry+, mAG+, EdU- |
| | pHH3+, mCherry+, mAG-, EdU- |
